# Mechanistic Insights into Dengue Virus Inhibition by a Clinical Trial Compound NITD-688

**DOI:** 10.1101/2024.12.15.628546

**Authors:** Yan Wang, Long Sun, Luciana Fernandes, Yu-Hsiu Wang, Jing Zou, Samuel J. Franklin, Yanping Hu, Lee K. Palmer, Jason Yeung, Daniela Barriga, William K. Russell, Stephanie A. Moquin, Pei-Yong Shi, Colin Skepper, Xuping Xie

## Abstract

Dengue, caused by the dengue virus (DENV), presents a significant public health challenge with limited effective treatments. NITD-688 is a potent pan-serotype DENV inhibitor currently in Phase II clinical trials. However, its mechanism of action is not fully understood. Here, we present the molecular details of how NITD-688 inhibits DENV. NITD-688 binds directly to the nonstructural protein 4B (NS4B) with nanomolar affinities across all four DENV serotypes and specifically disrupts the interaction between NS4B and nonstructural protein 3 (NS3) without significantly changing the interactions between NS4B and other viral or host proteins. NS4B mutations that confer resistance to NITD-688 reduce both NITD-688 binding to NS4B and disruption of the NS4B/NS3 interaction. Specifically, NITD-688 blocks the interaction of NS3 with a cytosolic loop within NS4B. This inhibits the formation of new NS4B/NS3 complexes and disrupts pre-existing complexes in *vitro* and DENV-infected cells, ultimately inhibiting viral replication. Consistent with this mechanism, NITD-688 retains greater potency in cellular assays with delayed treatment compared to JNJ-1802, another NS4B inhibitor that has been studied in Phase II clinical trials. Together, these findings provide critical insights into the mechanism of action of NITD-688, facilitating the development of novel flavivirus NS4B inhibitors and informing future clinical interventions against DENV.

## Introduction

Dengue, caused by the mosquito-borne dengue virus (DENV), poses a significant threat to public health. There are four closely related but distinct serotypes of DENV, each capable of causing diseases ranging from mild flu-like illnesses to life-threatening dengue hemorrhagic fever (DHF) and dengue shock syndrome (DSS). Dengue has been reported in 176 countries^1^, extending beyond its traditionally endemic tropical and subtropical regions into previously unaffected areas. Annually, there are an estimated 390 million DENV infections globally, causing approximately 100 million symptomatic cases and 40,000 deaths^2,3^. Due to globalization, urbanization, and climate change, dengue continues to spread worldwide^4–7^. Half of the world’s population is now at risk of contracting dengue^8^.

Unfortunately, the currently available measures to stop dengue face limitations. DENV vaccine development has been challenged by the risk of antibody-dependent enhancement (ADE) of infection^9^. Dengvaxia (Sanofi-Pasteur), the only DENV vaccine approved and available in the United States and several DENV-endemic countries, is limited in use to people with laboratory-confirmed previous DENV infections^10^. Two DENV vaccine candidates with improved effectiveness, TAK-003 (Takeda) and TV003/TV005 (NIH) are in phase 3 clinical trials^11–14^. Neither has been approved by the United States Food and Drug Administration, although TV003 has been approved for use in some endemic and non-endemic countries. However, their ability to induce long-lasting balanced immune responses to all four serotypes remains unknown. Although vector controls have demonstrated potential in reducing dengue incidence^15,16^, their effectiveness typically relies on comprehensive community-wide approaches, which often encounter implementation challenges across areas with diverse cultures and economies. Despite decades of efforts, DENV therapeutics, including antibody and small molecule drugs, are not yet clinically available^17–21^, and thus dengue management remains supportive.

Direct-acting antivirals (DAAs) are molecules designed to inhibit infection by targeting viral proteins that are crucial for viral infection. They have demonstrated remarkable success in treating human immunodeficiency virus (HIV) and hepatitis C virus (HCV) infection^22,23^. However, developing DENV DAAs has proven to be challenging^24^. DENV belongs to the *Flavivirus* genus within the *Flaviviridae* family, which includes several other significant human pathogens, such as Japanese encephalitis virus (JEV), West Nile virus (WNV), Zika virus (ZIKV), and tick-borne encephalitis virus (TBEV). DENV is an enveloped virus with a single-stranded positive-sense RNA genome of ∼11,000 nucleotides in length. The viral genome contains a single open reading frame (ORF) flanked by 5′ and 3′ untranslated regions (UTRs). During viral infection, this ORF is translated into a large polyprotein that is further cleaved by both viral and host proteases into three structural proteins (capsid [C], precursor membrane [prM], and envelope [E]) and seven nonstructural proteins (NS1, NS2A, NS2B, NS3, NS4A, NS4B, and NS5). While the structural proteins form the viral particles, the nonstructural proteins play indispensable roles in viral RNA synthesis^25–27^, virion assembly^28,29^, and counteracting host innate immunity^30,31^. Among them, NS3 has serine protease (with the cofactor NS2B), RNA triphosphatase, and RNA helicase activities^32,33^. NS5 functions as a methyltransferase, guanylyltransferase, and RNA-dependent RNA polymerase^34–36^. These two multifunctional enzymes have been the primary antiviral targets over the years^18,37,38^. Unfortunately, most inhibitors targeting NS3 and NS5 have not yet progressed to clinical trials. Only AT-752 (Atea Pharmaceuticals), a guanosine nucleotide analog^39^, has reached phase 2 clinical trials (NCT05466240), though the development of the compound has since been halted.

In addition to NS3 and NS5, flavivirus NS4B has been another attractive antiviral target. NS4B is a small, structurally dynamic integral membrane protein with no known enzymatic activity^26,40^. It plays critical roles in viral infection by remodeling the endoplasmic reticulum membrane, counteracting host innate immunity, and inducing cell dysregulation^41^. At a molecular level, these functions are mediated through an interaction network involving NS4B, viral proteins (NS1, NS2B, NS3, and NS4A), as well as host factors^30,41–45^. Despite significant challenges in developing pan-genotype and pan-serotype small molecules with favorable safety profiles and drug-like physicochemical and pharmacokinetic properties, several DENV NS4B inhibitors have been uncovered in the past decade, including aminothiazole^46^, spirooxindolone^47^, SDM25N^48^, NITD-688^19^, JNJ-A07, and JNJ-1802^20,21^. Among them, NITD-688 and JNJ-1802 have excellent pan-serotype antiviral potencies in cell culture, as well as efficacy in animal models, and have both advanced to Phase 2 clinical trials (NCT06006559; NCT05201794)^49^.

A thorough understanding of the inhibitors’ antiviral mechanisms is essential for rational drug design and optimization to improve drug potency, selectivity, specificity, and safety. However, there is a considerable knowledge gap regarding how NS4B inhibitors perturb DENV infection, due to the lack of high-resolution tertiary structure of NS4B or robust assays. All reported NS4B inhibitors were identified through cell-based phenotypic screens, followed by target deconvolution via resistance selection-based genetic approaches, as exemplified by the discovery of the first DENV NS4B inhibitor, NITD-618^46^. Additional biochemical, biophysical, and virological assays are imperative to further delineate the underlying molecular basis of antiviral activity. In particular, the mode of action of two clinical front-runners, NITD-688 and JNJ-1802, has not been fully elucidated. NITD-688, exhibiting single to double-digit nanomolar potency against four serotypes of DENV, was found to bind directly to recombinant DENV NS4B proteins through nuclear magnetic resonance (NMR) spectroscopy^19^. JNJ-1802, exhibiting picomolar to low nanomolar efficacy against four serotypes of DENV, was shown to prevent viral replication complex formation by blocking NS4B/NS3 interactions in co-immunoprecipitation experiments^20,21^. Distinct NS4B mutations have been identified in viruses resistant to JNJ-1802 and a close analog of NITD-688. Given concerns about the emergence of resistant viruses during treatment, it is crucial to know whether NITD-688-resistant viruses are cross-resistant to JNJ-1802 and vice versa. Furthermore, understanding additional molecular details about the mechanism of action of these two compounds could be useful for developing next-generation NS4B inhibitors.

In this study, we have examined the antiviral mechanism of NITD-688 using a multipronged approach, including viral genetics, biophysical, biochemical, imaging, and virological methods. We find that NITD-688 binds directly to DENV NS4B with nanomolar affinities and is capable of disrupting preformed NS4B/NS3 complexes *in vitro* and in DENV-infected cells. Consistent with this mechanism, NITD-688 exhibits less reduction in potency in cellular assays with delayed treatment compared to JNJ-1802, another clinically studied NS4B inhibitor. Our findings can provide new avenues for developing NS4B-targeted flavivirus inhibitors and future clinical treatment options against dengue.

## Results

### Characterization of NITD-688-resistant viruses

The tetrahydrothienopyridine NITD-688 was reported as a DENV NS4B inhibitor with nanomolar potencies against all four serotypes of DENV^19^. Previously, resistance selection was performed with a close analog of NITD-688. To understand the specific susceptibility of NITD-688 to the development of resistance, we performed serial passages of the DENV-2 NGC strain in the presence of increasing concentrations of NITD-688 (Fig. 1a). Following 15 passages (approximately 50 days), all seven independent NITD-688 selections (S1-S7) yielded variants resistant to the inhibitor, with 240.8- to 766.5-fold increase in EC_50_ values when compared to the un-passaged wild-type (WT) virus (Fig. 1b). In contrast, four independent DMSO selections (S8-S11) remained highly sensitive to NITD-688, similar to the parental WT virus. Whole-genome sequencing of the P15 viruses demonstrated NS4B as a hotspot for mutations during DENV passaging in cell cultures (Fig. S1 & Table S1). A total of thirteen amino acid substitutions in NS4B were identified. Ten of them (R33C/Y/S/H, V175A, A193V, T195A, W205L, T215A, and A222V) were present only in NITD-688 selections but not in DMSO selections, or only present in DMSO selections at very low frequencies (<4%). Most prevalent was A222V, present in all seven NITD-688 selections at >99% frequency. Next, A193V was present in all seven NITD-688 selections with varied frequencies: high frequency (>99%) in two selections, modest frequency (9.5-24.4%) in another two selections, and low frequency (<4%) in the rest of the selections. T195A was present in four selections, with three having >97% frequency. T215A and W205L each predominated in one of the seven selections. R33C/Y/S/H were present in four selections with <37% frequencies. While L111F and S238F were present in some NITD-688 selections at >90% frequency, they also emerged in DMSO selections at high frequencies, suggesting they may emerge in response to passaging in cell cultures but not drug pressure. V175A was present in only one NITD-688 selection at 21% frequency. T179I was only observed in the DMSO passaged virus. Mutations were also observed in prM, E, NS1, NS2A, NS2B, NS3, NS4A, and NS5. However, the mutations in proteins other than NS4B were either not unique to the NITD-688 selection or were inconsistent across the seven NITD-688 selections, and, therefore, were not further investigated in this study.

**Fig 1.**
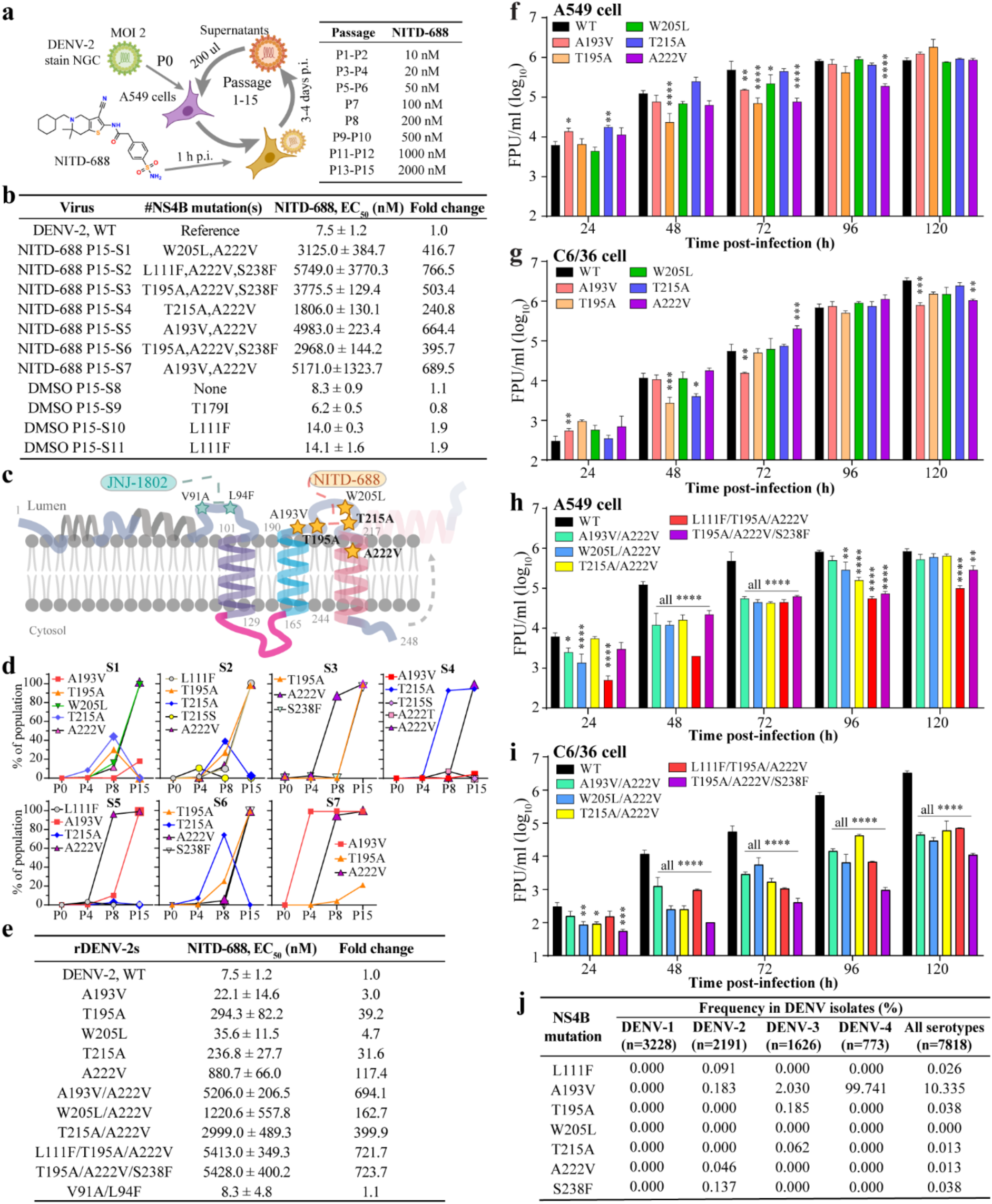
Selection and characterization of NITD-688 resistant DENV. **a** Scheme for the process of selecting NITD-688-resistant viruses in cell culture. See details in Materials and Methods. **b** EC_50_ values of NITD-688 against P15 viruses. The means ± SD were derived from two independent experiments for P15 viruses and six independent experiments for the reference WT DENV-2, each performed with three technical replicates. Fold change was calculated by normalizing the EC_50_ of each mutant to that of WT. ^#^the consensus amino acid changes (with >90% frequencies) within NS4B. **c** Membrane topology of NS4B with identified mutants. Mutations unique in NITD-688-selected P15 viruses are colored in orange. V91A and L94F, conferring major resistance to JNJ-1802 are shown in blue. **d** Emergence of NS4B mutations in P4, P8, and P15 of NITD-688-selected viruses. Each amino acid mutation is shown in the same color across selections (S1-S7). **e** EC_50_ values of NITD-688 against recombinant DENV-2 containing NS4B mutation(s). Data are presented as mean±SD from two independent experiments. **f** Viral growth kinetics of single mutant viruses on A549 cells. **g** Viral growth kinetics of single mutant viruses on C6/36 cells. **h** Viral growth kinetics of double and triple mutant viruses on A549 cells. **i** Viral growth kinetics of double and triple mutant viruses on C6/36 cells. **f**-**i**, means and standard deviations from 2-3 independent experiments are shown. Two-way ANOVA with Dunnett’s multiple comparison test was used to perform the group comparison. *p<0.05; **p<0.01; ***p<0.001; ****p<0.0001. **j** Occurrence of NITD-688-resistant NS4B mutations in the general population of DENVs. n, numbers of DENV genome sequences used for analysis.

To examine the dynamics of NS4B mutations over serial passages, the NS4B sequences of passages 4 (P4), 8 (P8), and 15 (P15) viruses under NITD-688 selection were determined using PacBio sequencing. Seven NS4B mutations prevalent in P15 viruses were analyzed in-depth (Fig. 1d). A222V emerged in P4 and dominated all selections by P15. T195A occurred in five out of seven selections in P8 and dominated in three selections by P15. A193V appeared as early as P4, prevalent in two selections by P15, and appeared in two additional selections at lower frequencies. Notably, T215A/S was initially detected in two selections in P4 and subsequently expanded to four selections by P8. However, T215A predominated only in one selection by P15. W205L emerged independently from T195A, T215A, and A222V, and later co-prevailed with A222V in one selection. L111F and S238F co-evolved with T195A and A222V and predominated in one to two selections by P15. Furthermore, the PacBio sequencing results unveiled a progression in mutation complexity, transitioning from a single mutation to double or even triple mutations within the quasispecies (Fig. S2). The positive selection of A222V in combination with other mutations is associated with heightened resistance in response to the increasing concentrations of NITD-688. The identified NS4B residues have varied conservation levels among the flavivirus genus (Fig. S3). W205 is conserved among four DENV serotypes, ZIKV, JEV, and WNV; T195, T215A, and A222 are only conserved among four DENV serotypes, while A193 is conserved in DENV-2, DENV-3, JEV, WNV, TBEV, and ZIKV.

To assess the contribution of each amino acid substitution to drug resistance and viral fitness, we generated a panel of DENV-2 mutants in the infectious clone of the NGC strain^29^, including five single mutants (A193V, T195A, W205L, T215A, and A222V), three double mutants (A193V/A222V, T215A/A222V, and W205L/A222V), and two triple mutants (L111F/T195A/A222V and T195A/A222V/S238F). The double and triple mutations were chosen for their prevalence in NITD-688 selections at P15 (Fig. 1d, S1, S2). All mutant viruses were successfully rescued and generated plaques with sizes comparable to WT DENV-2 (Fig. S4). The sensitivity of mutant viruses to NITD-688 was determined by a High Content Imaging Cellular Flavivirus Immunoassay (HCI-CFI) in A549 cells (Fig. 1e). Among single mutants, A222V conferred the most significant resistance to NITD-688, with EC_50_ values increased by 117.4-fold when compared to WT virus. Viruses with T195A and T215A displayed a 39.2-fold and a 31.6-fold increase in EC_50_, respectively. Viruses with A193V and W205L showed only a modest increase in EC_50_ values of 3.0- and 4.7-fold, respectively (Fig. 1e). The addition of either A193V or T215A to viruses containing A222V resulted in a further increase in EC_50_: from a 117.4-fold increase in EC_50_ when compared to WT virus for the A222V virus alone, to either 694.1-fold or 399.9-fold increase in EC_50_ for the double mutants A193V/A222V or T215A/A222V, respectively. In contrast, the addition of W205L to the A222V mutant virus did not drastically increase the EC_50_. Triple mutants L111F/T195A/A222V and T195A/A222V/S238F conferred the most resistance to NITD-688, with EC_50_ values shifted by over 721.7-fold. Notably, double or triple mutants reached EC_50_ fold shifts comparable to the P15 population of selected viruses (Fig. 1b, e), suggesting NS4B mutations are sufficient to confer complete resistance to NITD-688.

**Figure 2.**
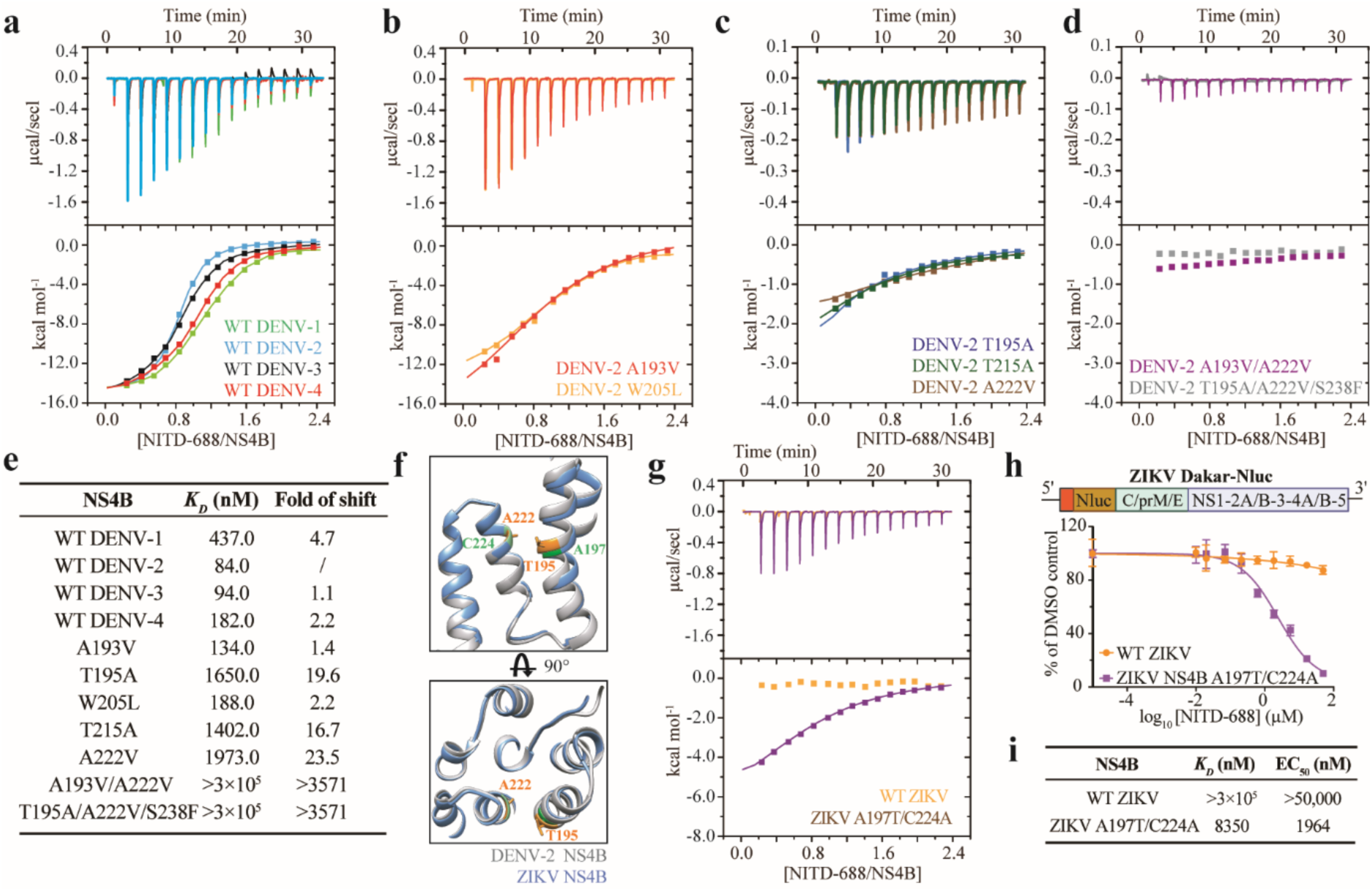
NITD-688 directly binds to DENV NS4B. **a** ITC analysis of NITD-688 binding to NS4B across DENV four serotypes. The top panel shows the raw ITC data. The bottom panel shows the fitting curves (binding model 1:1). **b** ITC analysis of NITD-688 binding to DENV-2 NS4B mutants A193V or W205L. **c** ITC analysis of NITD-688 binding to DENV-2 NS4B mutants T195A, T215A, or A222V. **d** ITC analysis of NITD-688 binding to DENV-2 NS4B mutants A193V/A222V or T195A/A222V/S238F. **e** Summary of *K*_D_ values estimated in panels a-d. All mutations were made in the context of DENV-2 NS4B. **f** Alignment of the Alphafold2-predicted structures of DENV-2 and ZIKV NS4B regions at amino acid positions 180 to 240. Residues A222 and T195 in DENV-2 NS4B are labeled in orange, while residues A197 and C224 in ZIKV NS4B are colored in green. **g** ITC analysis of NITD-688 binding to wildtype (WT) ZIKV NS4B or mutant A197T/C224A. **h** Antiviral activity of NITD-688 against WT ZIKV-Nluc or A197T/C224A mutant in A549 cells at 48 h post-infection. Mutant A197T/C224A was engineered into the backbone of Dakar strain-derived reporter ZIKV containing a Nanoluciferase gene, ZIKV Dakar-Nluc^75^. **i** Summary of binding *K*_D_ and EC_50_ values of NITD-688 against WT ZIKV or mutant A197T/C224A.

We then examined the growth kinetics of each recombinant virus in mammalian and mosquito cells to assess the effect of NS4B mutations on viral fitness. Among single mutants, T195A and A222V consistently showed delayed growth in human A549 cells between 48 and 96 h post-infection (Fig. 1f). However, none of the single mutants consistently reduced viral growth in mosquito C6/36 cells (Fig. 1g). In contrast, all three double mutants exhibited significantly slower and delayed growth in A549 cells and even greater attenuation in C6/36 cells (Fig. 1h-i). Both triple mutants exhibited the most attenuation in A549 and C6/36 cells (Fig. 1h-i).

To determine the prevalence of residues that emerged in the resistance selection, we examined 7818 sequences of the DENV genome available, as of 11 Dec 2024, in the Bacterial and Viral Bioinformatics Resource Center (BV-BRC)^50^. Overall, these residues show low prevalence in known dengue sequences (<0.038%), except for V193, which was observed in 10.335% of dengue sequences (Fig. 1j). Among the residues that showed the largest shifts in potency (A195, A215, V222), A195, A215, and V222 appeared in three DENV-3 strains, one DENV-3 strain, and one DENV-2 strain, respectively. None of the strains contained any combinations of three mutations. Although V193 was found in all DENV-4 strains, it had low frequencies in DENV-2 and DENV-3. This may account for the lower sensitivity of tested DENV-4 strains to NITD-688 than DENV-2 and DENV-3 strains^19^. Collectively, our data indicate that NS4B mutations that confer the strongest resistance to NITD-688 are associated with viral fitness costs, occur infrequently in DENV databases, and may not be highly transmissible.

### NITD-688 inhibits JNJ-1802-resistant DENV variants, and vice versa

It was previously reported that NS4B mutations T195A and A222S transiently emerged during the selection of JNJ-1802-resistant viruses^20^. Notably, A222S exhibited no shift in susceptibility and T195A exhibited only a small decrease in susceptibility to JNJ-1802. The same study revealed two NS4B mutations, V91A and L94F, which conferred strong resistance to JNJ-1802. We then asked whether viruses resistant to NITD-688 exhibit cross-resistance to JNJ-1802 and vice versa. We constructed an additional double mutant virus, V91A/L94F (Fig. S4a). This mutant, along with two NITD-688-resistant variants, L111F/T195A/A222V, and T195A/A222V/S238F, were assessed in parallel for their sensitivity to NITD-688 and JNJ-1802. Consistent with previous findings^20^, V91A/L94F showed 2941.3-fold resistance to JNJ-1802 (Table S2), while it remained sensitive to NITD-688, similar to the WT DENV-2 (Fig. 1e). On the other hand, JNJ-1802 maintained potency against NITD-688 resistant strains L111F/T195A/A222V and T195A/A222V/S238F (Table S2). These results demonstrated marginal cross-resistance between variants resistant to NITD-688 and JNJ-1802, supporting that these two NS4B inhibitors target distinct regions within DENV NS4B.

### NITD-688 directly binds to DENV NS4B from all four DENV serotypes

We employed isothermal titration calorimetry (ITC), a label-free biophysical approach, to assess the direct binding between NITD-688 and mature DENV NS4B. ITC allows measuring the binding affinity between two or more molecules in solution^51^. Recombinant NS4B proteins with an N-terminal hexahistidine tag were purified and reconstituted in detergent micelles (Fig. S5a). Size exclusion chromatography (SEC) and SDS-PAGE analysis indicated the high purity of recombinant NS4B proteins (Fig. S5b, c). NITD-688 demonstrated binding affinities to four serotypes of DENV NS4B ranging from 84-437 nM, with affinity ranking from high to low: DENV-2∼DENV3>DENV-4>DENV-1 (Fig. 2a&e). Notably, a similar trend was observed for activity in cellular assays^19^, with NITD-688 being similarly potent against DENV-2 and DENV-3, less potent against DENV-4, and least potent against DENV-1.

To assess the impact of the identified NS4B mutations on NITD-688 binding, we purified a panel of DENV-2 NS4B mutant proteins (Fig. S5c). This panel included five single mutants (A193V, T195A, W205L, T215A, and A222V), one double mutant (A193V/A222V), and one triple mutant (T195A/A222V/S238F). The single mutations were chosen because of their presence only in NITD-688 selections but not in DMSO selections (Table S1), while the double and triple mutants were selected because they demonstrated the largest increases in EC_50_ values in cellular assays (Fig. 1e). Mutations A193V or W205L slightly reduced the NITD-688 binding to NS4B (Fig. 2b,e). In contrast, T195A, T215A, and A222V reduced the binding by 16.7- to 23.5-fold (Fig. 2c,e). Both A193V/A222V and T195A/A222V/S238F mutants showed no binding at up to 300 µM of NITD-688 (Fig. 2d,e). These data suggest that NS4B residues T195, T215, and A222 comprise a putative binding site for NITD-688, and that loss of binding to NS4B is the mechanism of resistance.

### ZIKV becomes susceptible to NITD-688 after the introduction of residues critical for NITD-688 binding

Next, we sought to determine whether altering these key residues in NS4B could confer NITD-688 sensitivity to other flaviviruses. NITD-688 displays broad activity against the four DENV serotypes, but not against any other tested flaviviruses, likely because the key residues T195, T215, and A222 are not conserved among other flaviviruses (Fig. S3). The tertiary structure of flavivirus NS4B is currently unavailable; however, AlphaFold2 structural predictions show residues T195 and A222 near neighboring alpha helices, with an estimated C_α_ distance of 7.3 Å (Fig. 2f). Additionally, Alphafold2 predicted high structural homology between DENV-2 and ZIKV NS4B (Fig. S6a). Thus, we engineered the DENV NS4B residues T195 and A222 into ZIKV (corresponding to A197T and C224A) within the context of recombinant NS4B protein as well as the full-length viral genome. First, we expressed and purified recombinant WT and mutant (A197T/C224A) ZIKV NS4B proteins (Fig. S6b). ITC showed no binding of NITD-688 to WT ZIKV NS4B protein at tested concentrations up to 300 µM (Fig. 2g, i). In contrast, NITD-688 bound NS4B mutant A197T/C224A with an affinity of 8.4 μM. Next, we tested whether these mutations could render ZIKV susceptible to NITD-688. We engineered the A197T/C224A mutations into a ZIKV Dakar-Nluc backbone and recovered the recombinant mutant virus. ZIKV mutant A197T/C224A developed WT-like plaque morphologies (Fig. S6c). While NITD-688 did not inhibit WT ZIKV at concentrations up to 50 μM, it was able to effectively inhibit ZIKV Dakar-Nluc mutant A197T/C224A with an EC_50_ of 1.96 µM (Fig. 2h-i).

### NITD-688 primarily inhibits NS4B/NS3 interaction

NS4B plays a crucial role in viral infection through interacting with both viral and host factors^44,45,52,53^. To assess the impact of NITD-688 on the interactome of DENV NS4B, we conducted co-immunoprecipitation (co-IP) assays using co-expression of NS4B and other viral proteins in HEK-293T cells. Initially, co-IPs confirmed the specific interactions between NS4B and NS4B itself, NS4A, NS1, NS2A, and NS3 (Fig. 3a-e). We observed comparable levels of NS4B, NS4A, NS1, and NS2A co-immunoprecipitated by NS4B from cells treated with NITD-688 at concentrations up to 100 nM compared to those from DMSO-treated cells (Fig. 3a-d). Notably, NS4A pulled down more 2K-NS4B than mature NS4B (Fig. 3b), suggesting that the 2K sequence may interact with NS4A. However, NITD-688 did not inhibit this interaction. In contrast, the amount of NS3 that immunoprecipitated with NS4B was significantly reduced by NITD-688 treatment, even at the lowest concentration of 5 nM (Fig. 3e). Interestingly, NITD-688 was not able to disrupt the interaction between NS3 and the NS4B triple mutant (TM) T195A/A222V/S238F, as the same amount of NS3 was immunoprecipitated with TM NS4B even at 100 nM of NITD-688 (Fig. 3f).

**Figure 3.**
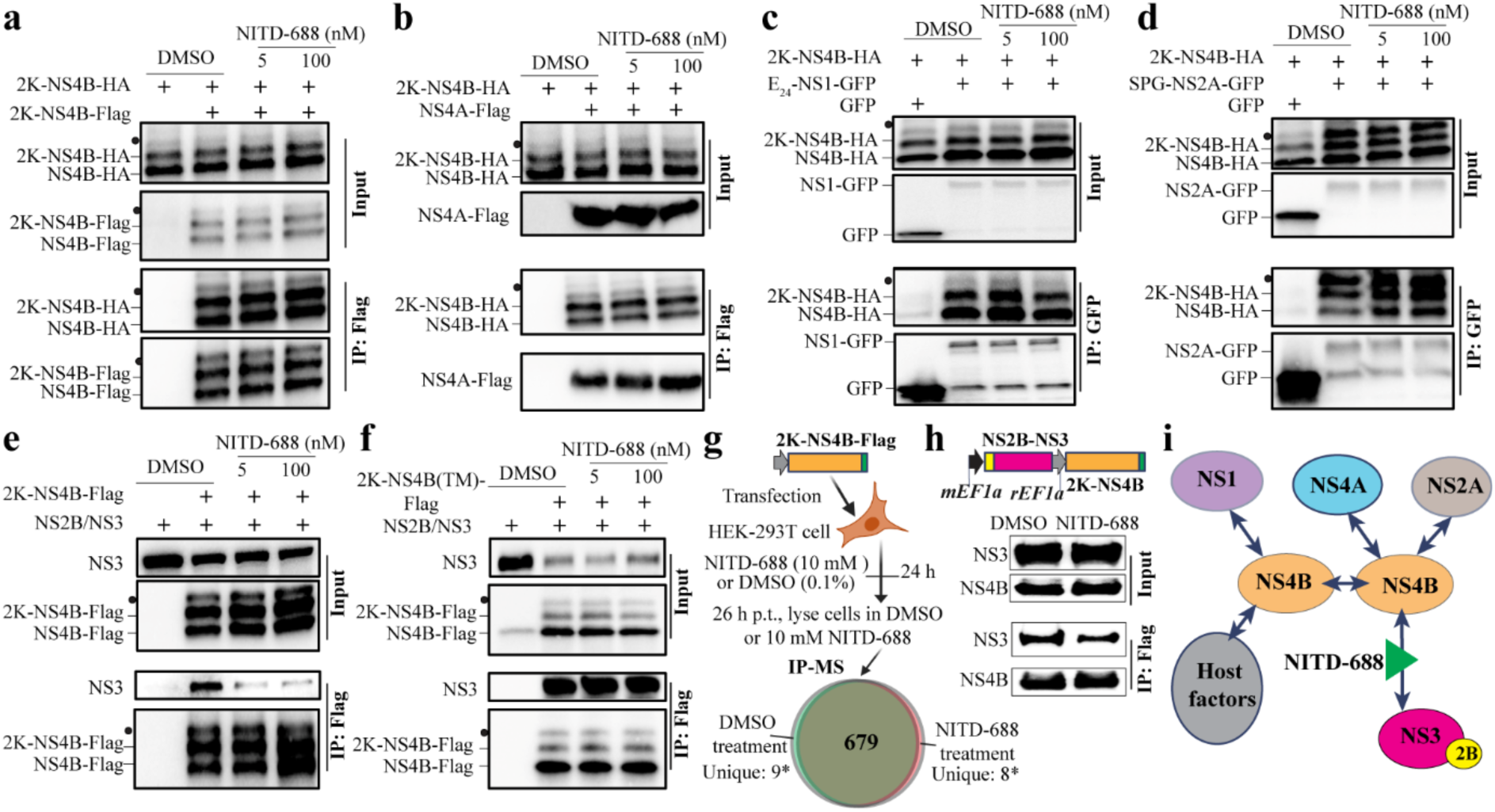
NITD-688’s impact on the interactome of NS4B. HEK-293T cells were transfected with two plasmids: one encoding 2K-NS4B-HA and the other encoding either 2K-NS4B-Flag (**a**), NS4A-Flag (**b**), E_24_-NS1-GFP^62^ (**c**), SPG-NS2A-GFP^29^ (**d**), or NS2B/NS3 (**e**). Modified NS4B (indicated as solid circles), uncleaved 2K-NS4B, and mature NS4B co-existed in transfected cells. As controls, NS2B/NS3 co-immunoprecipitated by an NS4B triple mutant (T195A/A222V/S238F, “TM”) was also tested (**f**). At 4 h post-transfection, cells were exposed to DMSO or inhibitors. At 20 h post-treatment, cells were lysed and subjected to immunoprecipitation followed by Western blot analysis. An empty vector (in panels **a**, **b**, **e**, and **f**) or a plasmid expressing GFP alone (in panels **c** and **d**) was used for negative controls. **g** Diagram of IP-MS process for assessing NITD-688’s impact on NS4B-host factor interaction. A Venn diagram reveals a total of 679 proteins overlapped in both DMSO and NITD-688-treated groups, nine proteins only present in the DMSO-treated group, and eight proteins only in the NITD-688-treated group. *detection with low confidence (1 or 2 of the three IP-MS replicates). See details in Table S4. **h** NITD-688 reduces NS4B/NS3 interactions. HEK-293T cells were transfected with one plasmid with a dual promoter: one expressing 2K-NS4B-Flag and the other expressing NS2B/NS3. Cells were treated with DMSO or NITD-688 (10 µM) at 24 h post-transfection. After two hours of treatment, cells were lysed and subjected to co-immunoprecipitation. **i** Diagram of NITD-688’s impact on the interactome of NS4B.

Next, we used an unbiased method, immunoprecipitation mass spectrometry (IP-MS), to evaluate the impact of NITD-688 on the interaction of NS4B with host proteins (Fig. 3g). Similar host proteins immunoprecipitated with NS4B in cells treated with NITD-688 compared to those from DMSO-treated cells. Specifically, 679 host proteins were immunoprecipitated under both DMSO or NITD-688 treatment, with no significant difference in protein abundance in NITD-688- or DMSO-treated cells.

Nine host proteins (MYH9, PPIAL4A, PIGS, AMOTL1, ACTB, TUBAL3, HNRNPF, KRT74, and IGF2BP3) were present only in DMSO-treated samples, while eight host proteins (TUBB2A, TUBB3, KNRNPDL, RBBP4, RANBP3, ANKRD35, SMC3, and HNRNPH3) were found only in NITD-688-treated samples. However, none of these proteins were consistent in all three replicates (Table S4). As a control, under the same experimental conditions, NITD-688 significantly reduced NS3 co-immunoprecipitated by NS4B (Fig. 3h). Like previous reports^53^, our IP-MS analysis reaffirmed the critical role of NS4B in multiple cellular pathways, including protein processing in the endoplasmic reticulum, regulation of protein stability, and antiviral mechanisms by IFN-stimulated genes (Fig. S7). Overall, these findings indicate that NITD-688 primarily blocks the NS4B/NS3 interaction without significantly affecting NS4B dimerization, or its interactions with NS1, NS2A, NS4A, or host factors (Fig. 3i).

### NITD-688 inhibits NS4B/NS3 interaction *in vitro*

We next sought to address how NITD-688 inhibits the NS4B/NS3 interaction at the molecular level. First, we conducted ITC to determine the affinity of the NS4B/NS3 interaction in the presence or absence of inhibitors using the purified recombinant DENV-2 NS4B proteins and full-length NS3 (Fig. S8a). In the absence of inhibitors, NS4B bound NS3 with an affinity of 12.3 μM (Fig. 4a,c). The cytosolic loop of NS4B (amino acids 121-171) bound NS3 with an affinity similar to the full-length NS4B (Fig. S8b). Consistent with our previous data obtained through surface plasmon resonance (SPR)^44^, this result further validates that the cytosolic loop of NS4B is responsible for binding to NS3.

**Figure 4.**
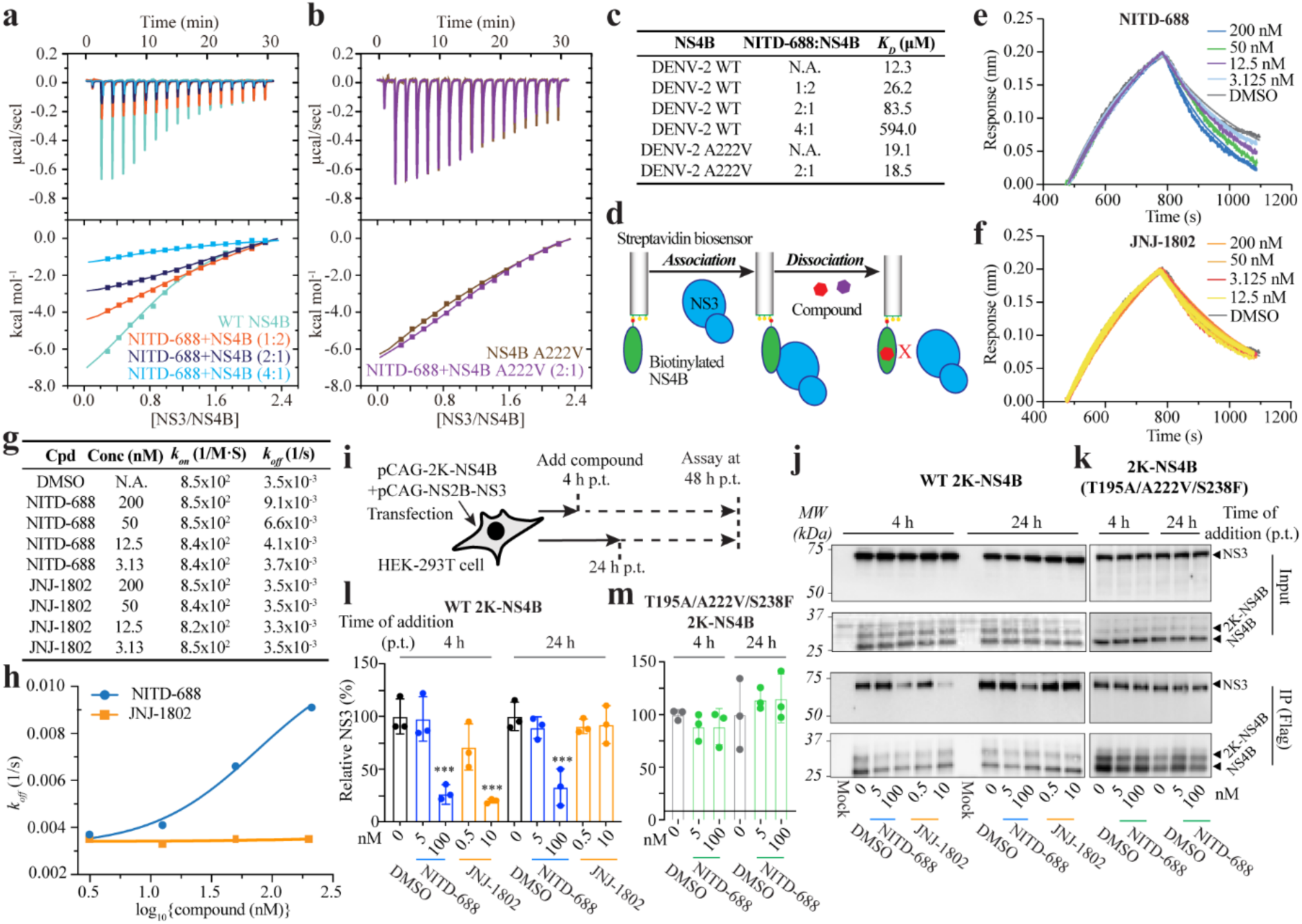
NITD-688 disrupts the NS4B/NS3 interaction. **a** ITC analysis of DENV-2 NS4B binding to NS3 in the presence or absence of NITD-688. DENV-2 NS4B was pre-incubated with indicated molar ratios of NITD-688 or 0.5 % DMSO. **b** ITC analysis of NITD-688’s impact on NS4B mutant A222V binding to NS3. **c** Summary of NS4B/NS3 binding *K*_D_ values estimated from panels a-b. **d** Scheme for BLI analysis of inhibitors’ impact on the dissociation of NS3/NS4B complex. Streptavidin (SA) biosensors were pre-soaked in biotinylated NS4B and followed by incubation with NS3 to form the NS3/NS4B complexes. The dissociation signals of NS3 from preformed NS4B/NS3 complexes were measured in 0.5% DMSO buffer or inhibitors. See details in Materials and Methods. **e** BLI curves show the NITD-688’s impact on the dissociation of NS3/NS4B complexes. **f** BLI curves show the JNJ-1802’s impact on the dissociation of NS3/NS4B complexes. **g** Summary of *k_on_* and *k_off_* values calculated from panels e&f. **h** Dose-*k_off_* curves. NITD-688 and JNJ-1802 were colored in blue and orange, respectively. **i** Scheme of the co-IP experiments for analyzing NS4B/NS3 interaction in cells. Plasmids encoding NS2B-NS3 and 2K-NS4B with a C-terminal Flag tag (2K-NS4B-Flag) were co-transfected into HEK-293T cells. Cells were exposed to inhibitors at 4 h (early treatment) or 24 h (delayed treatment) post-transfection. At 48 h post-transfection, cells were harvested for immunoprecipitation followed by Western blot analysis. **j** Western blot analysis of NS3 co-IPed with WT NS4B-Flag. Representative plots from three independent pull-down experiments. **k** Western blot analysis of NS3 co-IPed with NS4B mutant T195A/A222V/S238F. Representative plots from three independent Co-IP experiments. **l** Quantification of NS3 co-IPed by WT NS4B. The NS3 co-IPed with NS4B from DMSO-treated cells was set at 1.0. **m** Quantification of NS3 co-IPed with NS4B mutant T195A/A222V/S238F. The NS3 co-IPed with NS4B from DMSO-treated cells was set at 1.0. (**l-m**) Means and standard deviations from three independent experiments are shown. One-way ANOVA with Dunnett’s multiple comparison correction was used for group comparison between each treated group and the DSMO-treated group. ***p<0.001.

We then measured NS4B/NS3 binding in the presence of inhibitors using ITC. NS4B was pre-incubated with inhibitors before binding to NS3. We observed reduced NS4B/NS3 binding as the concentration of NITD-688 increased (Fig. 4a). Specifically, NS4B/NS3 binding affinities decreased from 26.2 to 594.0 µM as the molar ratio of NITD-688 to NS4B increased from 1:2 to 4:1 (Fig. 4c). Moreover, NITD-688 was unable to reduce binding of NS3 to NS4B A222V (Fig. 4b, c), consistent with the decreased binding affinity of NITD-688 to NS4B A222V (Fig. 2e). Notably, A222V slightly reduced the binding affinity of NS4B to NS3 from 12.3 µM to 19.1 µM, which may explain its reduced viral fitness in cell cultures. ITC also confirmed that NITD-688 did not bind to NS3 or the cytosolic loop of NS4B alone (Fig. S8c, d). This indicates that NITD-688 does not disrupt the interaction between NS4B and NS3 by directly binding to the cytosolic loop of NS4B (aa 121-171); rather, NITD-688 likely binds to NS4B near the residue A222 and disrupts the interaction with NS3 through an indirect mechanism.

Like NITD-688, JNJ-1802 also bound the full-length NS4B but not the cytosolic loop of NS4B (Fig. S8d-e). In addition, JNJ-1802 effectively inhibited the NS4B/NS3 binding in a dose-dependent manner (Fig. S8f). Notably, compared to NITD-688, JNJ-1802 has a 4.2-fold higher binding affinity to NS4B (20 nM for JNJ-1802 vs. 84 nM for NITD-688) (Fig. 2e, S8e), and correspondingly a 3.6- and 3.0-fold greater inhibition of the NS4B/NS3 binding at the molar ratios of inhibitors to NS4B of 1:2 and 2:1, respectively (Fig. 4c versus Fig. S8f). This higher binding affinity and increased ability to dissociate NS3 from NS4B may contribute to the greater potency of JNJ-1802 in standard cellular assays when compared to NITD-688 (0.019 nM for JNJ-1802 vs. 7.5 nM for NITD-688) (Fig.1e, Table S2). In summary, our data demonstrated that both NITD-688 and JNJ-1802 can specifically bind to DENV NS4B and inhibit the *de novo* formation of NS4B/NS3 complexes.

We assessed the effects of these two NS4B inhibitors on preformed NS4B/NS3 complexes using Biolayer Interferometry (BLI). BLI allows for the quantification of binding strength and kinetics, including association and dissociation rate constants^54^. The NS4B/NS3 complexes were assembled by incubating streptavidin biosensors pre-loaded with biotinylated recombinant DENV-2 NS4B in an NS3 protein solution. Subsequently, the NS4B/NS3 complex-captured biosensors were immersed into buffers containing DMSO or inhibitors, and the dissociation of NS3 from the NS4B/NS3 complexes was monitored (Fig. 4d). In the absence of inhibitors, NS3 bound NS4B with an affinity of 13 µM (Fig. S9a), consistent with the ITC data. The dissociation of NS3 from the NS4B/NS3 complexes was significantly accelerated by NITD-688 but not by JNJ-1802 (Fig. 4e-h). Specifically, the calculated dissociation rate constant (*k*_off_) of NS4B/NS3 complexes increased by 2.6-fold when the concentration of NITD-688 increased from 3.1 to 200 nM, while it remained unchanged in the presence of JNJ-1802 at concentrations up to 200 nM (Fig. 4g, h).

Taken together, our data indicates that both NITD-688 and JNJ-1802 can prevent the *de novo* formation of NS4B/NS3 complexes. However, only NITD-688 can disrupt preformed NS4B/NS3 complexes.

### NITD-688 disrupts preformed NS4B/NS3 complexes in cells

It is expected that more NS4B/NS3 complexes would form as viral protein levels increase over time. Therefore, we revisited the co-IP assays and examined the amount of NS3 that co-IPed with NS4B when inhibitors were added at 4 h (representing early treatment to prevent *de novo* formation of NS4B/NS3 complexes) and at 24 h (representing delayed treatment to disrupt preformed NS4B/NS3 complexes) post-transfection (Fig. 4i). A concentration of 10 nM of JNJ-1802 (526-fold the EC_50_ against DENV-2, 0.019 nM) decreased the amount of NS3 that co-IPed with NS4B by 80% when added at 4 h post-transfection (Fig. 4j-l), but only by 8% when added at 24 h post-transfection. This result aligned with previous reports that JNJ-1802 only prevents the *de novo* formation of NS3/NS4B complexes^20^. In contrast, NITD-688 significantly reduced the amount of NS3 that immunoprecipitated with NS4B regardless of the time of addition of the inhibitor. Specifically, in the presence of 100 nM of NITD-688 (13-fold the EC_50_ against DENV-2, 7.5 nM), the amount of NS3 that co-IPed with NS4B was decreased by 73% and 67% when the inhibitor was added at 4 or 24 h post-transfection, respectively. Importantly, NITD-688 only marginally reduced the amount of NS3 that immunoprecipitated with the NS4B mutant T195A/A222V/S238F (Fig. 4k-m). These findings suggest that NITD-688 can disrupt both the formation of NS4B/NS3 complexes and those already established in cells, even during delayed treatment.

### NITD-688 disrupts NS4B/NS3 complexes in infected cells

To evaluate whether NITD-688 could disrupt NS4B/NS3 complexes during viral infection, we developed an immunostaining assay to probe NS4B proteins in DENV-infected cells (Fig. 5). This assay utilized two monoclonal antibodies (mAbs), 10-3-7 and 44-4-7, which recognize distinctive epitopes mapped to residues 5-15 and 141-147 of NS4B, respectively^55^. The epitope targeted by mAb 10-3-7 is situated at the N-terminus of NS4B on the ER lumen side, whereas the epitope of mAb 44-4-7 overlaps with the cytosolic loop (121-171) where NS3 binds^44^. Using BLI, we initially confirmed that both mAbs specifically bound to the full-length NS4B proteins with double digital nanomolar affinities (Fig. S9b-c). Furthermore, BLI revealed that NS3 bound to the NS4B-mAb 10-3-7 complex, suggesting that non-binding sites for NS3 and mAb 10-3-7 on NS4B do not overlap. In contrast, NS3 increased the dissociation of the NS4B-mAb 44-4-7 complex (Fig. S9d), indicating that NS3 competes with mAb 44-4-7, but not with mAb 10-3-7, in binding to NS4B.

**Figure 5.**
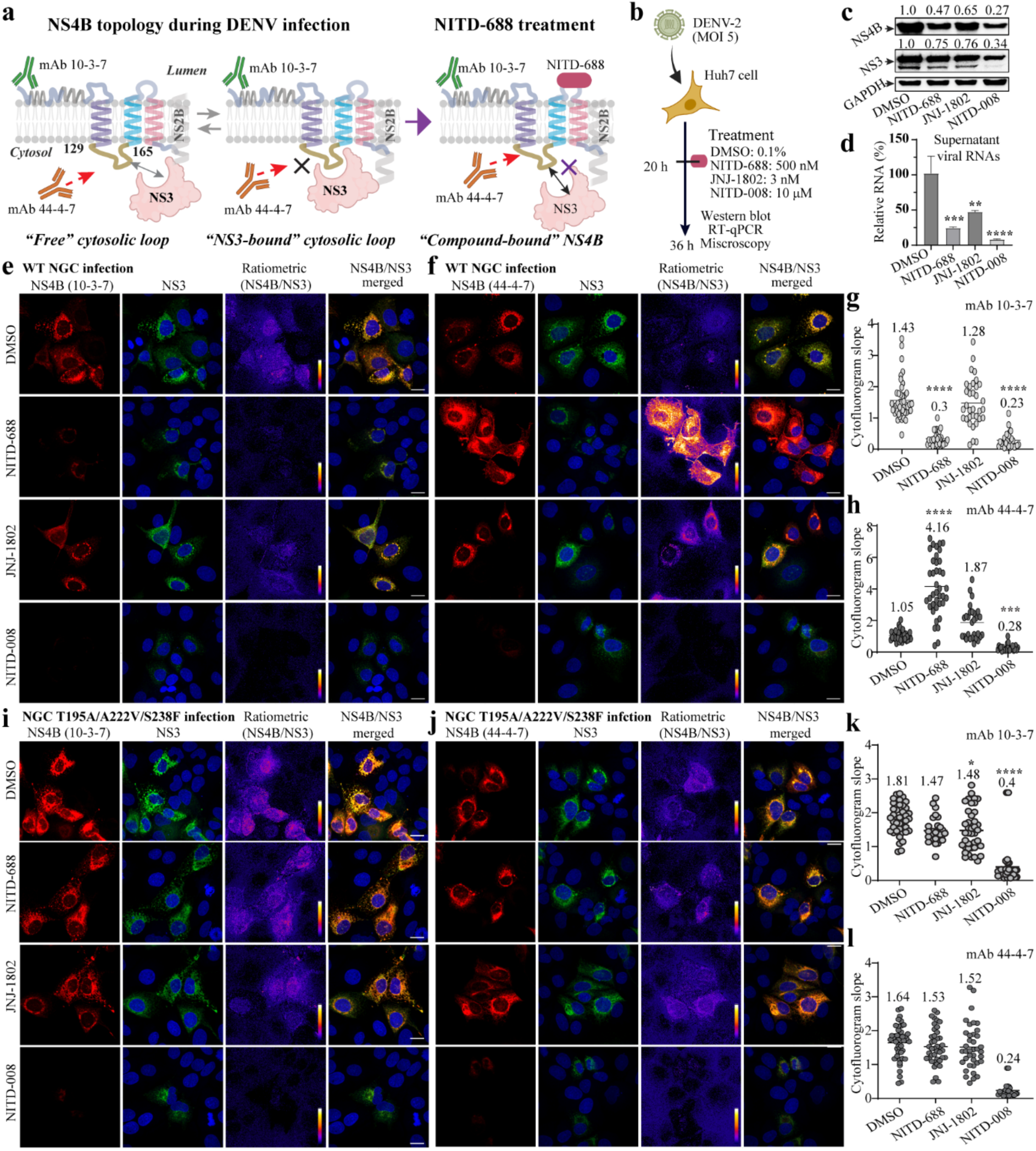
NITD-688 disrupts NS4B/NS3 complexes in DENV-infected cells. **a** Cartoon of NTID-688’s impact on NS4B/NS3 complexes in DENV-infected cells. **b** Scheme of DENV-2 infection with delayed treatment. **c** Intracellular viral protein expression after DENV-2 infection. Numbers indicate the relative abundance of proteins. The protein level in DMSO-treated cells was set at 1.0. **d** Extracellular viral RNA levels. The RNA level in DMSO-treated cells was set at 100%. Error bars indicate the standard deviations from three independent experiments. **e-f** Representative images of NS4B and NS3 staining in WT DENV-2-infected cells when NS4B was probed by mAb 10-3-7 (**e**) or 44-4-7 (**f**). Scar bar, 20 μm. The fluorescence ratiometric color scale ranges from 0 (dark purple) to 9.0 (bright yellow), representing the increase in fluorescence ratios of NS4B over NS3. **g-h** Cytofluorogram analysis of NS3 and NS4B within WT DENV-2-infected cells. NS4B was probed with mAb 10-3-7 (**g**) or 44-4-7 (**h**). Each dot represents the fitted cytofluorogram slope from an individual cell (>24 cells per group). (**i-j**) Representative images of NS4B and NS3 staining in mutant DENV-2 T195A/A222V/S238F-infected cells. NS4B was probed by (**i**) mAb 44-4-7 or (**j**) 10-3-7. Scale bar, 20 μm. The fluorescence ratiometric color scale ranges from 2.0 (dark purple) to 6.0 (bright yellow). (**k-l**) Cytofluorogram analysis of NS3 and NS4B within mutant virus T195A/A222V/S238F-infected cells. NS4B was probed by mAb 10-3-7 (**k**) or 44-4-7 (**l**). Each dot represents the cytofluorogram slope of an individual cell. In total, 50-60 cells were analyzed. The one-way ANOVA Dunn’s with multiple comparisons was used to compare the statistical differences between groups. *p<0.05, **p<0.01, ***p<0.001, ****p<0.0001.

During viral infection, NS4B cytosolic loops exist in both the “free” and the NS3-bound forms. mAb 44-4-7 can access “free” cytosolic loops, but steric hindrance inhibits interaction with NS3-bound loops. We hypothesized that if NITD-688 disrupts the NS3/NS4B interactions within the replication complex, the cytosolic loops should become more accessible to staining by mAb 44-4-7 after NITD-688 treatment in comparison to DMSO treatment. In contrast, NITD-688 would not impact NS4B staining by mAb 10-3-7 since the epitopes recognized by mAb 10-3-7 are accessible regardless of NS3 binding (Fig. 5a).

To test this hypothesis, we conducted a delayed treatment experiment where NS4B/NS3 complexes had formed in DENV-infected cells before the addition of inhibitors (Fig. 5b). Huh7 cells are susceptible to DENV infection and also suitable for imaging analysis. As expected, NITD-688 and JNJ-1802 effectively inhibited a nanoluciferase reporter DENV-2 strain NGC (NGC-Nluc) in Huh7 cells at 48 h post-infection, with EC_50_ values of 5.39 nM and 31 pM, respectively (Fig.S10). NITD-008 had an EC_50_ of 0.38 µM in Huh7 cells, consistent with the literature^56^. For a delay treatment, Huh7 cells were infected with DENV-2 and followed by exposure to 500 nM NITD-688 (93-fold the EC_50_) or 3 nM JNJ-1802 (97-fold the EC_50_) at 20 h post-infection. Additionally, 10 µM NITD-008 (26-fold the EC_50_) served as a control to benchmark antiviral mechanism independent of NS4B. DMSO served as a negative control. We initially validated the compounds’ inhibition of viral protein expression and virion production. Total intracellular proteins from the delayed treatment experiments were extracted, denatured, and analyzed by the Western blot with mAb 44-4-7. NITD-688 reduced NS4B expression levels by 53%. NITD-008 exhibited the most inhibition, with NS4B expression reduced by 73%, while JNJ-1802 showed the least inhibition, with NS4B expression reduced only by 35% (Fig. 5c). Notably, compared to NS4B, NS3 protein expression was less reduced by all three inhibitors, probably due to the distinct half-life of different viral proteins. Correspondingly, reductions in viral RNA production were detected, with the most substantial decrease observed with NITD-008 treatment, followed by NITD-688, and JNJ-1802 (Fig. 5d).

Next, we applied immunofluorescence staining to detect intracellular viral protein under the same experimental conditions. NS4B was stained using mAb 10-3-7 or mAB 44-4-7, with NS3 staining used for internal normalization. NS4B displayed perinuclear co-localization with NS3 across all experimental groups (Fig. 5e, f). Consistent with the Western blot results, NITD-688 reduced NS3 signals compared to DMSO, indicating its inhibition of viral replication. More importantly, profound differences in NS4B staining were observed in NITD-688-treated cells. Compared to DMSO-treated cells, NS4B levels, consistent with NS3 levels, decreased considerably in NITD-688-treated cells when probed with mAb 10-3-7; conversely, NS4B staining appeared much brighter when probed with mAb 44-4-7 (Fig. 5e, f). This characteristic was unique to the NITD-688 treatment. JNJ-1802 displayed negligible impact on NS3 or NS4B staining, while NITD-008 markedly reduced both NS3 and NS4B signals to much lower levels, irrespective of staining with mAb 10-3-7 or 44-4-7. Further pixel correlation analysis confirmed this observation (Fig. S11). Specifically, after NITD-688 treatment, the averaged fitted slope of the cytofluorogram from individual cells, where a higher slope indicates a stronger NS4B signal at a given NS3 intensity, shifted drastically from 0.3 to 4.16 when NS4B was probed with mAb 10-3-7 or with mAb 44-4-7, respectively (Fig. 5g, h). In contrast, treatments with JNJ-1802 and NITD-008 did not significantly change the cytofluorogram slope (1.28 vs. 1.87) when NS4B was probed with mAb 10-3-7 or 44-4-7. The striking disparity observed in NS4B staining after NITD-688 treatment, as detected through immunostaining versus Western blot analysis with mAb 44-4-7, strongly supports the hypothesis that during DENV infection, NS4B cytosolic loops are largely inaccessible to mAb 44-4-7 due to the NS3 binding and that NITD-688 treatment disrupts preformed NS4B/NS3 complexes, thereby freeing the cytosolic loop of NS4B and rendering it more accessible to mAb 44-4-7.

Consistent with the inability of NITD-688 to bind NS4B featuring resistance mutations, staining of the NS4B triple mutant T195A/A222V/S238F with mAb 44-4-7 was unaffected by treatment with NITD-688 (Fig. 5i-l). Specifically, mutant NS4B signals were comparable between DMSO- and NITD-688-treated cells, regardless of which mAb was used for staining (Fig. 5i, j). Quantitative analysis revealed that compared to DMSO, NITD-688 did not alter the cytofluorogram slope when mutant NS4B was probed with either mAb (Fig. 5k, l). JNJ-1802 or NITD-008 treatment did not change the staining of mutant NS4B either, although NITD-008 notably reduced the overall expression of NS4B and NS3 in mutant virus-infected cells, similar to its effects on WT DENV-2. These results are consistent with a mechanism in which NITD-688 disrupts NS4B/NS3 complexes in DENV-infected cells through direct binding to NS4B.

### NITD-688 remains effective in inhibiting DENV during delayed treatment in cellular assays

NS4B/NS3 complexes accumulate over time during DENV infection. Considering the ability of NITD-688 to disrupt preformed NS4B/NS3 complexes, we hypothesized that this inhibitor would remain effective at blocking viral replication during later stages of infection. In line with this notion, Fig. 5g indicates that NITD-688 effectively inhibits DENV replication when treatment was initiated 20 h post-infection. To delve deeper into the biological significance of NITD-688’s antiviral mechanism, we compared its effectiveness to JNJ-1802 in two additional antiviral experiments. NITD-008 served as a control inhibitor with an inhibitory mechanism independent of NS4B.

In the initial experiment (Fig. 6a), A549 cells were exposed to inhibitors from 1 to 20 h post-infection with NGC-Nluc. Luciferase activities were measured 24 h post-infection to assess the impact of delayed addition on the antiviral activity of NITD-688, JNJ-1802, and NITD-008. As expected, activity declined when inhibitors were administered later during the infection course. However, notable differences in the magnitudes of activity loss were observed between the two NS4B inhibitors (Fig. 6b, c). The potency of NITD-688, similar to NITD-008, began to decrease when compounds were added 12 h post-infection, reaching a maximal decrease of 21.0-fold when compounds were added 20 h post-infection. In contrast, the potency of JNJ-1802 started to decrease at 8 h post-infection and ultimately decreased by over 96.8-fold when JNJ-1802 was added 20 h post-infection. This data shows that when tested in a cellular infection assay, the potency of NITD-688 was less impacted than JNJ-1802 in delayed treatment.

**Fig 6.**
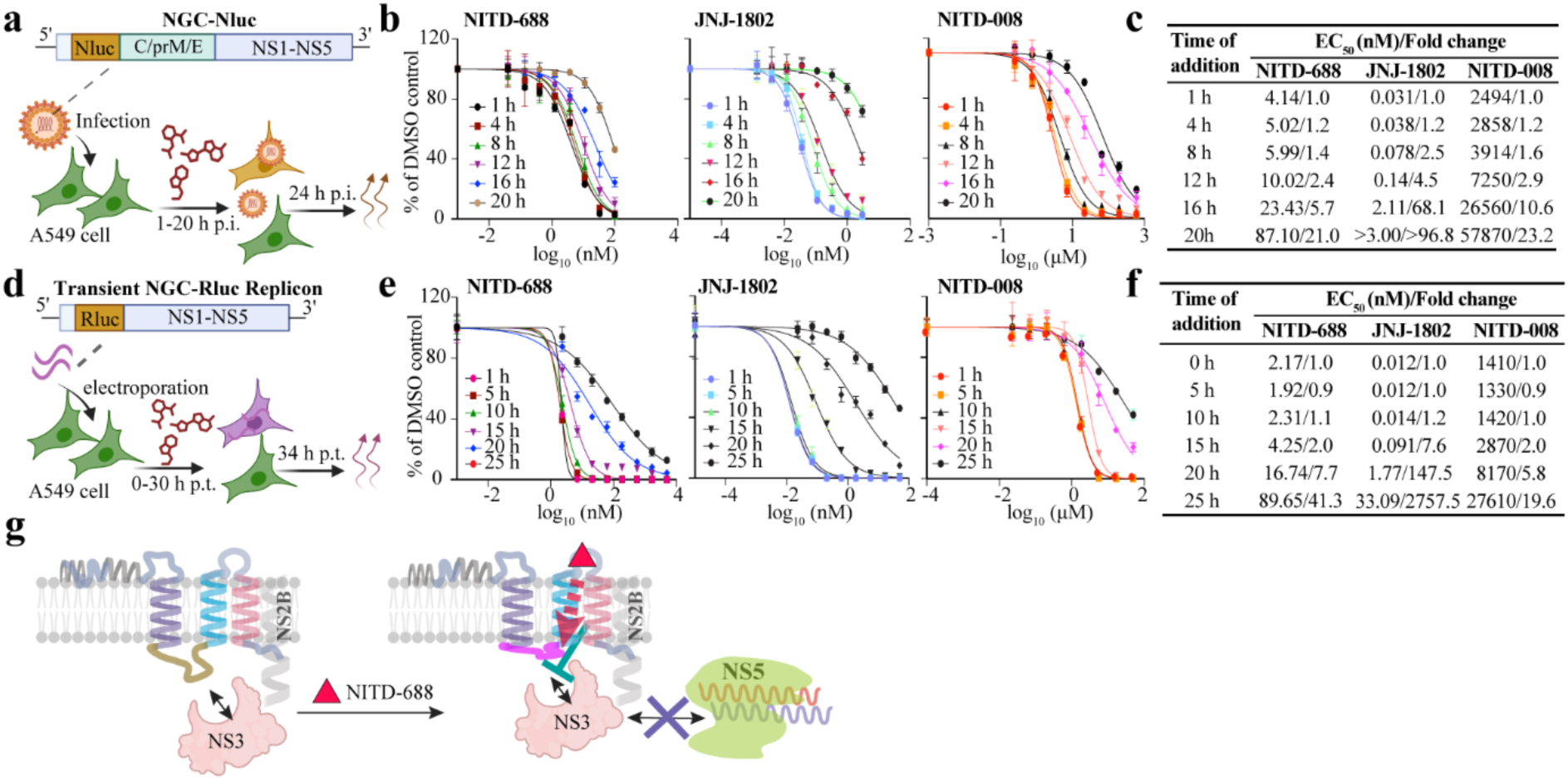
Antiviral activity of NITD-688 in cell cultures. **a** Scheme for DENV-2 infection followed by inhibitor treatment at various times of addition. Cells were infected with DENV-2 NGC containing a Nanoluciferase reporter (NGC-Nluc)^75^. At indicated time points, a series of diluted inhibitors was added. EC_50_s of each inhibitor at each time of addition were determined at 24 h post-infection. **b** Dose-response curves of inhibitors against NGC-Nluc. **c** Summary of EC_50_s values and fold changes of inhibitors against NGC-Nluc. For each inhibitor, the fold change in EC_50_ was calculated by comparing EC_50_ values at each given time point to that at one hour post-infection. **d** Scheme for the time of addition after transfection of DENV-2 replicon RNA. Cells were electroporated with *in vitro* transcribed DENV-2 NGC-Rluc replicon RNAs^67^, in which viral structural proteins were replaced with a Renilla luciferase gene (NGC-Rluc replicon). At the indicated time points, a series of diluted inhibitors was added. At 34 h post-infection, EC_50_s of each inhibitor at each given time of addition were determined. **e** Dose-response curves of inhibitors against transient NGC-Rluc replicon. **f** Summary of EC_50_ values and fold change of inhibitors against transient NGC-Rluc replicon. For each inhibitor, the fold change in EC_50_ was calculated by comparing the EC_50_ values at each given time point to that of immediate treatment (0 h) after electroporation. **g** A proposed working model for NITD-688.

Subsequently, a transient transfection experiment was performed in A549 cells using an *in vitro* synthesized RNA of a DENV-2 NGC-Rluc replicon (Fig. 6d). In this replicon, the structural genes of DENV were deleted, resulting in a loss of virion production upon transfection into cells, thereby eliminating the multiple cycles of re-infection. This approach allows for the analysis of compound effects on viral replication solely within cells containing viral RNAs. Transfected cells were treated with inhibitors from the time immediately following electroporation (0 h) to 25 h post-electroporation. EC_50_ values measured at each time point of addition were determined 34 h after transfection before the signal plateaued. Consistently, NITD-688, similar to NITD-008, showed a later and smaller decrease in potency compared to JNJ-1802 when compound treatment was delayed (Fig. 6e, f). Specifically, when compounds were added 25 h post-electroporation, the efficacy of NITD-688 was reduced by 41.3-fold, while that of JNJ-1802 was decreased by 2758.3-fold.

Collectively, these findings underscore the promise of NITD-688 as a robust inhibitor against DENV, particularly in delayed treatment scenarios.

## Discussion

NITD-688 is currently in Phase II clinical trials for treating patients with dengue fever. While NS4B was known as the target of NITD-688^19^, the explicit molecular mechanism remained undefined. This study demonstrates that NITD-688 disrupts the interaction between mature NS4B and NS3 in biochemical and cellular assays. NITD-688 can block the formation of NS4B/NS3 complexes and disrupt preformed complexes. Unsurprisingly, NITD-688 retains potency under delayed treatment conditions where preformed NS4B/NS3 complexes exist.

Previous studies demonstrated that NITD-688 directly binds to NS4B of DENV-2 by NMR^19^. Consistent with these results, our ITC data demonstrates that NITD-688 binds to NS4B from all four DENV serotypes. In addition, the binding affinity of NITD-688 to NS4B in general correlates with its potency in cellular assays^19^, showing the weakest affinity and potency against DENV-1 and the strongest against DENV-2 and DENV-3. NS4B is an integral membrane protein residing in the ER membrane, characterized by three biochemically validated transmembrane segments^57^. The NS4B/NS3 interaction has been demonstrated to be essential for viral replication^44,58^. A single mutation, Q134A, in DENV NS4B can abolish the NS4B/NS3 interaction and DENV RNA replication^58^. Additionally, DENV NS4B interacts with NS3, dissociating it from single-stranded RNA and enhancing NS3 helicase activity *in vitro*^59^. It has been recently shown that the interaction between NS2B/NS3 and precursor NS4A-2K-NS4B functionally associates with vesicle packet (VP) formation^60^. However, it remains unclear how the interaction NS2B/NS3 and NS4B-2K-NS4B mediates the VP formation, given the short half-life and low abundance of the precursor in infected cells. Our study has focused on the mature NS4B because it represents the most abundant species during DENV infection. Our biochemical assays further demonstrate that mature NS4B interacts with NS3 through the cytosolic loop (peptide 121-171) of NS4B *in vitro*. Additionally, for the first time, we have shown that the cytosolic loops within mature NS4B largely bind NS3 in cells infected with DENV-2 (Fig.5). When this interaction is interrupted by NITD-688 after infection, viral replication is significantly reduced. These new pieces of evidence underscore the biological significance of the mature NS4B and NS3 interaction during DENV infection.

*In vitro* selection for resistance using NITD-688 revealed five unique amino acid substitutions in NS4B that are prevalent in response to the selective pressure of NITD-688. Of those, T195A, T215A, and A222V were revealed as the residues contributing the most to resistance, as recombinant NS4B protein bearing these individual mutations showed the largest decreases in binding affinity of NITD-688, and recombinant viruses containing these single mutations resulted in the largest increases in EC_50_. In contrast, A193V and W205L each only slightly decreased the binding affinity of NITD-688 to NS4B and resulted in smaller EC_50_ fold shifts against recombinant viruses. Interestingly, while T195A and T215A substitutions resulted in roughly equivalent decreases in binding affinity of NITD-688 to mutant NS4B as well as roughly equivalent increases in EC_50_ values against recombinant viruses, T195A occurred with a much higher frequency than T215A in P15 viruses (Fig. 1d, e). This could be due to the rapid emergence of the combinations of T195A/A222V with other mutations that can confer better resistance to NITD-688 or replicate better than T215A/A222V. It should be noted that the recombinant double and triple mutants exhibit strong resistance to NITD-688, similar to the P15 NITD-688 selections (Fig. 1e versus 1b). This suggests that other mutations inside and outside of NS4B may not contribute significantly to viral resistance to NITD-688.

As an orthogonal demonstration of the importance of the residues A195 and A222V for NITD-688 binding to NS4B, introducing these DENV NS4B amino acids into ZIKV rendered it sensitive to NITD-688. Specifically, NITD-688 bound to ZIKV NS4B A197T/C224A and inhibited the replication of ZIKV A197T/C224A. In contrast, WT ZIKV NS4B protein and WT ZIKV showed neither binding nor inhibition at the highest tested concentration of NITD-688. Together, these data suggest that T195, T215, and A222 play a critical role in the binding of NITD-688 to NS4B and its antiviral activity. These three substitutions (T195A, T215A, and A222V) were also identified in previous resistance selections using a precursor compound to NITD-688, Compound 1^19^, suggesting that the NITD-688 class of compounds bind to the same region of NS4B. Additionally, T195, T215, and A222 are conserved among all four serotypes of DENV, but not in other flaviviruses, further supporting the specificity of NITD-688’s inhibition of DENV.

Compelling evidence from biochemical (BLI) and cellular (co-IP and microscopy) assays demonstrates that NITD-688 disrupts the interaction between mature NS4B and NS3. NITD-688 does not significantly affect the interaction of NS4B with other viral proteins (NS1, NS2A, and NS4A) or host factors. We also demonstrate, using ITC and BLI, that the NS4B peptide 121-171 (also known as the cytosolic loop) binds NS3 with similar affinity as full-length NS4B, suggesting that this region is responsible for binding to NS3, in line with our previous SPR studies^44^. Interestingly, the residues that are most critical for the binding of NITD-688 to NS4B (T195A, T215A, and A222V) are outside of the cytosolic loop of NS4B. This suggests that NITD-688 does not disrupt the interaction between mature NS4B and NS3 by directly competing with the NS3 binding site; instead, we speculate that the binding of NITD-688 to NS4B results in a conformational change in NS4B that reduces binding to NS3, disrupting the replication complex and ultimately halting viral RNA synthesis (Fig. 6g).

Targeting NS4B/NS3 interactions was proposed as the antiviral mechanism of JNJ-1802, another DENV NS4B inhibitor in Phase II clinical trials^20^. Kiemel, D. et al recently reported that NITD-688 and JNJ-A07 (a close analog of JNJ-1802) share a similar mechanism of action by blocking interactions between NS2B/NS3 and the precursor NS4A-2K-NS4B, as well as preventing the formation of vesicle packets (VPs) at the early stage of viral replication^60^. However, our results reveal several differences between NITD-688 and JNJ-1802 at the molecular level. First, resistance mutations for these two compounds appear in different regions of NS4B, suggesting that they bind to different regions of the protein. This is supported by the fact that there is minimal cross-resistance between resistant viruses selected on each compound. NITD-688 likely binds the residues T195, T215, and A222 residing between the last two transmembrane segments of NS4B, whereas JNJ-1802 likely binds near V91/L94, close to the first transmembrane segment^20^. Notably, JNJ-1802 binds to mature NS4B (without 2K peptide) with a 4.1-fold higher affinity than NITD-688, resulting in approximately 4-fold more inhibition of NS4B/NS3 interactions in the ITC assay. While this may partially explain the ∼100-fold higher potency of JNJ-1802 in cellular assays, additional mechanisms likely account for the increased potency, such as the impact of JNJ-1802 on the VP formation or the integrity of replication vesicles^60,61^.

Second, NITD-688 has a unique ability to block the *de novo* formation of NS4B/NS3 complexes and to disrupt preformed NS4B/NS3 complexes. In contrast, JNJ-1802 only inhibits the initial formation of NS4B/NS3 complexes. Several lines of evidence support this conclusion. NITD-688 (but not JNJ-1802) inhibits the co-IP of NS3 with NS4B during delayed treatment, enhances the dissociation of NS3 from NS4B/NS3 complexes *in vitro* as observed by BLI, and releases NS4B cytosolic loops from the NS3-bound state during viral infection as observed with antibody staining experiments. Finally, likely due to its ability to disrupt preformed NS4B/NS3 complexes, the potency of NITD-688 in cell-based assays is less affected by delayed treatment than JNJ-1802. Although the underlying molecular differences in the mechanism of action of NITD-688 and JNJ-1802 remain to be determined, we speculate that NS4B may undergo a conformational change upon NS3 binding, which could affect the binding regions targeted by JNJ-1802, but not those targeted by NITD-688. Differences in the efficacy of these two compounds in delayed treatment may inform strategies for dengue treatment in the clinic and warrant further investigation.

In contrast to our results that the efficacy of NITD-688 is less affected by delayed treatment compared to JNJ-1802, Kiemel, D. et al^60^ have reported that while both NITD-688 and JNJ-A07 (a close analog of JNJ-1802) can block VP formation when cells are treated early (4 h after transfection), neither compound can disrupt the already formed VPs under delayed treatment conditions (16 h after transfection). Our study, however, demonstrates that NITD-688 can disrupt the interaction between mature NS4B and NS3, and effectively reduce viral replication even with delayed treatment, arguing that NITD-688 inhibits viral replication through a mechanism independent of VP formation. Notably, our co-IPs have shown greater sensitivity to inhibitors compared to those reported by Kiemel, D. et al^60^. This is likely due to the inclusion of inhibitors in the lysis buffers of our co-IPs, which prevents the re-association of NS4B/NS3 during the cell lysis and bead binding steps.

*In vitro* resistance selection is an important tool for determining the resistance profile of compounds that will be used to treat patients. In this study, we demonstrate that NITD-688-resistant viruses can begin emerging in cell culture as early as passage 4 (approximately 16 days post-infection), with single amino acid substitutions resulting in 3.0 to 117.4-fold shifts in potency, and triple mutants up to 723.7-fold shift in potency. While all the double and triple mutants showed significant costs to fitness in both human and mosquito cells, decreasing the likelihood they could emerge and/or spread widely, the fitness of the single mutants was less severely impacted. Thus, patients treated with NITD-688 should be closely monitored for the emergence of resistance, and efforts should be made to develop combination antivirals to mitigate the threat of resistance.

In summary, we have further elucidated the antiviral mechanism of NITD-688. NITD-688 directly binds to NS4B and disrupts NS4B/NS3 interactions, inhibiting viral replication. In contrast to JNJ-1802, NITD-688 can both block the formation of NS4B/NS3 complexes and disrupt preformed complexes. These molecular insights into the distinct antiviral mechanisms of the two leading DENV NS4B inhibitors have significant implications for developing next-generation flavivirus NS4B inhibitors.

## Methods

### Cells, antibodies, and compounds

BHK-21, HEK-293T, Huh7, Vero E6, and A549 cells were purchased from the American Type Culture Collection (ATCC, Bethesda, MD). BHK-21 and A549 cells were maintained in Roswell Park Memorial Institute (RPMI) 1640 (Gibco) medium supplemented with 10% fetal bovine serum (FBS) (HyClone Laboratories) and 1% penicillin/streptomycin (PS) at 37°C with 5% CO_2_. HEK-293T, Huh7, and Vero E6 cells are maintained in Dulbecco’s modified Eagle’s medium (DMEM) (Gibco) supplemented with 10% FBS, and 1% PS. *Aedes albopictus* C6/36 cells were grown in RPMI 1640 containing 10% FBS and 1% PS at 28°C. All mediums, antibiotics, and supplements were purchased from Gbico. Cells were tested negative for mycoplasma.

The following antibodies were used: mouse monoclonal antibodies 44-4-7 and 10-3-7 cross-reactive with DENV NS4B^55^, a rabbit polyclonal antibody cross-reactive with DENV NS3 (Cat. No. GTX124252, GeneTex), a mouse anti-Flag tag antibody (Cat. No. F3165, Sigma-Aldrich), a mouse anti-HA tag antibody (Cell Signaling Technology, Cat. No. 2367S), a mouse anti-GFP antibody (Cell Signaling Technology, Cat. No. 2955), and goat anti-rabbit or anti-mouse IgGs conjugated to horseradish peroxidase (HRP) (Sigma-Aldrich).

NITD-688, NITD-008, and JNJ-1802 were synthesized according to the published procedures^19,20,56^.

### Plasmid construction

Standard molecular biology techniques were used for plasmid constructions. For purification of the NS4B proteins, cDNA encoding the NS4B protein from DENV-1 (Genbank ID: U88536.1), DENV-2 (Genbank ID: U87411.1), DENV-3 (Genbank ID: AY099336.1), DENV-4 (Genbank ID: AF326825.1), ZIKV (Genbank ID: AY632535.2), or NS4B peptide 121-171 from DENV-2 with an N-terminal 6×His-tag was cloned into the pET28a vector (Sigma-Aldrich). Mutations were introduced into DENV-2 or ZIKV NS4B by overlapping PCRs. For purification of full-length NS3, the cDNA encoding the NS3 protein from DENV-2 strain NGC with a C-terminal 6×His-tag was cloned into the pET28a vector. Two mammalian expression vectors pCAG (Addgene) or pXJ were used to express NS1, NS2A, NS2B/NS3, NS4A, and 2K-NS4B in HEK-293T cells. For constructs pXJ-E_24_-NS1-GFP, the cDNA encoding the last 24 residues of E protein and NS1 was fused with GFP and inserted into vector pXJ^62^. For constructs pXJ-SGP-NS2A-GFP, the cDNA of NS2A was fused with the signal peptide from Gaussia luciferase (SGP) at the N-terminal and GFP at the C-terminal, and inserted into vector pXJ^29^. For construct pCAG-NS2B/NS3, the cDNA encoding NS2B-NS3 was inserted into the pCAG vector (Reference). For constructing pXJ-NS4A-Flag, the cDNA encoding NS4A with a C-terminal flag tag was cloned into the pXJ vector. For construct pXJ-2K-NS4B-HA, the cDNA encoding 2K-NS4B with a C-terminal Ha-tag was cloned into the pXJ vector. For constructing pCAG-2K-NS4B-Flag, the cDNA encoding 2K-NS4B was fused with a Flag tag at the C-terminal and inserted into the pCAG vector. T195A/A222V/S238F mutations were introduced into the pCAG-2K-NS4B-Flag by overlapping PCR. For co-expression of NS2B-NS3 and 2K-NS4B in the same plasmid, the cDNA encoding DENV-2 2B-NS3 or 2K-NS4B was cloned into a dual promoter vector pEF1a (OriGENE) using NEBuilder® HiFi DNA Assembly kit. The 2B-NS3 was inserted after the mEF1a proter, and the 2K-NS4B was inserted after the rEF1a promoter. All plasmids were validated by restriction enzyme digestion and Sanger sequencing. Primers for plasmid construction are listed in Table S3.

### Viruses

Infectious clone-derived DENV-2 strain New Guinea C (NGC) was used in this study^62^. For resistant viruses, NS4B mutation(s) were introduced by overlap PCR. Full-length DENV-2 RNAs were *in vitro* transcribed from cDNA plasmids pre-linearized by the restriction enzyme XbaI using a T7 mMESSAGE mMACHINE kit (Invitrogen). To recover the DENV-2 viruses, 10 μg of RNA transcripts was electroporated into BHK-21 cell. Cells were suspended in RPMI 1640 medium containing 10% FBS, and 1% PS in a T-175 flask, and incubated at 37℃, 5% CO_2_ for 24 h. The next day, the supernatant was replenished with a medium containing 2% FBS and 1% PS and incubated at 30°C 5% CO_2_ for additional days. Viruses were harvested on days 5 to 7 post-electroporation (p.t.) and stored at −80°C.

The reporter virus of the ZIKV Dakar strain was generated using a previously described protocol^29^. Mutations A197T and C224A were introduced into ZIKV NS4B by overlap PCR. Full-length ZIKV RNA was *in vitro* transcribed using a T7 mMESSAGE mMACHINE kit from cDNA plasmids linearized by the restriction enzyme ClaI. 10 μg of RNA transcripts was electroporated into Vero-E6 cells. Cells were suspended in DMEM medium supplemented with 10% FBS and 1% PS in a T-175 flask, and incubated at 37℃, 5% CO_2_ for 24 h. The next day, the medium was changed to DMEM medium supplemented with 2% FBS and 1% PS. Viruses were harvested on days 3 to 5 p.t. and stored at −80°C. All primers used for constructing mutant viruses are listed in Table S3.

### RNA extraction and RT-PCR

Viral RNA was extracted using TRIzol™ LS (Invitrogen) and Direct-zol™ RNA Miniprep Plus kit (Zymo Research, Irvine, CA). Viral RNAs were extracted according to the manufacturer’s instructions. The extracted RNAs were dissolved in 20 μl nuclease-free water.

To verify the mutations in the viral genome, two microlitres of RNA samples were used for reverse transcription by using the SuperScriptTM IV One-Step RT-PCR System (Invitrogen) with primer pairs (DENV-2 XhoI-F and DENV-2 NruI-R for DEVN-2, ZIKV NheI-F, and ZIKV AscI-R for ZIKV in Table S3). The RT-PCR products were gel-purified using the QIAquick Gel Extraction Kit (QIAGEN) and sent for Sanger sequencing.

To quantify viral RNA samples, quantitative RT-qPCR assays were performed using the iTaq SYBR Green One-Step Kit (Bio-Rad) with a pair of primers (DENV-2 NGC-F, DENV-2 NGC-R in Table S3) on the LightCycler 480 system (Roche) following the manufacturer’s protocols.

### Selection of NITD-688-resistant viruses

A549 cells were incubated with DENV-2 (MOI 2) in the presence of NITD-688 at a starting concentration of 10 nM or DMSO. Supernatants were collected when more than 30% of cells showed CPE. 200 µl of supernatants were used for the next infection on fresh A549 cells with increased concentrations of NITD-688. The concentrations of NITD-688 in all 15 passages are the following: Passage 1-2 (P1-2): 10 nM; P3-4: 20 nM; P5-6: 50 nM; P7: 100 nM; P8: 200 nM; P9-10: 500 nM; P11-12: 1000 nM; P13-15: 2000 nM. The P15 viruses were used for testing their sensitivity to NITD-688 and for whole-genome sequencing by Illumina next-generation sequencing (NGS). The cDNAs encoding NS4B of P4, P8, and P15 viruses were amplified using the SuperScript One-Step RT-PCR system with primer paris NS4B-Pac-F and NS4B-Pac-R (Table S3) and sent for PacBio sequencing at GENEWIZ from Azenta Life Sciences.

### Plaque assay

Plaque assay of DENV was performed according to a published protocol^46^. Briefly, BHK-21 cells were seeded with a cell density of 2×10^5^ cells per well in a 24-well plate. The next day, viral samples were 10-fold serially diluted in RPMI 1640 with 2% FBS, 1% PS, and 100 µl of each dilution was added to BHK-21 cells. The plates were incubated for 1 h at 30°C and swirled every 15 min to ensure complete infection, after which the inoculum was removed. 500 µl of 0.8% methyl cellulose overlay containing 2% FBS was added to each well, and the plates were incubated at 37°C for 5 days. The plates were then fixed with 3.7% formaldehyde for 30 min and stained with 1% crystal violet for 5 min. Visible plaques were counted to determine viral titers as plaque-forming units per ml (PFU/ml).

Plaque assay of ZIKV was performed on Vero-E6 cells. Briefly, 2×10^5^ cells per well were seeded in a 24-well plate. The next day, viral samples were 10-fold serially diluted in DMEM with 2% FBS and 1% PS. 100 µl of each dilution was added to Vero-E6 cells. The plates were incubated for 1 h at 37°C and swirled every 15 min to ensure complete infection, after which the inoculum was removed. 500 µl of 0.8% methylcellulose overlay containing 2% FBS was added to each well. The plates were incubated at 37°C for 4 days followed by fixation and staining with 1% crystal violet as described above.

### Viral growth kinetics

A549 (2×10^5^ cells) or C6/36 (6×10^5^ cells) were seeded onto 12-well plates. At 24 h after seeding, cells were infected with DENV-2 (MOI 0.1) and incubated for 1 h at 30°C (A549 cells) or 28°C (C6/36 cells). After infection, the inoculum was replaced with RPMI 1640 medium supplemented with 2% FBS and 1% PS. Cells were incubated at 37°C (A549 cells) and 28°C (C6/36 cells) for 5 days. Culture supernatants were harvested every 24 h, and viral titers were determined by plaque assay.

### HCI-CFI

The High Content Imaging Cellular Flavivirus Immunoassay (HCI-CFI) was used to determine the EC_50_ of inhibitors against DENVs. A549 cells (1.4 × 10^4^ cells) were seeded into a 96-well plate. Twenty-four hours after seeding, cells were infected with DENV (MOI 0.3) and exposed to 3-fold serially diluted inhibitors or DMSO. After incubation at 37°C for 48 h, the cells were fixed in 4% paraformaldehyde (Thermo Fisher Scientific) in PBS for 30 min at room temperature and permeabilized in permeabilization buffer (0.1% Triton X-100 in PBS) for 10 min, followed by 1 h incubation in blocking buffer (1% FBS and 0.05% Tween-20 in PBS) at room temperature. The cells were then incubated with primary antibody Flavivirus group antigen 4G2 antibody conjugated with Alexa Fluor 488 (Novus Biologicals). Cells were washed in PBS and counter-stained by Hoechst 33342 solution (Thermo Fisher Scientific). Cells were imaged and counted using a CellInsight™ CX7 High Content Analysis Platform (ThermoFisher Scientific). The infection rates in each well were normalized to that of DMSO-treated wells to calculate the percentage of infection. The EC_50_ value was determined by a four-parameter nonlinear regression model in GraphPad Prism 10.

### Sequence analysis

To calculate the natural occurrence of NITD-688 resistance mutations in the general population of DENVs, entries containing the whole genome sequence of DENV were downloaded from the Bacterial and Viral Bioinformatics Resource Center (BV-BRC) (https://www.bv-brc.org/)^50^. As of 11 December 2024, a total of 7818 whole genome sequences of DENVs were available. The numbers of genome sequences for DENV-1, −2, −3, and −4 were 3228, 2191, 1626, and 773, respectively. A multiple sequence alignment was performed for each DENV serotype using MAFFT^63^. The results were analyzed in Jalview^64^ to calculate the frequency of each NITD-688 resistance mutation in each DENV serotype or all DENV sequences. The amino acid sequence alignment of flavivirus NS4B with representative strain sequences was performed in ClustalW^65^. To calculate the consensus sequence of other flaviviral NS4B, whole genome sequences of YFV (n=444), WNV (n=2929), ZIKV (n=1195), JEV (n=477), and TBEV (n=461) were also downloaded from the BV-BRC. Using Biopython scripts, the full genome sequences of DENV-1 to DENV-4, YFV, WNV, ZIKV, JEV, and TBEV were translated into individual protein sequences. Full-length NS4B sequences were extracted, and multiple sequence alignment was performed using MAFFT. Consensus sequences were analyzed and shown using the WebLogo server^66^.

### Antiviral assay using NGC-Nluc virus

Huh7 cells (1.2×10^4^ cells/per well) were seeded in a white, opaque 96-well plate (Corning) and incubated overnight at 37°C, 5% CO_2_. The next day, the culture media were removed, and the cells were infected with 100 µl of DENV-2 NGC-Nluc (MOI 0.1) in various concentrations of inhibitors. At 48 h post-infection, luciferase signals were measured by using the NanoGlo substrate (Promega) according to the manufacturer’s instructions. Plates were read in a BioTek Cytation 5 plate reader (Agilent) with a gain of 120.

For the delay treatment experiment, A549 cells (1.5×10^4^ cells/per well) were seeded in a white, opaque 96-well plate (Corning) and incubated overnight at 37°C, 5% CO_2_. The next day, the cells were infected with 100 µl of DENV-2 NGC-Nluc (MOI 1.0). After 1 h of incubation, the virus was removed, and cells were washed with PBS. At each time point (1, 4, 8, 12, 16, and 20 h post-infection), a set of NGC-Nluc-infected cells were treated with either 0.5 % v/v DMSO or 3-fold serially diluted inhibitors: NITD-688 (starting at 100 nM), NITD-008 (starting 500 μM), or JNJ-1802 (starting at 3 nM). At 24 h post-infection, the supernatants were removed, and cells were washed with PBS. Then 50 µl of NanoGlo substrate (Promega) diluted 50 times in NanoGlo Assay Buffer (Promega) was added to the cells. Plates were read in a BioTek Cytation 5 plate reader (Agilent) with a gain of 120.

### Time of addition experiment using transient NGC-Rluc replicon

A plasmid pACYC-NGC-Rluc containing the DENV-2 NGC replicon with the Renilla luciferase gene was used^67^. *In vitro*-transcribed RNA was synthesized using a T7 mMESSAGE mMACHINE kit (ThermoFisher Scientific). 10 µg of replicon RNAs were electroporated into A549 cells^67^. Electroporated cells were plated in 96-well plates (2×10^4^ cells/well) in RPMI 1640 medium supplemented with 10% FBS and 1% PS. Immediately after seeding (0 h) or at 5, 10, 15, 20, and 25 h post-electroporation, cells were treated with either 0.5 % DMSO or 3-fold serially diluted inhibitors: NITD-688 (starting at 5 μM), NITD-008 (starting at 50 μM), or JNJ-1802 (starting at 50 nM). At 34 h after electroporation, the supernatants were removed, and cells were washed with PBS and lysed in 25 µL of Rluc lysis buffer (Promega). Fifty µl of Rluc substrate (diluted 1:100 in Rlu Assay Buffer, Promega) was added to the cell lysates. Chemilumilence signals were measured using a BioTek Cytation 5 plate reader. The signals in each well were normalized to those of DMSO-treated wells. The EC_50_ value was determined by a four-parameter nonlinear regression model in GraphPad Prism 10.

### Protein purification

NS4B proteins were purified using a previously described protocol with minor changes^57^. The plasmid was transformed into *E. coli* BL21(DE3) Competent Cells (Millipore Sigma) and grown at 37°C in LB broth. When the culture reached an optical density at 600 nm (OD_600_) of 0.6-0.8, 0.5 mM IPTC was added to induce protein expression at 16°C for 16 h. The cells were pelleted down by centrifugation at 7,000 × rpm for 10 minutes and lysed in a lysis buffer containing 20 mM HEPES, 500 mM NaCl, and pH 8.0 using a Branson Ultrasonics Sonifier. Cell lysates were clarified by centrifugation at 7000 × rpm, 4°C for 10 minutes. The supernatants were centrifuged at 45,000 × rpm in a Beckman 45Ti rotor to pellet down the membrane fractions. The membrane pellet was resuspended in the same lysis buffer supplemented with 1% LMPG by stirring for 2 h at 4°C. The solubilized membrane fraction was centrifuged at 45,000 rpm in a Beckman 45Ti rotor to remove the insoluble proteins. The supernatant was then loaded into a 5 mL Ni-NTA resin (QIAGEN). The resin was washed with a wash buffer (20 mM HEPES 7.6, 500 mM NaCl, 0.03% LMNG, 25 mM imidazole, pH 8.0) and eluted with an elution buffer (20 mM HEPES 7.6, 500 mM NaCl, 0.03% LMNG, 500 mM imidazole, pH 8.0). Fractions containing NS4B were pooled and purified by an Increase 200 size exclusion column (GE) in buffer A containing 20 mM HEPES, 150 mM NaCl, 0.03% LMNG, pH 8.0. Protein was flash-frozen in liquid nitrogen and stored at −80°C. The cytosolic loop of NS4B was purified using a previously reported protocol^44^.

The full-length DENV-2 NS3 was purified in *E.coli* BL21(DE3) using a previously reported protocol^44^. Briefly, plasmid pET28a-6×His-NS3 encoding NS3 with an N-terminal his-tag and thrombin cleavage site was transformed into *E. coli* BL21(DE3) and grown at 37°C in LB broth. When the culture reached an OD_600_ of 0.6-0.8, 0.5 mM IPTC was added to induce protein expression at 16°C for 16 h. The cells were harvested by centrifugation at 7,000× rpm for 10 minutes and lyzed in a 20 mM HEPES buffer containing 500 mM NaCl and a cocktail of protease inhibitors (Roche) using a Branson Ultrasonics Sonifier. Cell debris and unlysed cells were removed by centrifugation at 10,000× g for 60 minutes. The supernatant was collected and loaded into a 5 ml Ni-NTA resin (QIAGEN). The resin was washed with a wash buffer (20 mM HEPES 7.6, 500 mM NaCl, 25 mM imidazole, pH 8.0). Proteins were eluted in a buffer containing 20 mM HEPES, 500 mM NaCl, and 500 mM imidazole, pH 8.0. Fractions containing NS3 were pooled and purified by Increase 200 size exclusion column (GE) in buffer B containing 20 mM HEPES, and 150 mM NaCl. Protein was flash-frozen in liquid nitrogen and stored at −80°C.

### Isothermal Titration Calorimetry

The Isothermal Titration Calorimetry (ITC) was conducted using MicroCal PEAQ-ITC (Malvern). For the NS4B/inhibitor binding assay, 6 mM NITD-688 or JNJ-1802 in 100% DMSO was diluted with buffer A to a final concentration of 0.3 mM NITD-688 in 5% DMSO (v/v). NS4B was diluted in buffer A supplemented with 5% DMSO (v/v) to a final concentration of 0.03 mM. Inhibitors were titrated against NS4B.

For the NS4B/NS3 binding assay, NS4B and NS3 were buffer exchanged and diluted in buffer A to a final concentration of 0.03 mM and 0.3 mM, respectively. NS3 was titrated against NS4B. To test whether NITD-688 or JNJ-1802 can prevent NS4B/NS3 interaction, 0.03 mM of NS4B was pre-incubated with different concentrations of NITD-688 or JNJ-1802 (0.015 to 0.06 mM compound in buffer A and 5% v/v DMSO). NS3 was diluted in buffer A supplemented with 5% DMSO (v/v) to a final concentration of 0.3 mM. NS3 was titrated against NS4B/inhibitor complex. All ITC titrations were performed for 17 injections at a temperature of 25°C. A constant reference power of 40 μcal/s was applied to both the sample and reference cells. Data was analyzed by use of the MicroCal PEAQ-ITC Analysis (Malvern). The curves were fitted using a 1:1 binding model.

### Biolayer Interferometry

Biolayer Interferometry (BLI) was conducted with the Octet R8 (Sartorius) to determine the affinity and kinetics of NS4B/NS3 interaction. The purified NS4B proteins were biotinylated with NHS-PEO_4_-Biotin (ThermoFisher Scientific) according to the manufacturer’s instructions. The excess biotin was removed by a 10-kDa Amicon tube (Amicon Centrifuge) and biotin-labeled NS4B in buffer A was concentrated to 100 nM. For the NS4B/NS3 binding assay, streptavidin Octet biosensor (Sartorius) was incubated with 100 nM biotin labeled NS4B followed by a wash with buffer A for 60 seconds. The NS4B-captured biosensors were then submerged in NS3 solutions at various concentrations (3.7, 11, 33, and 100 nM) for 300 seconds (NS3 association) followed by dissociation in buffer A for 300 seconds. The curve was fitted, and *K*_D_ was estimated using the global model.

To analyze the inhibitor’s effects on the dissociation of NS4B/NS3 complexes, the NS4B-captured biosensors were submerged in 100 nM of NS3 solutions for 300 seconds (association) followed by dissociation in buffer A containing various concentrations of NITD-688 or JNJ-1802 (3.125, 12.5, 50, and 200 nM) for 600 seconds. All steps were performed at 25°C with shaking. Data was analyzed using Octet Analysis Studio (Sartorius). The curve was fitted and *k_on_*/*k_off_* was calculated using the individual model.

### Epitope binding assay

Epitope binding assay was conducted by following a previously published protocol with modifications^68^. The binding affinities of mAb 10-3-7 or 44-4-7 to NS4B were initially determined by BLI. Briefly, mAb 10-3-7 or 44-4-7 was biotinylated with NHS-PEO_4_-Biotin according to the manufacturer’s instructions. 100 nM biotin-labeled mAbs in buffer A were captured onto the streptavidin Octet biosensors. After reaching a baseline in buffer A for 60 seconds, the mAb-captured biosensors were then submerged in buffer A containing various concentrations of NS4B (0.04, 0.11, 0.33, and 1 µM) for 300 seconds followed by dissociation in buffer A for 900 seconds.

For epitope binding assay, the mAb-captured biosensors were submerged in buffer A containing 100 nM NS4B for 300 s. After 30 seconds of washing in buffer A, the biosensors were submerged in buffer A or various concentrations of NS3 (0.33 or 1 µM) for 300 seconds, followed by dissociation in buffer A for 60 seconds. All steps were performed at 25℃ with shaking. Data was analyzed using Octet Analysis Studio. The curve was fitted, and *K*_D_ was estimated using the individual model.

### Co-IP, SDS-PAGE, and Western Blot

Co-immunoprecipitation (co-IP) was conducted in HEK-293T cells to study the impact of NITD-688 on the interaction of NS4B with viral proteins. For NS4B dimerization, 2 µg pXJ-2K-NS4B-HA and 2 µg pCAG-2K-NS4B-Flag were co-transfected. For NS4B/NS4A interaction, 2 µg pXJ-2K-NS4B-HA and 2 µg pXJ-NS4A-Flag were co-transfected. For NS4B/NS1 interaction, 2 µg pXJ-2K-NS4B-HA and 2 µg pXJ-E24-NS1-GFP or pXJ-EGFP were co-transfected. For NS4B/NS2A interaction, 2 µg pXJ-2K-NS4B-HA and 2 µg pXJ-SPG-NS2A-EGFP or pXJ-EGFP were co-transfected. For NS4B/NS3 interaction, 2 µg pCAG-2K-NS4B-Flag or pCAG-2K-NS4B (TM)-Flag and 2 µg pCAG-NS2B/NS3 were co-transfected. All transfections were performed using the X-tremeGENE transfection reagent (Roche) according to the manufacturer’s instructions. 24 h after transfection, cells were harvested and lysed in an IP lysis buffer (20 mM Tris pH 8.0, 150 mM NaCl, 1% DDM, and Roche protease inhibitor cocktail). The cell lysis was subjected to the following immunoprecipitation according to the manufacturer’s instructions: 1) anti-Flag magnetic beads (Pierce, Rockford, IL) for analyzing NS4B dimerization, NS4A/NS4B, and NS4B/NS3 interactions; 2) anti-GFP beads (ThermoFisher Scientific) for analyzing NS4B/NS1 and NS4B/NS2A interactions. After extensive washes, proteins were eluted in 2 × LDS sample buffer (ThermoFisher Scientific) with 100 µM dithiothreitol at 95°C for 15 min.

For the delayed treatment experiment, HEK-293T cells were co-transfected with 1 µg pCAG-2K-NS4B-Flag and 1 µg pCAG-NS2B-NS3 using X-tremeGene transfection regent. Four hours post-transfection, the medium was replaced with fresh DMEM containing 10% FBS to remove excess transfection reagents. In the early-treatment group, cells were immediately incubated with a medium containing either DMSO (0.1%), NITD-688 (5 nM or 100 nM), or JNJ-1802 (0.5 nM or 10 nM) and further incubated at 37°C, 5% CO_2_ for additional 44 hours. For the delayed treatment group, cells were treated with DMSO or the same concentrations of inhibitors at 24 h post-transfection, followed by incubation at 37°C with 5% CO_2_ for an additional 24 hours. At 48 h post-transfection, cells were collected and lysed in the IP buffer containing DMSO or inhibitors at the same concentrations as those used during cell treatments. The cell lysates were then subjected to co-immunoprecipitation using anti-flag magnetic beads according to the manufacturer’s instructions. Proteins were finally eluted and analyzed by SDS-PAGE and Western blot.

For SDS-PAGE and Western blot analysis, protein samples were resolved by SDS-polyacrylamide gel electrophoresis and transferred onto a PVDF membrane using a Trans-Blot Turbo Transfer System (Bio-Rad, Hercules, CA). The membranes were blocked in TBS-T containing 5% non-fat dry milk (NFDM) (Sigma-Aldrich) buffer for 1 h followed by 1 h incubation with corresponding primary antibodies diluted in TBS-T buffer containing 5% NFDM. After three TBS-T washes, the membranes were incubated with secondary antibodies for 1 h. After three additional TBS-T washes, chemiluminescent signals were developed using SuperSignal Femto Maximum Sensitivity Substrate (ThermoFisher Scientific) and detected in a ChemiDoc Imaging system (Bio-Rad). Densitometry analysis was conducted using ImageJ. Comparisons of groups were performed using one-way ANOVA, ∗p < 0.05, ∗∗p < 0.01, ∗∗∗p < 0.001, ∗∗∗∗p < 0.0001.

### IP-MS

The impact of NITD-688 on NS4B/host factor interactions was evaluated by immunoprecipitation-mass spectrometry (IP-MS). HEK-293T cells were transfected with 2 µg pCAG-2K-NS4B-Flag for 24 h and treated with DMSO or NITD-688 (10 mM) for 2 h. Then the cells were lysed at 4°C for 1 h in a lysis buffer (20 mM Tris, pH 7.5, 100 mM NaCl, 0.5% DDM, and EDTA-free protease inhibitor cocktail) containing 10 mM NITD-688 or DMSO accordingly. The lysates were clarified by centrifugation at 13000× rpm at 4°C for 15 min and subjected to immunoprecipitation using anti-Flag antibody-conjugated magnetic beads (Thermo Fisher Scientific) at 4°C for 4 h. The magnetic beads were then washed with ice-cold lysis buffer 5 times. The bound proteins on beads were eluted by boiling in 1× Laemmli sample buffer (Bio-Rad) at 95°C for 10 min. For mass spectrometry analysis, samples were reduced, alkylated, and digested using trypsin on an S-trap as described ^69^. The resultant peptides were quantified and normalized within sets by the Quantitative Fluorometric Peptide Assay. Samples were analyzed using a 60-minute gradient on our Orbitrap Eclipse. The resulting Liquid chromatography-mass spectrometry (LC-MS) data was processed in Proteome Discoverer 2.5 and SEQUEST using a human database. The minora node was used to perform Label-Free Quantitation (LFQ) using the MS peak areas for each of the peptide-spectral matches. The IP-MS was performed in triplicates. Table S4 shows the details of IP-MS results. Per replicate, protein abundance refers to the summed apex (maximum) intensity of the assigned quantified peptides.

### Confocal imaging analysis

Huh7 cells were infected with DENV2 (MOI 5). At 20 h post-infection, cells were treated with DMSO, NITD-688 (500 nM), JNJ-1802 (3 nM) or NITD-008 (10 μM) for another 16 h. The cells were then fixed in 4% paraformaldehyde, permeabilized in 0.1% Triton X-100, and stained with NS4B (mAb 10-3-7 or 44-4-7) and NS3 antibodies accordingly. High-resolution z-stack micrographs were collected using Yokogawa spinning disk confocal with a UIS2XLine Plan Apochromat (UPLAPO) 100×oil objective (NA 1.5, Olympus) and a 3.2× Magchanger. The images were deconvoluted using the cellSens software using a maximum-likelihood algorithm. Multi-point live cell imaging was performed in an onstage incubator (Tokai Hit). Cells were imaged as z-stacks in timelapse with a single shot z-drift correction before each imaging cycle to prevent focus drift.

Automated image analysis was performed with homemade macros in FIJI. For the analysis of timelapse movies, in-focus slices were selected from a z-stack at each timepoint based on normalized variance, using the autofocus hyperstack macro developed by R. Mort^70^. Cell segmentation for tracking and measurements was performed using the trainable Weka segmentation tool^71^. Pearson’s correlation analysis and the extraction of the cytofluorogram slope for each cell were performed using the JACoP plugin^72^. The evaluation of statistical significance was performed with one-way ANOVA using GraphPad Prism 10.

### Structure modeling and illustration

The atomic model of DENV-2 NS4B and ZIKV NS4B were predicted by Alphafold2^73^. All of the illustrations and alignment of atomic models were generated using the Chimera^74^.

### Statistics

Sample size was estimated based on similar research reported in the literature and no statistical method was used to predetermine sample size. No data were excluded from the analysis. All experiments were performed using at least two biological replicates to ensure reproducibility. All samples were analyzed equally with no subsampling. Investigators were generally not blinded as the experimental conditions required investigators to know the identity of the samples. Description of the statistical tests performed for each experiment and the p-values can be found in the corresponding figure legends. Dots on the bar graphs represent the values from an individual replicate of the experiment. All data were analyzed using the software Prism 10 (GraphPad). Representative data from repeated independent experiments are presented as mean ± standard deviation (SD). For viral growth kinetics analysis, log_10_-transformed values were obtained to achieve a normal distribution before being subject to statistical analysis.

## Data Availability

The mass spectrometry data are available via ProteomeXchange with the identifier PXD053801. Additional relevant data are provided in the source data file accompanying this paper.

## Acknowledgments

We thank our colleagues from Novartis and the University of Texas Medical Branch (UTMB) for their valuable discussions throughout this study. We thank Professor Dr. Michael P. Sheetz for allowing us to access his confocal microscopes. W.K.R. and the UTMB Mass Spectrometry Facility receives support from Cancer Prevention Research Institute of Texas (CPRIT) Grant number RP190682. P.Y.S. and X.X. were supported by NIH grants U19AI171413, and awards from the Sealy & Smith Foundation, the Kleberg Foundation, the John S. Dunn Foundation, the Amon G. Carter Foundation, the Gilson Longenbaugh Foundation, the Summerfield Robert Foundation.

## Author contributions

X.X. conceived and supervised the study. Y.W., L.S., J.Z., and X.X. acquired resistant mutant data. Y.W., L.F., and D.B. acquired antiviral activity data. Y.W. and L.S. generated the co-IP data. Y.W., J.Z., and Y.H. purified the proteins and generated the biophysical data. L.S. and Y.W. acquired and analyzed the confocal imaging data. Y.W. and S.J.F. analyzed the NGS data. C.S. and S.A.M. provided the inhibitors. L.S., L.K.P., and W.K.R. acquired and analyzed the IP-MS data. Y. W. and J.Y. did the sequence alignment. Y.W., L.S., Y.W., S.J.F., and X.X. generated figures. Y.W., L.S., and X.X. wrote the original draft. All authors reviewed and edited the manuscript. P.-Y. S. and X.X. acquired funding.

## Competing interests

D.B., S.A.M., and C.S. are employees of Novartis and may receive stock. UTMB has filed a patent application entitled “Stable reporter flavivirus” (application number US17/412,900) and P.-Y.S. and X.X. are the co-inventors. Other authors declare no conflicts of interest.

## Reference

1 The, L. Dengue: the threat to health now and in the future. Lancet 404, 311 (2024). 10.1016/S0140-6736(24)01542-3

2 Bhatt, S. et al. The global distribution and burden of dengue. Nature 496, 504–507 (2013).

3 Wilder-Smith, A., Ooi, E.-E., Horstick, O. & Wills, B. Dengue. The Lancet 393, 350–363 (2019).

4 Murray, N. E. A., Quam, M. B. & Wilder-Smith, A. Epidemiology of dengue: past, present and future prospects. Clinical epidemiology, 299–309 (2013).

5 Organization, W. H. Global strategy for dengue prevention and control 2012-2020. (2012).

6 Stanaway, J. D. et al. The global burden of dengue: an analysis from the Global Burden of Disease Study 2013. The Lancet infectious diseases 16, 712–723 (2016).

7 Sangkaew, S. et al. Risk predictors of progression to severe disease during the febrile phase of dengue: a systematic review and meta-analysis. The Lancet Infectious Diseases 21, 1014–1026 (2021).

8. Organization, W. H. Disease Outbreak News; Dengue – Global situation. (2023).

9 Halstead, S. B., Rojanasuphot, S. & Sangkawibha, N. Original antigenic sin in dengue. Am J Trop Med Hyg 32, 154–156 (1983). 10.4269/ajtmh.1983.32.154

10 Thomas, S. J. & Yoon, I. K. A review of Dengvaxia(R): development to deployment. Hum Vaccin Immunother 15, 2295–2314 (2019). 10.1080/21645515.2019.1658503

11 Rivera, L. et al. Three-year Efficacy and Safety of Takeda’s Dengue Vaccine Candidate (TAK-003). Clin Infect Dis 75, 107–117 (2022). 10.1093/cid/ciab864

12 Kirkpatrick, B. D. et al. The live attenuated dengue vaccine TV003 elicits complete protection against dengue in a human challenge model. Sci Transl Med 8, 330ra336 (2016). 10.1126/scitranslmed.aaf1517

13 Kallas, E. G. et al. Safety and immunogenicity of the tetravalent, live-attenuated dengue vaccine Butantan-DV in adults in Brazil: a two-step, double-blind, randomised placebo-controlled phase 2 trial. Lancet Infect Dis 20, 839–850 (2020). 10.1016/S1473-3099(20)30023-2

14 Walsh, M. R. et al. Safety and durable immunogenicity of the TV005 tetravalent dengue vaccine, across serotypes and age groups, in dengue-endemic Bangladesh: a randomised, controlled trial. Lancet Infect Dis 24, 150–160 (2024). 10.1016/S1473-3099(23)00520-0

15 Liyanage, P. et al. Evaluation of intensified dengue control measures with interrupted time series analysis in the Panadura Medical Officer of Health division in Sri Lanka: a case study and cost-effectiveness analysis. Lancet Planet Health 3, e211–e218 (2019). 10.1016/S2542-5196(19)30057-9

16 Ouedraogo, S. et al. Evaluation of Effectiveness of a Community-Based Intervention for Control of Dengue Virus Vector, Ouagadougou, Burkina Faso. Emerg Infect Dis 24, 1859–1867 (2018). 10.3201/eid2410.180069

17 Gunale, B. et al. An observer-blind, randomised, placebo-controlled, phase 1, single ascending dose study of dengue monoclonal antibody in healthy adults in Australia. Lancet Infect Dis (2024). 10.1016/S1473-3099(24)00030-6

18 Lim, S. P. et al. Ten years of dengue drug discovery: progress and prospects. Antiviral Res 100, 500–519 (2013). 10.1016/j.antiviral.2013.09.013

19 Moquin, S. A. et al. NITD-688, a pan-serotype inhibitor of the dengue virus NS4B protein, shows favorable pharmacokinetics and efficacy in preclinical animal models. Science Translational Medicine 13, eabb2181 (2021).

20 Goethals, O. et al. Blocking NS3–NS4B interaction inhibits dengue virus in non-human primates. Nature 615, 678–686 (2023).

21 Kaptein, S. J. F. et al. A pan-serotype dengue virus inhibitor targeting the NS3-NS4B interaction. Nature 598, 504–509 (2021). 10.1038/s41586-021-03990-6

22 Landovitz, R. J., Scott, H. & Deeks, S. G. Prevention, treatment and cure of HIV infection. Nat Rev Microbiol 21, 657–670 (2023). 10.1038/s41579-023-00914-1

23 Manns, M. P. & Maasoumy, B. Breakthroughs in hepatitis C research: from discovery to cure. Nat Rev Gastroenterol Hepatol 19, 533–550 (2022). 10.1038/s41575-022-00608-8

24 Lim, S. P. Dengue drug discovery: Progress, challenges and outlook. Antiviral Res 163, 156–178 (2019). 10.1016/j.antiviral.2018.12.016

25 Lindenbach, B. D. & Rice, C. M. Genetic interaction of flavivirus nonstructural proteins NS1 and NS4A as a determinant of replicase function. J Virol 73, 4611–4621 (1999). 10.1128/JVI.73.6.4611-4621.1999

26 Miller, S., Sparacio, S. & Bartenschlager, R. Subcellular localization and membrane topology of the Dengue virus type 2 Non-structural protein 4B. J Biol Chem 281, 8854–8863 (2006). 10.1074/jbc.M512697200

27 Osawa, T., Aoki, M., Ehara, H. & Sekine, S. I. Structures of dengue virus RNA replicase complexes. Mol Cell 83, 2781–2791 e2784 (2023). 10.1016/j.molcel.2023.06.023

28 Gebhard, L. G. et al. A Proline-Rich N-Terminal Region of the Dengue Virus NS3 Is Crucial for Infectious Particle Production. J Virol 90, 5451–5461 (2016). 10.1128/JVI.00206-16

29 Xie, X. et al. Dengue NS2A Protein Orchestrates Virus Assembly. Cell Host Microbe 26, 606–622 e608 (2019). 10.1016/j.chom.2019.09.015

30 Munoz-Jordan, J. L. et al. Inhibition of alpha/beta interferon signaling by the NS4B protein of flaviviruses. J Virol 79, 8004–8013 (2005). 10.1128/JVI.79.13.8004-8013.2005

31 Ashour, J., Laurent-Rolle, M., Shi, P. Y. & Garcia-Sastre, A. NS5 of dengue virus mediates STAT2 binding and degradation. J Virol 83, 5408–5418 (2009). 10.1128/JVI.02188-08

32 Falgout, B., Pethel, M., Zhang, Y. M. & Lai, C. J. Both nonstructural proteins NS2B and NS3 are required for the proteolytic processing of dengue virus nonstructural proteins. J Virol 65, 2467–2475 (1991). 10.1128/JVI.65.5.2467-2475.1991

33 Li, H., Clum, S., You, S., Ebner, K. E. & Padmanabhan, R. The serine protease and RNA-stimulated nucleoside triphosphatase and RNA helicase functional domains of dengue virus type 2 NS3 converge within a region of 20 amino acids. J Virol 73, 3108–3116 (1999). 10.1128/JVI.73.4.3108-3116.1999

34 Issur, M. et al. The flavivirus NS5 protein is a true RNA guanylyltransferase that catalyzes a two-step reaction to form the RNA cap structure. RNA 15, 2340–2350 (2009). 10.1261/rna.1609709

35 Ray, D. et al. West Nile virus 5’-cap structure is formed by sequential guanine N-7 and ribose 2’-O methylations by nonstructural protein 5. J Virol 80, 8362–8370 (2006). 10.1128/JVI.00814-06

36 Ackermann, M. & Padmanabhan, R. De novo synthesis of RNA by the dengue virus RNA-dependent RNA polymerase exhibits temperature dependence at the initiation but not elongation phase. J Biol Chem 276, 39926–39937 (2001). 10.1074/jbc.M104248200

37 Noble, C. G., Seh, C. C., Chao, A. T. & Shi, P. Y. Ligand-bound structures of the dengue virus protease reveal the active conformation. J Virol 86, 438–446 (2012). 10.1128/JVI.06225-11

38 Lim, S. P., Noble, C. G. & Shi, P. Y. The dengue virus NS5 protein as a target for drug discovery. Antiviral Res 119, 57–67 (2015). 10.1016/j.antiviral.2015.04.010

39 Good, S. S. et al. Evaluation of AT-752, a Double Prodrug of a Guanosine Nucleotide Analog with In Vitro and In Vivo Activity against Dengue and Other Flaviviruses. Antimicrob Agents Chemother 65, e0098821 (2021). 10.1128/AAC.00988-21

40 Bhardwaj, T., Kumar, P. & Giri, R. Investigating the conformational dynamics of Zika virus NS4B protein. Virology 575, 20–35 (2022). 10.1016/j.virol.2022.08.005

41 Wang, Y., Xie, X. & Shi, P. Y. Flavivirus NS4B protein: Structure, function, and antiviral discovery. Antiviral Res 207, 105423 (2022). 10.1016/j.antiviral.2022.105423

42 Arakawa, M. et al. Flavivirus recruits the valosin-containing protein-NPL4 complex to induce stress granule disassembly for efficient viral genome replication. J Biol Chem 298, 101597 (2022). 10.1016/j.jbc.2022.101597

43 Youn, S. et al. Evidence for a genetic and physical interaction between nonstructural proteins NS1 and NS4B that modulates replication of West Nile virus. J Virol 86, 7360–7371 (2012). 10.1128/JVI.00157-12

44 Zou, J. et al. Mapping the Interactions between the NS4B and NS3 proteins of dengue virus. J Virol 89, 3471–3483 (2015). 10.1128/JVI.03454-14

45 Zou, J. et al. Characterization of dengue virus NS4A and NS4B protein interaction. J Virol 89, 3455–3470 (2015). 10.1128/JVI.03453-14

46 Xie, X. et al. Inhibition of dengue virus by targeting viral NS4B protein. J Virol 85, 11183–11195 (2011). 10.1128/JVI.05468-11

47 Wang, Q. Y. et al. Discovery of Dengue Virus NS4B Inhibitors. J Virol 89, 8233–8244 (2015). 10.1128/JVI.00855-15

48 van Cleef, K. W. et al. Identification of a new dengue virus inhibitor that targets the viral NS4B protein and restricts genomic RNA replication. Antiviral Res 99, 165–171 (2013). 10.1016/j.antiviral.2013.05.011

49 Ackaert, O. et al. Safety, Tolerability, and Pharmacokinetics of JNJ-1802, a Pan-serotype Dengue Direct Antiviral Small Molecule, in a Phase 1, Double-Blind, Randomized, Dose-Escalation Study in Healthy Volunteers. Clin Infect Dis 77, 857–865 (2023). 10.1093/cid/ciad284

50 Olson, R. D. et al. Introducing the Bacterial and Viral Bioinformatics Resource Center (BV-BRC): a resource combining PATRIC, IRD and ViPR. Nucleic Acids Res 51, D678–d689 (2023). 10.1093/nar/gkac1003

51 Johnston, H. D. The thermodynamics (log K, [Delta]H°, [Delta]S°, [Delta]Cp°) of metal ligand interaction in aqueous solution.|nI.|p Design and construction of an isothermal titration calorimeter.|nII.|pThe interaction of cyanide ion with bivalent nickel, zinc, cadmium and mercury.|nIII.|pThe interaction of glycinate ion with bivalent manganese, iron, cobalt, nickel, copper, zinc and cadmium PhD thesis, Brigham Young University (1968).

52 Zou, J. et al. Dimerization of flavivirus NS4B protein. J Virol 88, 3379–3391 (2014). 10.1128/JVI.02782-13

53 Shah, P. S. et al. Comparative Flavivirus-Host Protein Interaction Mapping Reveals Mechanisms of Dengue and Zika Virus Pathogenesis. Cell 175, 1931–1945 e1918 (2018). 10.1016/j.cell.2018.11.028

54 Concepcion, J. et al. Label-free detection of biomolecular interactions using BioLayer interferometry for kinetic characterization. Comb Chem High Throughput Screen 12, 791–800 (2009). 10.2174/138620709789104915

55 Xie, X. et al. Generation and characterization of mouse monoclonal antibodies against NS4B protein of dengue virus. Virology 450-451, 250–257 (2014). 10.1016/j.virol.2013.12.025

56 Yin, Z. et al. An adenosine nucleoside inhibitor of dengue virus. Proc Natl Acad Sci U S A 106, 20435–20439 (2009). 10.1073/pnas.0907010106

57 Li, Y. et al. Secondary Structure and Membrane Topology of the Full-Length Dengue Virus NS4B in Micelles. Angew Chem Int Ed Engl 55, 12068–12072 (2016). 10.1002/anie.201606609

58 Chatel-Chaix, L. et al. A Combined Genetic-Proteomic Approach Identifies Residues within Dengue Virus NS4B Critical for Interaction with NS3 and Viral Replication. J Virol 89, 7170–7186 (2015). 10.1128/JVI.00867-15

59 Umareddy, I., Chao, A., Sampath, A., Gu, F. & Vasudevan, S. G. Dengue virus NS4B interacts with NS3 and dissociates it from single-stranded RNA. J Gen Virol 87, 2605–2614 (2006). 10.1099/vir.0.81844-0

60 Kiemel, D. et al. Pan-serotype dengue virus inhibitor JNJ-A07 targets NS4A-2K-NS4B interaction with NS2B/NS3 and blocks replication organelle formation. Nat Commun 15, 6080 (2024). 10.1038/s41467-024-50437-3

61 Kiemel, D. *Elucidation of the mechanism-of-action of highly potent Dengue virus NS4B inhibitors JNJ-A07 and JNJ-*1802 doctoral degree thesis, Ruprecht - Karls - University, (2024).

62 Xie, X., Gayen, S., Kang, C., Yuan, Z. & Shi, P.-Y. Membrane topology and function of dengue virus NS2A protein. Journal of virology 87, 4609–4622 (2013).

63 Katoh, K. & Standley, D. M. MAFFT multiple sequence alignment software version 7: improvements in performance and usability. Molecular biology and evolution 30, 772–780 (2013).

64 Waterhouse, A. M., Procter, J. B., Martin, D. M., Clamp, M. & Barton, G. J. Jalview Version 2— a multiple sequence alignment editor and analysis workbench. Bioinformatics 25, 1189–1191 (2009).

65 Larkin, M. A. et al. Clustal W and Clustal X version 2.0. bioinformatics 23, 2947–2948 (2007).

66 Crooks, G. E., Hon, G., Chandonia, J.-M. & Brenner, S. E. WebLogo: a sequence logo generator. Genome research 14, 1188–1190 (2004).

67 Ng, C. Y. et al. Construction and characterization of a stable subgenomic dengue virus type 2 replicon system for antiviral compound and siRNA testing. Antiviral research 76, 222–231 (2007).

68 Noy-Porat, T. et al. Characterization of antibody-antigen interactions using biolayer interferometry. STAR Protoc 2, 100836 (2021). 10.1016/j.xpro.2021.100836

69 Mascibroda, L. G. et al. INTS13 variants causing a recessive developmental ciliopathy disrupt assembly of the Integrator complex. Nature communications 13, 6054 (2022).

70 Sun, Y., Duthaler, S. & Nelson, B. J. Autofocusing in computer microscopy: selecting the optimal focus algorithm. Microscopy research and technique 65, 139–149 (2004).

71 Arganda-Carreras, I. et al. Trainable Weka Segmentation: a machine learning tool for microscopy pixel classification. Bioinformatics 33, 2424–2426 (2017).

72 Bolte, S. & Cordelières, F. P. A guided tour into subcellular colocalization analysis in light microscopy. Journal of microscopy 224, 213–232 (2006).

73 Mirdita, M. et al. ColabFold: making protein folding accessible to all. Nature Methods 19, 679–682 (2022). 10.1038/s41592-022-01488-1

74 Pettersen, E. F. et al. UCSF Chimera—a visualization system for exploratory research and analysis. Journal of computational chemistry 25, 1605–1612 (2004).

75 Baker, C. et al. Identifying optimal capsid duplication length for the stability of reporter flaviviruses. Emerging Microbes & Infections 9, 2256–2265 (2020).

