## Supplement Information for "Mechanistic Insights into Dengue Virus Inhibition by a Clinical Trial Compound NITD-688"

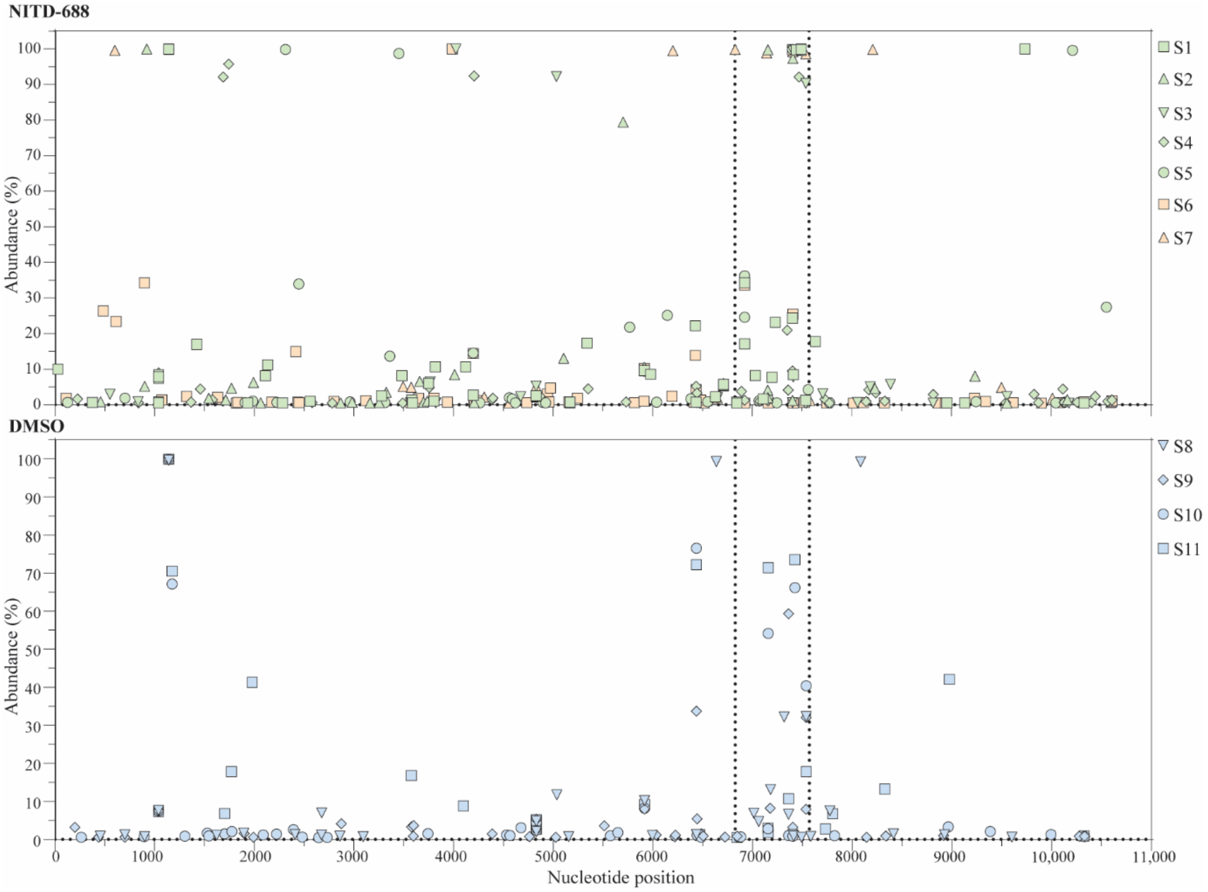


**Figure S1. Single nucleotide polymorphisms of the P15 viruses.** Mutations with frequency ≥0.5% identified in each selection by Illumina-based next-generation sequencing are shown. NS4B is shown as the region between two dashed lines. The genome positions refer to the infectious clone-derived DENV-2 NGC strain (GenBank access ID, AF038403).

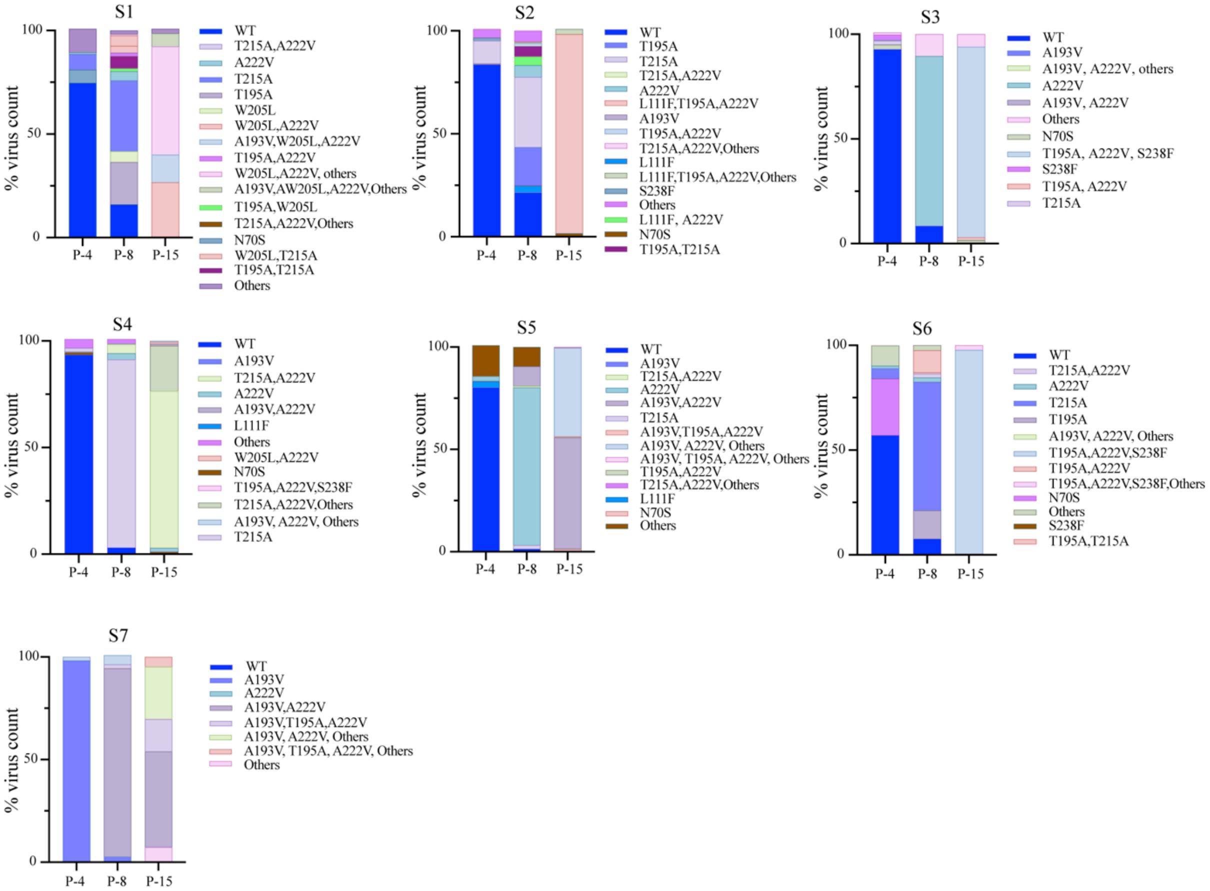


**Figure S2. PacBio sequencing of P4, P8 and P15 viruses.** The percentages of mutant viruses in each passage at P-4, P-8 and P-15 are shown. Mutant virus with abundance below 0.1% is indicated as “Others” in each panel.

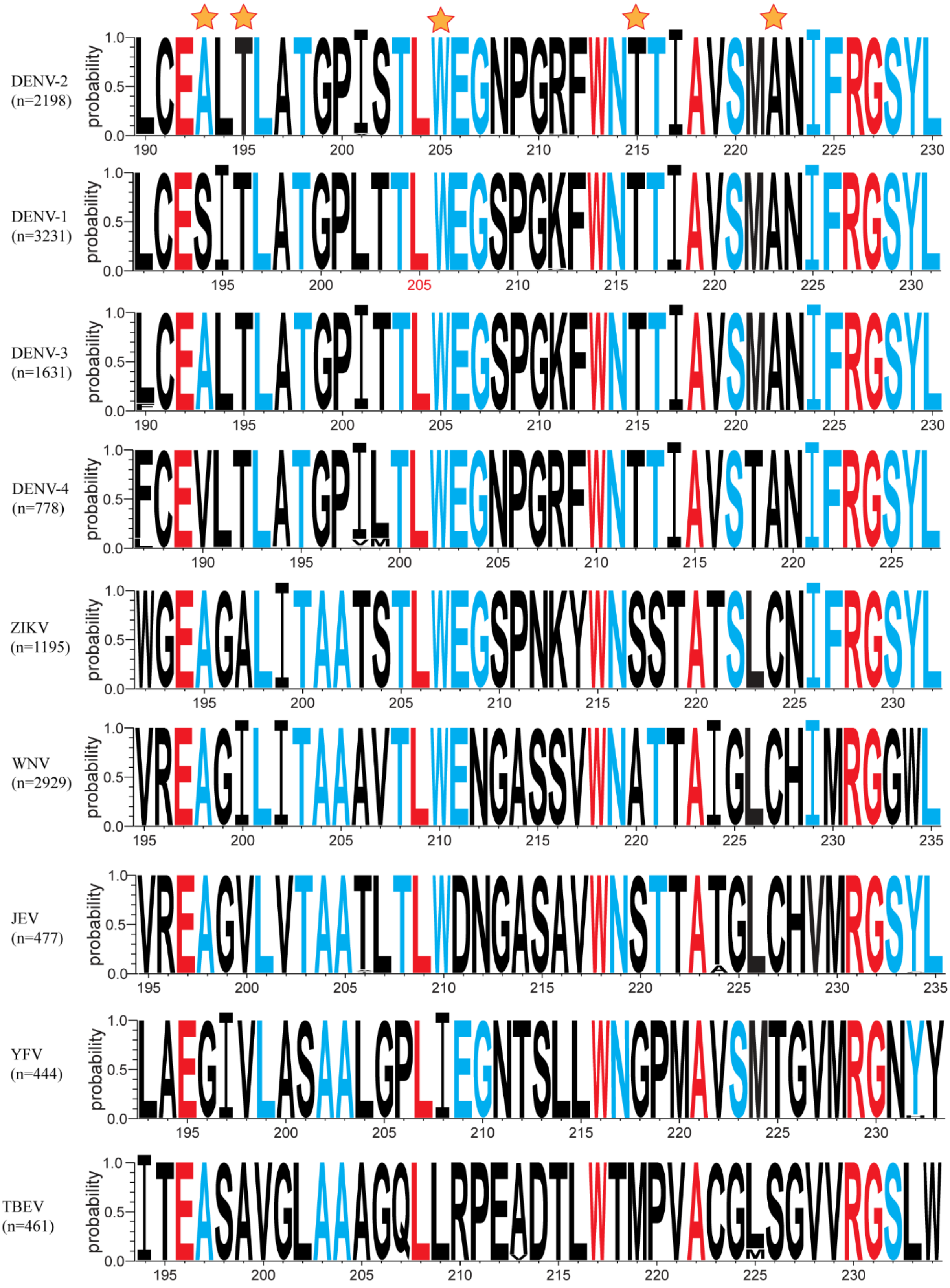


**Figure S3. Weblogo display of sequence alignment of flavivirus NS4B.** DENV-2 serves as a reference. Regions in other flaviviruses aligned with amino acids ranging from 190 to 230 of DENV-2 are shown. The overall height of the stack indicates the sequence conservation at that position in each flavivirus. The height of symbols within the stack indicates the relative frequency of each amino acid at that position. Stars show the mutations identified in NITD-688-resistant viruses. Identical and residuals with conservation higher than 50% among various flaviviruses are coded red and blue, respectively. n, the number of sequences each flavivirus used for alignment.

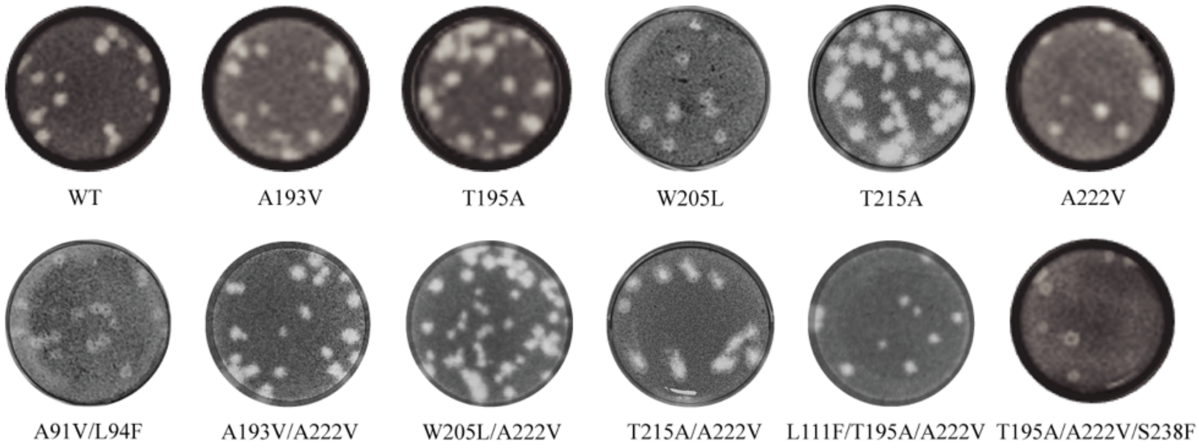


**Figure S4. Characterization of recombinant DENV-2 with NS4B mutations.** Plaque morphology developed on BHK-21 cells at day 5 post-infection.

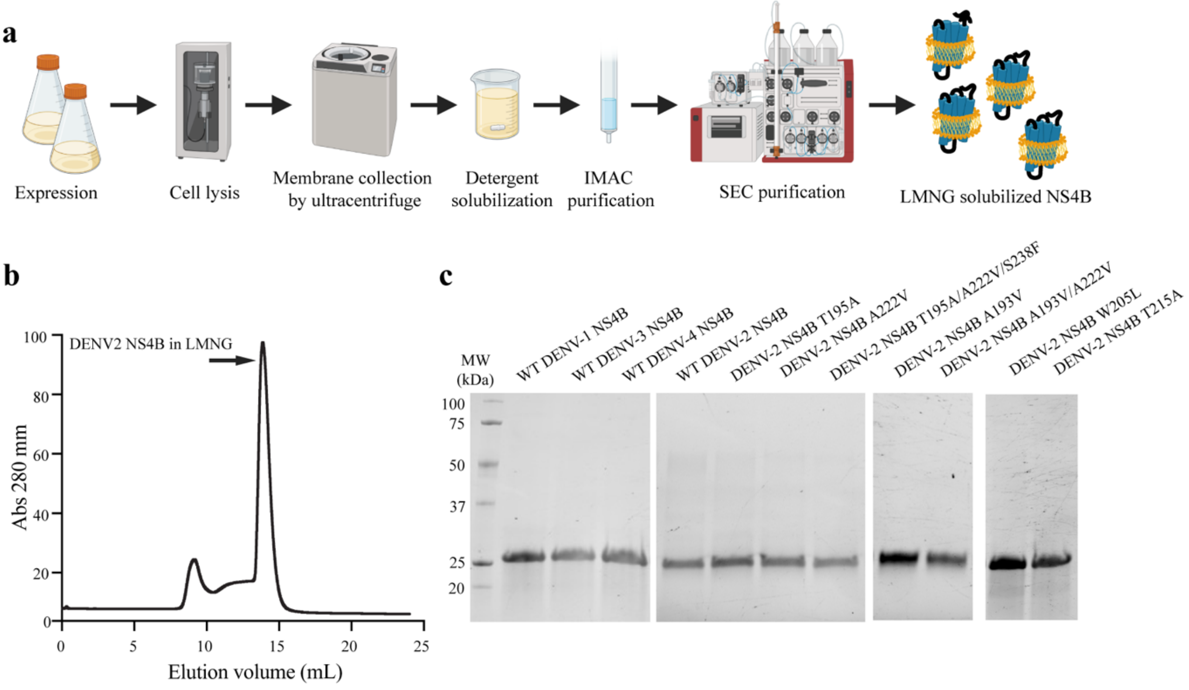


**Fig S5. Purification of recombinant NS4B proteins. a** Diagram shows the workflow for purifying recombinant NS4B. See details in Material and Methods. **b** Representative SEC profile of purified DENV-2 NS4B proteins. The arrow indicates the purified NS4B proteins. **c** SDS-PAGE analysis of purified WT DENV-1~4 and DENV-2 NS4B with indicated mutation(s).

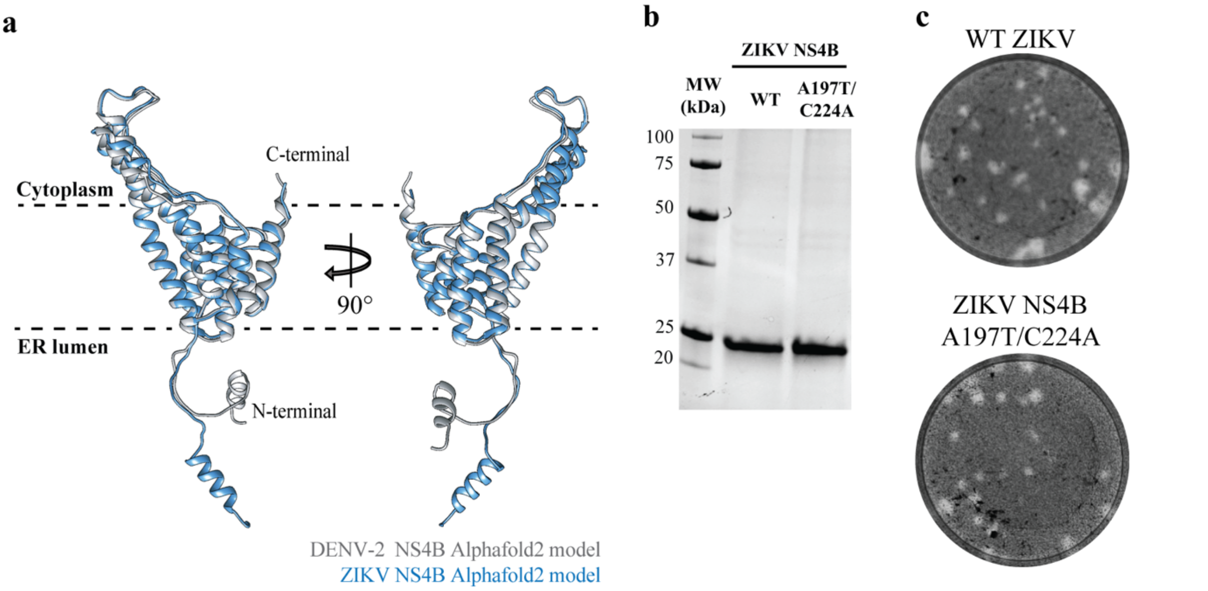


**Figure S6. Characterization of ZIKV NS4B. a** Alignment of DENV-2 (gray) and ZIKV (blue) NS4B structures. Both structures were predicted by Alphafold2. **b** SDS-PAGE gel analysis of purified WT and mutant A197T/C224A ZIKV NS4B proteins. (C) Plaque morphologies of recombinant WT ZIKV-Nluc and ZIKV-Nluc NS4B mutant A197T/C224A. Plaques were developed on VeroE6 cells at day 3 post-infection.

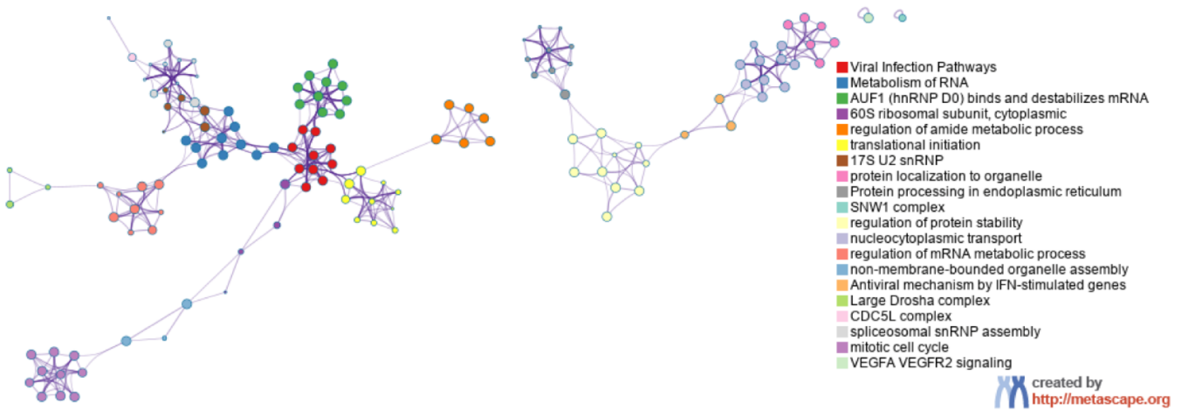


**Figure S7. Network of the enriched clusters of the common hits identified in DMSO and NITD-688 treated cells.** The plot was generated through a web tool Metascape. Each term is represented by a circle node, where its size is in proportion to the number of input genes that fall under each term. The color represents its cluster identity. The top 20 clusters are presented.

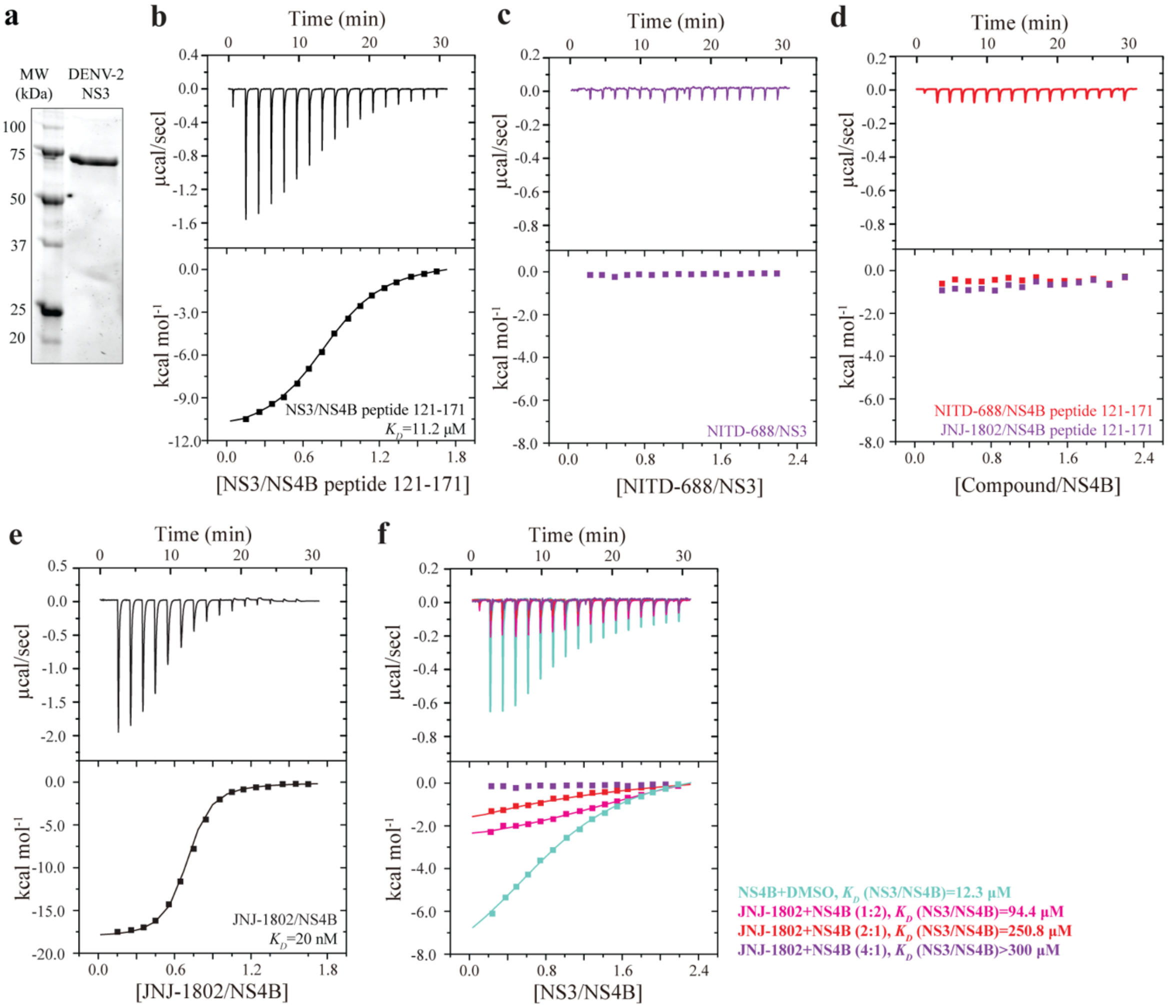


**Figure S8. ITC analysis of the binding between inhibitors and NS4B or NS3 proteins. a** SDS-PAGE analysis of purified DENV-2 NS3 proteins. **b** ITC analysis of DENV-2 NS3 binding to the cytosolic loop of NS4B (peptide 121-171). **c** ITC analysis of NITD-688 binding to DENV-2 NS3. **d** ITC analysis of NITD-688 or JNJ-1802 binding to the cytosolic loop of NS4B (peptide 121-171). **e** ITC analysis of JNJ-1802 binding to DENV-2 NS4B. **f** ITC analysis of DENV-2 NS4B binding to NS3 in the absence or presence of JNJ-1802. The estimated $K_D$ values of NS3/NS4B binding at each concentration of inhibitors are shown.

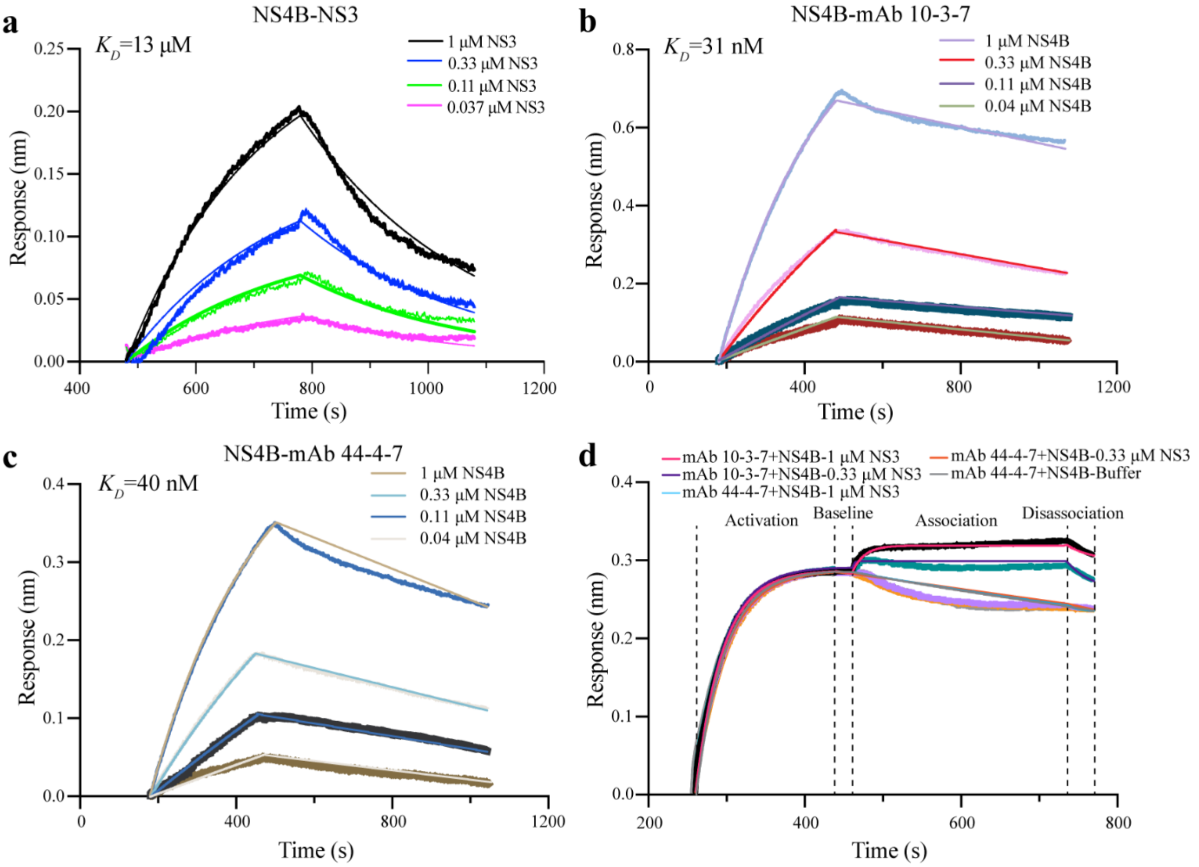


**Figure S9. Bio-Layer Interferometry (BLI) analysis of the binding of NS3 or monoclonal antibodies to NS4B proteins. a** Binding curves showing the association and dissociation of NS3 to biotinylated NS4B proteins immobilized on the biosensors at given concentrations of NS3 proteins. The estimated $K_D$ value is shown. **b** Binding curves showing the association and dissociation of DENV-2 NS4B to biotinylated mAb 10-3-7 immobilized on the biosensors at given concentrations of NS4B proteins. **c** Binding curves showing the association and dissociation of DENV-2 NS4B to biotinylated mAb 44-4-7 immobilized on the biosensors at given concentrations of NS4B proteins. **d** Epitope binning assay of mAb10-3-7, NS4B and NS3, or mAb 44-4-7, NS4B and NS3. Activation indicates the binding of NS4B proteins to biotinylated mAbs 10-3-7 or 44-4-7 immobilized on the biosensors. Association indicates the binding of NS3 to the assembled mAb/NS4B complexes. NS3 proteins associate with NS4B/mAb 10-3-7 complexes (no competition) and interfere with NS4B/mAb 44-4-7 binding (competition).

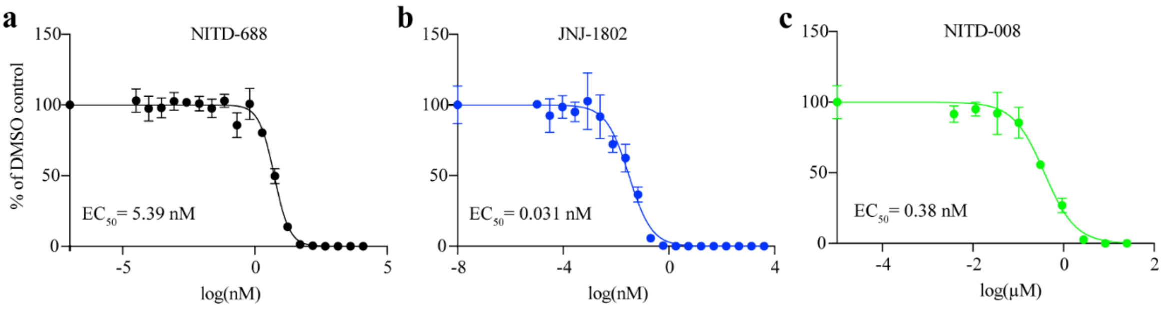


**Figure S10. Anti-DENV-2 activities of NITD-688, JNJ-1802, and NITD-688 on Huh7 cells.** Huh7 cells were infected with the NGC-Nluc and treated with various concentrations of inhibitors for 48 hours. **a** Dose-response curve and $EC_{50}$ of NITD-688 against NGC-Nluc. **b** Dose-response curve and $EC_{50}$ of JNJ-1802 against NGC-Nluc. **c** Dose-response curve and $EC_{50}$ of NITD-688 against NGC-Nluc.

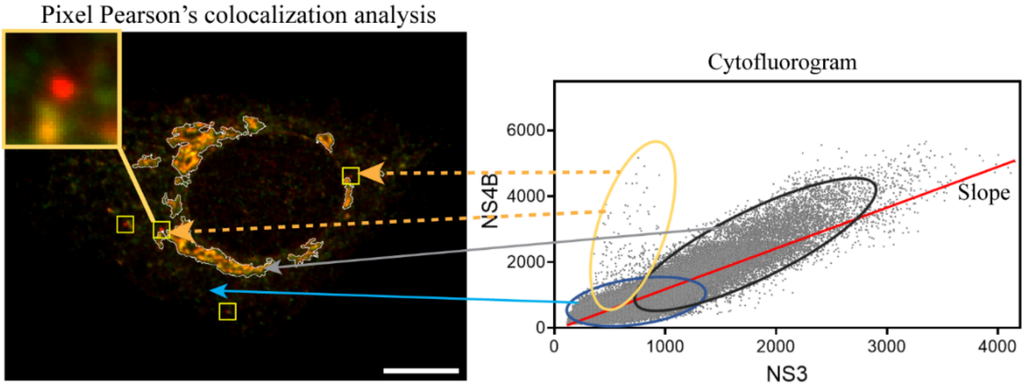


**Figure S11. Colocalization analysis of NS4B and NS3**. Representative example showing cytofluorogram analysis. The intensity of NS4B or NS3 at each pixel of images was measured and plotted in a two-dimensional graph. A linear regression model was fitted to the scattered dots from a single cell. The slope was calculated from the linear regression model. Pearson's correlation coefficient was also calculated.

Table S1. Amino acid changes in P15 viruses identified by next-generation sequencing.

| Genome Position | Mutations (viral proteins_position) | $NITD-688 selection (frequency, %) | | | | | | $DMSO selection (frequency, %) | | | | |
|---|---|---|---|---|---|---|---|---|---|---|---|---|
| | | S1 | S2 | S3 | S4 | S5 | S6 | S7 | S8 | S9 | S10 | S11 |
| 614 | prM_N59S | | | | | | 23.4 | | | | | |
| 898 | prM_I154V | | | | | | 34.3 | | | | | |
| 1142 | E_T69I | 99.9 | 99.9 | 99.8 | 99.8 | 99.9 | 99.9 | 99.8 | 99.7 | 99.9 | 99.9 | 99.9 |
| 2135 | E_Q400R | 11.2 | | | | | | | | | | |
| 2447 | NS1_K9R | | | | | 33.9 | | | | | | |
| 3575 | NS2A I33T | | | | | | | 4.9 | | | | 16.8 |
| 3818 | NS2A S114T | 10.7 | | | | | | | | | | |
| 3988 | NS2A_I171V | | | | | | 99.9 | | | | | |
| 4024 | NS2A_T183S | | | 99.9 | | | | | | | | |
| 4121 | NS2A N215S | 10.7 | | | | | | | | | | |
| 4199 | NS2B K23R | 2.7 | | | | 14.5 | 14.4 | | | | | |
| 4207 | NS2B_I26V | | 0.6 | | 92.3 | | 1.0 | | | | | |
| 5036 | NS3_I172T | | | | | | | | 11.7 | | | |
| 6203 | NS3_R561K | | | | | | | 99.4 | | | | |
| 6427 | NS4A T18A | 22.2 | | | | | 13.9 | | | | | |
| 6437 | NS4A_A21V | | | | 5.2 | | | | | 33.7 | 76.5 | 72.2 |
| 6922 | NS4B_R33S | 17.1 | | | | | | | | | | |
| 6922 | NS4B_R33C | 34.3 | | | 1.3 | 11.6 | 0.5 | | | | | |
| 6923 | NS4B_R33Y/H | | | | | 24.6 | | | | | | |
| 6923 | NS4B_R33H | | | | | | 33.6 | | | | | |
| 7156 | NS4B_L111F | | 99.7 | 1.1 | 1.7 | | | | | | 54.1 | 71.4 |
| 7349 | NS4B_V175A | | | | 21.0 | | | | | | | |
| 7361 | NS4B_T179I | | | | 4.2 | | | | 6.6 | 59.3 | 1.0 | 10.7 |
| 7403 | NS4B_A193V | 24.4 | 1.2 | 0.6 | 9.5 | 99.8 | 99.7 | 0.9 | 1.3 | 3.2 | 0.8 | |
| 7408 | NS4B_T195A | | 97.3 | 99.3 | | | 25.5 | 99.2 | | | | |
| 7439 | NS4B_W205L | 99.8 | | | | | | | | | | |
| 7468 | NS4B_T215A | | | | 92.1 | | | | | | | |
| 7490 | NS4B_A222V | 99.8 | 99.5 | 99.9 | 99.9 | 99.8 | 99.7 | 99.8 | 0.6 | | | |
| 7538 | NS4B_S238F | | | 90.3 | 0.5 | | | 98.5 | 32.3 | 32.1 | 40.4 | 17.8 |
| 7624 | NS5_A19T | | | | 17.7 | | | | | | | |
| 9737 | NS5_V723A | 99.9 | | | | | | | | | | |

$Amino acid substitutions with ≥10% frequency in any of the selections are shown. For comparison, the frequency (>0.5%) of the same mutations in other selections is indicated.

Table S2. $EC_{50}$ values of JNJ-1802 against DENV-2 WT and NS4B mutants.

| Virus | $EC_{50}$ (nM) | Fold change* |
|---|---|---|
| DENV-2 WT | 0.019 ± 0.004 | 1.0 |
| DENV-2 V91A/L94F | 55.89 ± 5.83 | 2941.6 |
| DENV-2 L111F/T195A/A222V | 0.044 ± 0.011 | 2.3 |
| DENV-2 T195A/A222V/S238F | 0.039 ± 0.003 | 2.1 |

*Fold change was calculated by comparing the $EC_{50}$ against each mutant to that against WT.

Table S3. List of primers used in this study.

| Primer Name | Sequence (5'-3') | Notes |
|---|---|---|
| DENV-2 L111F-F | CTCACAGCAGCTTTTTCTTACTGGTAGC | For constructing DENV-2 NS4B mutation L111F |
| DENV-2 L111F-R | GTAAGAAAAAAGCTGCTGTGAGAGTTATGGG | For constructing DENV-2 NS4B mutation L111F |
| DENV-2 A193V-F | GTGTGAGGTGTTAACCTTAGCGACCGGGC | For constructing DENV-2 NS4B mutation A193V |
| DENV-2 A193V-R | CGCTAAGGTTAACACCTCACACAGAGCCC | For constructing DENV-2 NS4B mutation A193V |
| DENV-2 T195A-F | GGCTTTAGCCTTAGCGACCGGGCCTATCTCC | For constructing DENV-2 NS4B mutation T195A |
| DENV-2 T195A-R | GCTAAGGCTAAAGCCTCACACAGAGCCC | For constructing DENV-2 NS4B mutation T195A |
| DENV-2 W205L-F | CCACATTGCTGGAAGGAAATCCAGGG | For constructing DENV-2 NS4B mutation W205L |
| DENV-2 W205L-R | TCCTTCCAGCAATGTGGAGATAGGC | For constructing DENV-2 NS4B mutation W205L |
| DENV-2 T215A-F | GAGGTTTTGGAACGCAACCATTGCAGTGTC | For constructing DENV-2 NS4B mutation T215A |
| DENV-2 T215A-R | CTGCAATGGTTGCGTTCCAAAACCTCCCTGG | For constructing DENV-2 NS4B mutation T215A |
| DENV-2 A222V-F | CAGTGTCAATGGTTAACATTTTTAGAGGG | For constructing DENV-2 NS4B mutation A222V |
| DENV-2 A222V-R | AAATGTTAACCATTGACACTGCAATGGT | For constructing DENV-2 NS4B mutation A222V |
| DENV-2 S238F-F | CTTCTCTTTTTCATCATGAAGAACACAACC | For constructing DENV-2 NS4B mutation S238F |
| DENV-2 S238-R | GTTCTTCATGATGAAAAAGAGAAGTCCAGC | For constructing DENV-2 NS4B mutation S238F |
| DENV-2 V91A/L94F-F | ATCGGAGCGCCCCTTTTTGCCATTGGATGC TACTCACAAG | For constructing ZIKV NS4B mutation V91A/L94F |
| DENV-2 V91A/L94F-R | CCAATGGCAAAAAGGGGCGCTCCGATGTCC ATCTTTGAC | For constructing ZIKV NS4B mutation V91A/L94F |
| DENV-2 NGC-F | CGTCGAGAGAAATATGGTCACACC | For RT-qPCR |
| DENV-2 NGC-R | CCACAATAGTATGACCAGCCT | For RT-qPCR |
| DENV-2 XhoI-F | GCGGCTAGAGGATACATCTCAAC | For constructing infection clones and RT-PCR |
| DENV-2 NruI-R | CAGTTTTGCTGAGCCTCGCGACACAGCGTG | For constructing infection clones and RT-PCR |
| ZIKV T197A-F | GGAGGCTGGGACACTGATCACAGCAGCAACC | For constructing ZIKV NS4B mutation T197A |
| ZIKV T197A-R | GCTGTGATCAGTGTCCCAGCCTCCCCCC | For constructing ZIKV NS4B mutation T197A |
| ZIKV C224A-F | CACTGGCCAACATCTTTAGAGG | For constructing ZIKV NS4B mutation C224A |
| ZIKV C224A-R | CTAAAGATGTTGGCCAGTGAGGTGGCTG | For generation ZIKV NS4B mutation C224A |
| ZIKV NheI-F | CCCCGTGAGAGCATGCTGCTAGCCC | For constructing ZIKV infectious clones |
| ZIKV AscI-R | CCTTGAGGGCGCGGCGCGCCTCC | For constructing ZIKV infectious clones |
| NS4B-Pac-F | AAGGAGTTTGCAGCTGG | For PacBio sequencing |
| NS4B-Pac-R | CTCTCCTATGTTACCGGTTC | For PacBio sequencing |
| pCAG-2K-F | GGAGGTACCATGACACCCCAAGATAACCAATT | For construction pCAG-2K-NS4B WT and TM |
| pCAG-NS4B-R | GAGCCCTCGAGCCTTCTCGTGTTGGTTGTG | For construction pCAG-2K-NS4B WT and TM |
| pCAG-2B-F | GGAGGTACCATGAGCTGGCCACTAAATG | For construction pCAG-2B-NS3 |
| pCAG-NS3-R | GCCAAGCTTTTACTTTCTTCCAGCTGC | For construction pCAG-2B-NS3 |
| pEF1a-NS2B-F | CGGTGATATCAAAGATCTTACAGCTAGCGCCA CCATGAGCTGGCCACTAAATGAGGCTAT | For constructing pEF1a-NS2B-NS3 |
| pEF1a-NS3-R | ATGTATCTTATCATGTCTGGGCGGCCGCCTAC TTTCTTCCAGCTGCAAACTCC | For constructing pEF1a-NS2B-NS3 |
| pEF1a-2K-F | GGATCCCGTACGCCTAGGGGGAATTCGCCAC CATGACACCCCAAGATAACCAATTGAC | For constructing pEF1a-2K-NS4B |
| pEF1a-NS4B-R | CTCTTCTGAGATGAGTTTCTGCTCGAGCCTTC TCGTGTTGGTTGTGTTCTTCATG | For constructing pEF1a-2K-NS4B and pEF1a-NS4A-2K-NS4B |
| pEF1a-NS4A-F | GGATCCCGTACGCCTAGGGGGAATTCGCCA CCATGTCCCTGACCCTGAACCTAATCACAG | For constructing pEF1a-NS4A-2K-NS4B |

### Table S4 host factors identified from IP-MS.

| Accession | # Unique Peptides | Score Sequest HT: Sequest HT | # Peptides (by Search Engine): Sequest HT | [#]Abundance Ratio: (IV) / (III) | [#]Abundance Ratio Adj. P-Value: (NITD-688) / (DMSO) | [#]Abundances (Grouped): DMSO | [#]Abundances (Grouped): NITD-688 |
|---|---|---|---|---|---|---|---|
| P04350 | 4 | 1911.17 | 28 | | | | |
| Q13501 | 2 | 22.89 | 2 | | | | |
| Q9UI12 | 2 | 12.57 | 2 | | | | |
| P22087 | 2 | 9.8 | 2 | | | | |
| P35579 | 1 | 9.68 | 2 | | | | |
| Q8N0X7 | 2 | 6.95 | 2 | | | | |
| Q9Y5M8 | 3 | 6.52 | 3 | | | | |
| Q9Y536 | 1 | 6.25 | 2 | | | | |
| Q03252 | 1 | 5.51 | 2 | | | | |
| P04637 | 2 | 3.48 | 2 | | | | |
| Q9C0E8 | 2 | 2.26 | 2 | | | | |
| Q6PKG0 | 2 | 2.25 | 2 | | | | |
| Q96S52 | 2 | 2.21 | 2 | | | | |
| P21796 | 2 | 2.09 | 2 | | | | |
| Q8IY63 | 1 | 1.79 | 2 | | | | |
| P25205 | 2 | 1.7 | 2 | | | | |
| Q92896 | 2 | 1.69 | 2 | | | | |
| Q6YHU6 | 3 | 1.66 | 3 | | | | |
| Q13885 | 1 | 2041.33 | 30 | | | | |
| P52597 | 1 | 50.03 | 4 | | | | |
| Q7RTS7 | 1 | 30.73 | 4 | | | | |
| P21964 | 2 | 11.95 | 2 | | | | |
| Q9UID3 | 3 | 11.46 | 3 | | | | |
| P04181 | 2 | 8.5 | 2 | | | | |
| P46736 | 2 | 7.75 | 2 | | | | |
| Q96S59 | 1 | 4.62 | 2 | | | | |
| Q9BRP8 | 2 | 4.28 | 2 | | | | |
| O43681 | 2 | 3.64 | 2 | | | | |
| O00425 | 1 | 2.51 | 3 | | | | |
| Q8N283 | 1 | 0 | 2 | | | | |
| O75390 | 2 | 0 | 2 | | | | |
| Q9Y520 | 2 | 0 | 2 | | | | |
| Q9UQE7 | 2 | 0 | 2 | | | | |
| P31942 | 1 | 0 | 2 | | | | |
| Q14315 | 1 | 48.7 | 6 | | | | |
| Q15645 | 2 | 9.25 | 2 | | | | |
| O43150 | 2 | 7.03 | 2 | | | | |
| P35749 | 1 | 3.02 | 2 | | | | |
| P36578 | 6 | 24 | 6 | 0.763 | 0.999694499 | 1142486.948 | 871456 |
| P50991 | 22 | 192.34 | 23 | 1.021 | 0.999694499 | 14475596.72 | 14775709.88 |
| P62753 | 5 | 39.27 | 5 | 1.14 | 0.999694499 | 1490204.9 | 1698660.553 |
| Q14011 | 11 | 233.81 | 11 | 0.972 | 0.999694499 | 14395455.54 | 13991744.73 |
| Q3ZCQ8 | 5 | 18.25 | 5 | 1.042 | 0.999694499 | 804679.4241 | 838324.7535 |
| Q8WU90 | 2 | 1.91 | 2 | 1.225 | 0.999694499 | 306776.9111 | 375686.8338 |
| Q8WY22 | 3 | 11.15 | 3 | 1.01 | 0.999694499 | 636315.5307 | 642533.6055 |
| Q96I25 | 20 | 101.12 | 20 | 1.077 | 0.999694499 | 7607806.546 | 8190800.724 |
| Q96RL7 | 2 | 1.68 | 2 | 1.104 | 0.999694499 | 2618057.432 | 2890128.015 |
| Q9UBB6 | 12 | 86.8 | 12 | 1.073 | 0.999694499 | 2733696.698 | 2932663.33 |
| Q9UJV9 | 12 | 69.4 | 12 | 1.325 | 0.999694499 | 6278636.54 | 8320852.846 |
| Q9UNM6 | 20 | 246.3 | 20 | 0.998 | 0.999694499 | 15896450.53 | 15864850.67 |
| P46379 | 32 | 569.41 | 32 | 1.024 | 0.997236921 | 46252541.96 | 47347803.58 |
| P61978 | 16 | 231.86 | 16 | 1.087 | 0.997236921 | 9797628.938 | 10654891.73 |
| P62805 | 5 | 53.58 | 5 | 1.068 | 0.997236921 | 3088846.099 | 3298161.25 |
| Q12906 | 9 | 55.75 | 9 | 1.172 | 0.997236921 | 2376618.36 | 2784585.676 |
| Q15046 | 6 | 37.85 | 6 | 1.043 | 0.997236921 | 661414.8797 | 689649.7066 |
| Q92890 | 16 | 143.75 | 16 | 1.107 | 0.997236921 | 11469245.61 | 12694918.18 |
| Q96A65 | 4 | 19.65 | 4 | 1.108 | 0.997236921 | 213726.0929 | 236820.2755 |
| Q9NSD9 | 3 | 7.06 | 3 | 1.201 | 0.997236921 | 144107.9537 | 173062.7325 |
| Q9NYZ3 | 2 | 9.06 | 2 | 1.029 | 0.997236921 | 68399.82413 | 70393.7946 |
| Q9P035 | 3 | 17.23 | 3 | 0.977 | 0.997236921 | 463507.4046 | 452858.9211 |
| Q9ULV4 | 6 | 15.05 | 6 | 1.079 | 0.997236921 | 785853.4859 | 847591.1382 |
| Q9Y5B9 | 13 | 67.73 | 13 | 1.003 | 0.997236921 | 2675096.319 | 2682787.852 |
| Q13155 | 3 | 7.16 | 3 | 0.717 | 0.996743093 | 681386.1297 | 488248.4386 |
| Q15208 | 28 | 673.49 | 35 | 1.122 | 0.990986166 | 53771994.48 | 60348999.74 |
| Q13435 | 32 | 316.27 | 32 | 1.134 | 0.988789719 | 25503290.44 | 28914219.7 |
| A0A0C4DH68 | 1 | 577.18 | 2 | 0.923 | 0.98631319 | 4793929.039 | 4422627.894 |
| A1L0T0 | 5 | 37.23 | 5 | 0.93 | 0.98631319 | 1152066.752 | 1071847.563 |
| A2NJV5 | 4 | 6199.78 | 5 | 1.02 | 0.98631319 | 10363149994 | 10572788663 |
| A5YKK6 | 11 | 40.75 | 11 | 0.734 | 0.98631319 | 1057373.688 | 775639.1335 |
| A6NDU8 | 2 | 2.04 | 2 | 1.283 | 0.98631319 | 170744.5505 | 219083.5062 |
| A6NMY6 | 2 | 9.35 | 2 | 0.862 | 0.98631319 | 70530.93669 | 60762.47737 |
| Q71U36 | 1 | 2457.02 | 38 | 0.956 | 0.98631319 | 4868419.552 | 4656568.575 |
| NS4B-Tagged | 15 | 2705.5 | 15 | 1.25 | 0.98631319 | 219721767.7 | 274548197.4 |
| O00165 | 2 | 18.21 | 2 | 1.267 | 0.98631319 | 362751.7661 | 459537.7727 |
| O00231 | 16 | 202.09 | 16 | 0.965 | 0.98631319 | 15312857.46 | 14778244.85 |
| O00232 | 14 | 116.39 | 14 | 1.001 | 0.98631319 | 7631824.157 | 7636377.423 |
| O00303 | 7 | 22.7 | 7 | 0.925 | 0.98631319 | 956261.1405 | 884911.3655 |
| Q9BVA1 | 1 | 2052.41 | 31 | 0.932 | 0.98631319 | 88461.45909 | 82442.5834 |
| O00410 | 38 | 485.04 | 38 | 0.885 | 0.98631319 | 28497550.1 | 25225200.8 |
| O00471 | 2 | 2.56 | 2 | 1.012 | 0.98631319 | 131313.091 | 132940.5861 |
| O00487 | 8 | 109.5 | 8 | 0.697 | 0.98631319 | 6780880.513 | 4724129.297 |
| O00571 | 23 | 198.74 | 24 | 0.944 | 0.98631319 | 9781439.268 | 9237562.288 |
| O14545 | 17 | 140.32 | 17 | 0.892 | 0.98631319 | 9952289.456 | 8879125.72 |
| O14654 | 16 | 110.5 | 16 | 1.032 | 0.98631319 | 4092378.171 | 4222929.337 |
| O14744 | 58 | 8036.2 | 58 | 1.054 | 0.98631319 | 1478989985 | 1558906776 |
| O14787 | 4 | 63.4 | 10 | 1.128 | 0.98631319 | 233764.5848 | 263725.4027 |
| O14893 | 4 | 3.78 | 4 | 1.766 | 0.98631319 | 820623.2891 | 1449537.888 |
| O14980 | 50 | 1031.25 | 50 | 1.014 | 0.98631319 | 64546642.76 | 65435898.83 |
| O14981 | 3 | 0 | 3 | 0.887 | 0.98631319 | 488072.2846 | 432829.946 |
| O15042 | 25 | 119.06 | 25 | 0.862 | 0.98631319 | 9265907.777 | 7988343.443 |
| O15269 | 7 | 47.49 | 7 | 0.878 | 0.98631319 | 2465434.577 | 2165643.592 |
| O15294 | 8 | 44.39 | 8 | 1.008 | 0.98631319 | 61433305.98 | 61925084.27 |
| O15371 | 10 | 90.23 | 10 | 0.904 | 0.98631319 | 1791516.239 | 1618898.032 |
| O15372 | 6 | 57 | 6 | 0.889 | 0.98631319 | 1881467.497 | 1672243.374 |
| O15397 | 17 | 230.4 | 18 | 0.835 | 0.98631319 | 12830562.17 | 10710545.92 |
| O43143 | 30 | 345.89 | 30 | 0.873 | 0.98631319 | 38360475.68 | 33488980.93 |
| O43156 | 4 | 25.86 | 4 | 0.893 | 0.98631319 | 499760.7295 | 446103.4454 |
| Q9BUF5 | 3 | 693.09 | 13 | 3.27 | 0.98631319 | 52923.46754 | 173069.834 |
| O43167 | 2 | 0 | 2 | 0.362 | 0.98631319 | 197172.5222 | 71368.07117 |
| O43175 | 8 | 70.33 | 8 | 1.252 | 0.98631319 | 842711.4569 | 1055412.353 |
| O43242 | 24 | 198.95 | 24 | 1.088 | 0.98631319 | 13112647.31 | 14269452.21 |
| O43290 | 8 | 39.85 | 8 | 1.186 | 0.98631319 | 1466924.595 | 1740280.484 |
| O43318 | 16 | 128.01 | 16 | 1.054 | 0.98631319 | 3904923.369 | 4117333.038 |
| O43390 | 4 | 27.26 | 4 | 1.632 | 0.98631319 | 582885.964 | 950989.1731 |
| O43395 | 4 | 7.35 | 4 | 1.081 | 0.98631319 | 488789.235 | 528330.4361 |
| O43505 | 2 | 1.62 | 2 | 0.92 | 0.98631319 | 468715.0301 | 431321.7958 |
| O43592 | 22 | 272.58 | 22 | 1.102 | 0.98631319 | 17252501.89 | 19004364.35 |
| O43615 | 2 | 4.69 | 2 | 1.139 | 0.98631319 | 170427.8018 | 194198.5523 |
| O43660 | 5 | 16.19 | 5 | 1.152 | 0.98631319 | 638267.3965 | 735175.7969 |
| O43719 | 10 | 60.63 | 10 | 1.638 | 0.98631319 | 2186079.469 | 3580798.911 |
| O43809 | 4 | 33.41 | 4 | 1.338 | 0.98631319 | 774841.5658 | 1037053.285 |
| O43933 | 2 | 3.48 | 2 | 0.868 | 0.98631319 | 802161.2586 | 696389.8892 |
| P63261 | 1 | 407.82 | 24 | 7.761 | 0.98631319 | 84736.88564 | 657628.2339 |
| O60613 | 3 | 12.05 | 3 | 0.626 | 0.98631319 | 279694.1967 | 174999.7104 |

| | | | | | | | |
|---|---|---|---|---|---|---|---|
| O60678 | 3 | 5.2 | 3 | 1.084 | 0.98631319 | 764387.4896 | 828355.6964 |
| O60825 | 3 | 3.87 | 3 | 0.937 | 0.98631319 | 170641.8205 | 159949.1082 |
| O60841 | 5 | 16.2 | 5 | 0.861 | 0.98631319 | 707849.758 | 609352.189 |
| O60884 | 13 | 120 | 13 | 1.236 | 0.98631319 | 8746331.174 | 10809588.44 |
| O75122 | 6 | 43.89 | 6 | 0.719 | 0.98631319 | 521236.0665 | 374759.8972 |
| O75155 | 10 | 49.38 | 11 | 0.983 | 0.98631319 | 2072669.699 | 2037991.586 |
| O75306 | 2 | 1.8 | 2 | 0.948 | 0.98631319 | 248989.5133 | 236164.7266 |
| O75533 | 64 | 851.86 | 64 | 0.997 | 0.98631319 | 54848880.48 | 54671687.5 |
| O75643 | 42 | 294.22 | 42 | 1.078 | 0.98631319 | 11372742.7 | 12259018.35 |
| O75688 | 17 | 146.68 | 17 | 0.761 | 0.98631319 | 12448154.58 | 9471761.303 |
| O75821 | 5 | 41.85 | 5 | 1.032 | 0.98631319 | 1596535.226 | 1647051.401 |
| O75940 | 4 | 25.25 | 4 | 0.938 | 0.98631319 | 581328.4992 | 545263.9732 |
| O75955 | 5 | 9.25 | 5 | 0.938 | 0.98631319 | 210942.9911 | 197779.3281 |
| O76071 | 3 | 1.61 | 3 | 1.578 | 0.98631319 | 309685.5252 | 488778.9429 |
| O94822 | 17 | 73.96 | 17 | 0.948 | 0.98631319 | 2329951.835 | 2209024.698 |
| O94829 | 4 | 2.38 | 4 | 1.737 | 0.98631319 | 292830.7937 | 508774.6395 |
| O94888 | 2 | 8.81 | 2 | 0.953 | 0.98631319 | 252459.9555 | 240639.2093 |
| O94905 | 7 | 45.16 | 7 | 0.619 | 0.98631319 | 1679861.993 | 1039273.86 |
| O94906 | 30 | 231.7 | 30 | 1.027 | 0.98631319 | 11612274.78 | 11922370.2 |
| O95071 | 9 | 11.34 | 9 | 2.943 | 0.98631319 | 812153.1434 | 2390468.246 |
| O95197 | 3 | 17.56 | 3 | 0.929 | 0.98631319 | 1832236.065 | 1702533.35 |
| O95218 | 3 | 10.68 | 3 | 1.072 | 0.98631319 | 415405.2045 | 445418.3392 |
| O95347 | 15 | 51.75 | 15 | 0.897 | 0.98631319 | 2586869.808 | 2319586.911 |
| O95373 | 20 | 217.33 | 21 | 1.008 | 0.98631319 | 14596579.31 | 14714391.81 |
| O95400 | 2 | 18.7 | 2 | 0.893 | 0.98631319 | 236997.3009 | 211524.2982 |
| O95816 | 8 | 52.34 | 8 | 1.007 | 0.98631319 | 2291992.044 | 2307680.547 |
| O95831 | 3 | 32.16 | 3 | 1.365 | 0.98631319 | 643983.1063 | 878905.1083 |
| O95881 | 8 | 117.13 | 8 | 1.031 | 0.98631319 | 10258091.75 | 10580886.9 |
| O96005 | 2 | 2.33 | 2 | 0.881 | 0.98631319 | 231737.3042 | 204196.1719 |
| P00761 | 8 | 2400.24 | 8 | 0.981 | 0.98631319 | 3110615901 | 3052994264 |
| P00846 | 2 | 5.13 | 2 | 0.967 | 0.98631319 | 186208.1599 | 180149.9965 |
| P01012 | 2 | 6.03 | 2 | 0.185 | 0.98631319 | 676151.6787 | 124913.438 |
| P01859 | 2 | 91.56 | 2 | 1.056 | 0.98631319 | 47954924.62 | 50647271.22 |
| P02533 | 8 | 423.81 | 28 | 1.008 | 0.98631319 | 5899106.789 | 5946216.744 |
| P02538 | 4 | 476.48 | 35 | 1.026 | 0.98631319 | 21868997.39 | 22439181.51 |
| P02769 | 17 | 171.15 | 17 | 1.033 | 0.98631319 | 10451966.37 | 10801426.41 |
| P04259 | 2 | 490.56 | 33 | 0.804 | 0.98631319 | 669300.6538 | 538033.479 |
| P04264 | 49 | 2282.87 | 56 | 0.953 | 0.98631319 | 245607049.3 | 234002664.4 |
| P04406 | 3 | 18.19 | 3 | 0.67 | 0.98631319 | 585834.0221 | 392234.6719 |
| P04843 | 23 | 239.26 | 23 | 0.897 | 0.98631319 | 13744055.23 | 12328512.4 |
| P04844 | 5 | 29.14 | 5 | 0.844 | 0.98631319 | 782771.6713 | 660768.25 |
| P05023 | 13 | 102.18 | 13 | 1.009 | 0.98631319 | 3363686.468 | 3394958.709 |
| P05141 | 4 | 110.88 | 12 | 0.879 | 0.98631319 | 19618171.42 | 17252753.06 |
| P05198 | 10 | 65.86 | 10 | 1.028 | 0.98631319 | 3170225.828 | 3257910.093 |
| P05388 | 16 | 265.01 | 16 | 0.927 | 0.98631319 | 16370764.54 | 15172674.8 |
| P06576 | 13 | 221.02 | 13 | 0.945 | 0.98631319 | 9231348.068 | 8723402.751 |
| P06730 | 5 | 37.06 | 5 | 0.816 | 0.98631319 | 2144482.113 | 1750078.268 |
| P06748 | 6 | 49.29 | 6 | 1.017 | 0.98631319 | 3966638.719 | 4034190.812 |
| P07437 | 6 | 2715.61 | 35 | 1.059 | 0.98631319 | 47404223.97 | 50218307.03 |
| P07814 | 40 | 344.43 | 40 | 0.893 | 0.98631319 | 13466592.7 | 12020404.99 |
| P07900 | 9 | 158.85 | 20 | 1.158 | 0.98631319 | 3098924.024 | 3588805.666 |
| P07910 | 10 | 66.21 | 10 | 0.79 | 0.98631319 | 5158332.657 | 4077645.597 |
| P08238 | 12 | 212.63 | 25 | 0.948 | 0.98631319 | 15383022.33 | 14577576.74 |
| P08579 | 4 | 72.31 | 6 | 1.013 | 0.98631319 | 9536386.083 | 9661985.869 |
| P08670 | 9 | 31.22 | 10 | 0.793 | 0.98631319 | 737505.1047 | 584829.1837 |
| P08708 | 7 | 70.68 | 7 | 1.071 | 0.98631319 | 6408282.195 | 6861477.426 |
| P08779 | 14 | 545.24 | 29 | 0.984 | 0.98631319 | 20392382.74 | 20072879.48 |
| P08865 | 4 | 16.62 | 4 | 0.897 | 0.98631319 | 1020898.728 | 916226.3202 |
| P09012 | 2 | 19.41 | 4 | 0.976 | 0.98631319 | 185496.1762 | 181012.66 |
| P09651 | 12 | 254.34 | 13 | 1.002 | 0.98631319 | 8461614.666 | 8476801.886 |
| P09661 | 8 | 65.65 | 8 | 0.846 | 0.98631319 | 3948454.901 | 3340362.911 |
| P09874 | 32 | 255.97 | 32 | 0.911 | 0.98631319 | 18141541.79 | 16519405.3 |
| P0CG47 | 1 | 3501.79 | 17 | 0.914 | 0.98631319 | 305617.7044 | 279402.9688 |
| P0DMV9 | 44 | 1958.55 | 51 | 1.025 | 0.98631319 | 216083989.1 | 221578307.5 |
| P10599 | 6 | 150.65 | 6 | 0.942 | 0.98631319 | 19104379.84 | 18003973.12 |
| P10644 | 8 | 45.92 | 8 | 0.993 | 0.98631319 | 2239031.193 | 2222747.381 |
| P10809 | 6 | 15.9 | 6 | 1.127 | 0.98631319 | 932354.3639 | 1050331.502 |
| P11021 | 21 | 348.22 | 23 | 0.919 | 0.98631319 | 7621959.591 | 7002448.603 |
| P11142 | 44 | 1983.81 | 53 | 0.975 | 0.98631319 | 273616406.6 | 266665418.3 |
| P11172 | 2 | 3.85 | 2 | 0.865 | 0.98631319 | 190912.6615 | 165230.8932 |
| P11441 | 8 | 84.46 | 8 | 1.015 | 0.98631319 | 5315600.564 | 5392889.293 |
| P11940 | 13 | 99.09 | 18 | 0.872 | 0.98631319 | 5997360.517 | 5229295.452 |
| P12236 | 1 | 74.81 | 9 | 0.813 | 0.98631319 | 281136.7826 | 228611.0177 |
| P12956 | 19 | 145.21 | 19 | 0.906 | 0.98631319 | 6042220.48 | 5475222.245 |
| P13010 | 22 | 129.92 | 22 | 1.15 | 0.98631319 | 8258122.483 | 9495677.922 |
| P13639 | 5 | 32.32 | 6 | 1.068 | 0.98631319 | 949673.3518 | 1014329.538 |
| P13645 | 28 | 1143.54 | 37 | 1.06 | 0.98631319 | 115541226.8 | 122435232.1 |
| P13647 | 14 | 368.76 | 29 | 1.011 | 0.98631319 | 8651932.416 | 8746601.903 |
| P14314 | 3 | 11.52 | 3 | 0.972 | 0.98631319 | 479168.2336 | 465867.0554 |
| P14625 | 8 | 53.44 | 10 | 0.952 | 0.98631319 | 1130360.977 | 1075591.915 |
| P14678 | 6 | 127.07 | 6 | 0.88 | 0.98631319 | 18311875.16 | 16110064.47 |
| P14866 | 7 | 14.02 | 7 | 0.776 | 0.98631319 | 829383.3742 | 643499.8323 |
| P14868 | 19 | 134.17 | 19 | 1.33 | 0.98631319 | 5341705.741 | 7105445.816 |
| P14923 | 6 | 4.16 | 6 | 0.966 | 0.98631319 | 131128.2238 | 126722.0264 |
| P15531 | 3 | 31.12 | 3 | 0.715 | 0.98631319 | 1068946.207 | 764736.4772 |
| P15880 | 11 | 109.86 | 11 | 1.038 | 0.98631319 | 11977248.91 | 12436261.23 |
| P15924 | 11 | 23.56 | 11 | 0.744 | 0.98631319 | 906682.0929 | 674528.662 |
| P17844 | 15 | 202.46 | 23 | 1.073 | 0.98631319 | 19164141.89 | 20572488.69 |
| P17980 | 14 | 125.73 | 14 | 0.965 | 0.98631319 | 6482740.56 | 6253157.227 |
| P17987 | 24 | 270.9 | 24 | 1.234 | 0.98631319 | 9214711.813 | 11375552.47 |
| P18077 | 3 | 5.43 | 3 | 0.829 | 0.98631319 | 680966.1872 | 564806.5049 |
| P18124 | 9 | 35.84 | 9 | 0.888 | 0.98631319 | 3008746.397 | 2670810.384 |
| P19338 | 26 | 303.25 | 26 | 0.874 | 0.98631319 | 17501659.05 | 15302057.46 |
| P19474 | 10 | 70.9 | 10 | 0.803 | 0.98631319 | 4045108.088 | 3250237.563 |
| P20042 | 8 | 28.71 | 8 | 0.922 | 0.98631319 | 2500248.092 | 2304253.294 |
| P20700 | 3 | 2.6 | 4 | 0.857 | 0.98631319 | 429386.1168 | 368116.2109 |
| P21333 | 67 | 655.55 | 72 | 0.907 | 0.98631319 | 27416866.84 | 24855547.88 |
| P22626 | 13 | 129.2 | 14 | 0.932 | 0.98631319 | 5316209.282 | 4955730.169 |
| P23246 | 9 | 103.29 | 11 | 0.687 | 0.98631319 | 6504815.611 | 4470585.328 |
| P23284 | 11 | 99.26 | 11 | 1.001 | 0.98631319 | 11577915.74 | 11585649.75 |
| P23396 | 29 | 860.79 | 29 | 0.996 | 0.98631319 | 102237378.8 | 101791321.4 |
| P23458 | 4 | 4.08 | 4 | 0.933 | 0.98631319 | 160202.8553 | 149481.6996 |
| P23588 | 57 | 2255.79 | 57 | 0.953 | 0.98631319 | 167471677 | 159565368.3 |
| P24468 | 2 | 1.83 | 2 | 0.933 | 0.98631319 | 198831.9054 | 185561.463 |
| P25398 | 12 | 192.7 | 12 | 1.058 | 0.98631319 | 6059570.442 | 6408282.207 |
| P25686 | 4 | 33.47 | 4 | 0.971 | 0.98631319 | 1276332.574 | 1239124.907 |
| P25705 | 19 | 236.49 | 19 | 1.029 | 0.98631319 | 13435823.21 | 13823549.68 |
| P25786 | 3 | 2.01 | 3 | 0.807 | 0.98631319 | 220106.8376 | 177711.1597 |
| P25787 | 4 | 8.68 | 4 | 0.933 | 0.98631319 | 1836788.767 | 1713781.566 |
| P25789 | 2 | 3.61 | 2 | 0.799 | 0.98631319 | 293921.1983 | 234884.6292 |
| P26599 | 19 | 357.18 | 20 | 1.228 | 0.98631319 | 17620086.45 | 21633079.32 |
| P26640 | 5 | 26.04 | 5 | 1.022 | 0.98631319 | 334996.6004 | 342336.7761 |
| P26641 | 20 | 282.21 | 20 | 1.018 | 0.98631319 | 22973171.02 | 23382386.45 |
| P27348 | 3 | 10.93 | 3 | 1.09 | 0.98631319 | 654593.0824 | 713311.7453 |
| P27635 | 4 | 6.96 | 4 | 0.962 | 0.98631319 | 1187156.078 | 1141781.823 |
| P27708 | 11 | 33.96 | 11 | 0.918 | 0.98631319 | 733635.6558 | 673665.4263 |
| P27816 | 8 | 67.7 | 8 | 1.089 | 0.98631319 | 968499.4283 | 1054440.903 |
| P27824 | 38 | 874.62 | 38 | 1.022 | 0.98631319 | 92310867.52 | 94337423.1 |
| P28066 | 2 | 22.06 | 2 | 1.21 | 0.98631319 | 178000.936 | 215325.4371 |
| P28072 | 3 | 29.85 | 3 | 1.035 | 0.98631319 | 692657.0109 | 717099.7813 |
| P28074 | 5 | 28.16 | 5 | 1.007 | 0.98631319 | 697396.3305 | 702603.9283 |

| | | | | | | | |
|---|---|---|---|---|---|---|---|
| P29144 | 2 | 0 | 2 | 0.846 | 0.98631319 | 188341.8147 | 159358.9059 |
| P29692 | 2 | 17.78 | 2 | 0.828 | 0.98631319 | 431443.4586 | 357404.7439 |
| P30050 | 6 | 125.8 | 6 | 0.973 | 0.98631319 | 6217796.575 | 6049150.931 |
| P30101 | 2 | 4.31 | 2 | 0.923 | 0.98631319 | 164832.4281 | 152168.6396 |
| P30153 | 5 | 34.05 | 8 | 0.676 | 0.98631319 | 510798.6923 | 345549.5329 |
| P31689 | 17 | 249.91 | 17 | 0.933 | 0.98631319 | 19609872.28 | 18298300.13 |
| P31943 | 7 | 377.36 | 19 | 1.142 | 0.98631319 | 35049252.66 | 40041594.24 |
| P32322 | 4 | 25.3 | 5 | 1.349 | 0.98631319 | 167564.0705 | 225978.6899 |
| P32969 | 3 | 15.23 | 3 | 0.856 | 0.98631319 | 718870.6737 | 615534.1016 |
| P35268 | 5 | 55.6 | 5 | 1.207 | 0.98631319 | 5927440.583 | 7154620.112 |
| P35527 | 33 | 924.82 | 34 | 0.897 | 0.98631319 | 68861204.01 | 61800081.15 |
| P35606 | 5 | 12.18 | 5 | 1.824 | 0.98631319 | 350443.2336 | 639182.1319 |
| P35637 | 20 | 1628.33 | 22 | 0.944 | 0.98631319 | 124038364.6 | 117032691.7 |
| P35908 | 31 | 790.49 | 41 | 1.055 | 0.98631319 | 52731713.76 | 55638899.46 |
| P35998 | 22 | 265.18 | 22 | 1.171 | 0.98631319 | 7556580.655 | 8848903.755 |
| P36542 | 3 | 9.07 | 3 | 0.394 | 0.98631319 | 884717.9064 | 348716.5195 |
| P36551 | 2 | 0 | 2 | 1.001 | 0.98631319 | 3455528.188 | 3459942.857 |
| P37108 | 2 | 17.67 | 2 | 1.045 | 0.98631319 | 388484.2943 | 405873.3267 |
| P38159 | 7 | 5.36 | 7 | 0.771 | 0.98631319 | 911465.3775 | 702824.6875 |
| P38646 | 7 | 71.99 | 8 | 0.94 | 0.98631319 | 544093.4412 | 511710.2046 |
| P38919 | 2 | 7.62 | 4 | 1.096 | 0.98631319 | 126699.1208 | 138895.0725 |
| P39019 | 14 | 119.35 | 14 | 1.016 | 0.98631319 | 11296414.04 | 11471601.53 |
| P39023 | 7 | 25.54 | 7 | 1.264 | 0.98631319 | 1050580.037 | 1328110.884 |
| P39656 | 6 | 39.84 | 6 | 0.914 | 0.98631319 | 2556261.641 | 2337118.779 |
| P40227 | 11 | 83.26 | 11 | 0.902 | 0.98631319 | 3175946.459 | 2864049.57 |
| P40616 | 3 | 14.6 | 3 | 1.038 | 0.98631319 | 552833.8167 | 573612.933 |
| P40939 | 16 | 70.02 | 16 | 1.164 | 0.98631319 | 2178312.547 | 2535291.934 |
| P41091 | 14 | 133.91 | 14 | 1.116 | 0.98631319 | 7850997.618 | 8759426.267 |
| P41252 | 22 | 169.09 | 22 | 0.887 | 0.98631319 | 11857532.46 | 10522897.16 |
| P42345 | 6 | 1.74 | 6 | 0.877 | 0.98631319 | 2455471.448 | 2154360.707 |
| P42677 | 5 | 60.69 | 5 | 0.889 | 0.98631319 | 10210699.09 | 9075958.236 |
| P42704 | 3 | 10.44 | 3 | 0.997 | 0.98631319 | 131798.4788 | 131403.5987 |
| P43243 | 6 | 4.08 | 6 | 0.731 | 0.98631319 | 633631.6066 | 463369.8823 |
| P43246 | 5 | 9.9 | 5 | 0.928 | 0.98631319 | 358131.6492 | 332186.3219 |
| P43686 | 19 | 143.01 | 20 | 0.966 | 0.98631319 | 12385989.59 | 11958948.53 |
| P46777 | 9 | 55.82 | 9 | 0.766 | 0.98631319 | 2356261.429 | 1804429.227 |
| P46779 | 3 | 13.77 | 3 | 0.813 | 0.98631319 | 260204.41 | 211421.9408 |
| P46781 | 10 | 71.71 | 10 | 1.017 | 0.98631319 | 7146586.654 | 7269165.294 |
| P46782 | 16 | 297.04 | 16 | 0.936 | 0.98631319 | 36543325.64 | 34218472.69 |
| P46783 | 9 | 109.68 | 9 | 0.933 | 0.98631319 | 9952237.94 | 9288535.664 |
| P46821 | 3 | 8.78 | 3 | 3.295 | 0.98631319 | 122242.399 | 402811.3443 |
| P46940 | 2 | 0 | 2 | 1.003 | 0.98631319 | 436498.9693 | 437598.383 |
| P46977 | 3 | 23.7 | 3 | 0.972 | 0.98631319 | 642743.7605 | 624463.8414 |
| P47897 | 11 | 51.89 | 11 | 1 | 0.98631319 | 2593565.264 | 2592776.775 |
| P48444 | 2 | 11.25 | 2 | 0.86 | 0.98631319 | 268237.6489 | 230721.3236 |
| P48556 | 9 | 64.23 | 9 | 0.96 | 0.98631319 | 5137335.568 | 4929792.538 |
| P48634 | 3 | 5.82 | 3 | 0.878 | 0.98631319 | 55558.27683 | 48801.44866 |
| P48643 | 18 | 164.37 | 19 | 1.082 | 0.98631319 | 5941602.651 | 6427112.846 |
| P48681 | 2 | 3.55 | 2 | 1.021 | 0.98631319 | 247366.7552 | 252513.5632 |
| P49207 | 2 | 0 | 2 | 0.985 | 0.98631319 | 100294.2828 | 98834.2668 |
| P49327 | 3 | 5.7 | 3 | 1.075 | 0.98631319 | 322430.921 | 346492.9033 |
| P49368 | 20 | 192.36 | 20 | 1.034 | 0.98631319 | 11386750.33 | 11774011.49 |
| P49755 | 5 | 15.47 | 5 | 1.099 | 0.98631319 | 714263.7417 | 785211.0982 |
| P49902 | 16 | 125.9 | 16 | 0.86 | 0.98631319 | 6717936.196 | 5774672.29 |
| P50402 | 2 | 21.17 | 2 | 0.946 | 0.98631319 | 1029008.167 | 973835.5641 |
| P50748 | 5 | 14.01 | 5 | 1.01 | 0.98631319 | 437623.4184 | 442214.6911 |
| P50914 | 3 | 25.2 | 3 | 1.462 | 0.98631319 | 1444574.738 | 2111500.047 |
| P50990 | 24 | 217.13 | 24 | 1.017 | 0.98631319 | 8213009.259 | 8353308.987 |
| P51572 | 5 | 29.32 | 5 | 0.938 | 0.98631319 | 1985875.791 | 1862732.284 |
| P51648 | 3 | 15.61 | 3 | 1.164 | 0.98631319 | 431275.2716 | 501837.01 |
| P51665 | 10 | 108.48 | 10 | 0.999 | 0.98631319 | 4884480.255 | 4881400.483 |
| P51991 | 5 | 38.52 | 5 | 1.038 | 0.98631319 | 1466906.097 | 1522457.821 |
| P52272 | 21 | 100.57 | 21 | 1.069 | 0.98631319 | 39362898.75 | 42086598.54 |
| P52292 | 12 | 155.46 | 12 | 1.051 | 0.98631319 | 5223692.411 | 5491722.618 |
| P52306 | 16 | 171.35 | 17 | 1.118 | 0.98631319 | 7005625.262 | 7829731.216 |
| P52732 | 81 | 1592.86 | 81 | 1.055 | 0.98631319 | 145138952.8 | 153076526.7 |
| P53618 | 7 | 42.75 | 7 | 0.805 | 0.98631319 | 1869341.477 | 1504657.619 |
| P53621 | 6 | 9.82 | 6 | 0.896 | 0.98631319 | 338776.4041 | 303492.6953 |
| P54105 | 2 | 57 | 2 | 1.141 | 0.98631319 | 1330704.64 | 1518034.809 |
| P54136 | 16 | 137 | 16 | 0.991 | 0.98631319 | 4919641.47 | 4876679.398 |
| P54709 | 2 | 6.79 | 2 | 1.463 | 0.98631319 | 220713.0949 | 323007.6967 |
| P55036 | 9 | 105.09 | 9 | 1.052 | 0.98631319 | 4926777.236 | 5185368.99 |
| P55060 | 54 | 993.91 | 54 | 1.019 | 0.98631319 | 80997931.9 | 82559231.14 |
| P55072 | 74 | 1938.57 | 74 | 1.031 | 0.98631319 | 166523349.3 | 171640896.3 |
| P55081 | 3 | 17.29 | 3 | 0.837 | 0.98631319 | 947889.2999 | 793111.0693 |
| P55084 | 5 | 49.28 | 5 | 0.904 | 0.98631319 | 2077164.672 | 1877576.577 |
| P55795 | 1 | 237.28 | 12 | 0.882 | 0.98631319 | 321275.1285 | 283385.8108 |
| P55884 | 15 | 103.02 | 15 | 0.991 | 0.98631319 | 6474452.872 | 6414434.005 |
| P56192 | 12 | 77.2 | 12 | 1.097 | 0.98631319 | 2445519.893 | 2682285.252 |
| P57088 | 7 | 51.42 | 7 | 0.825 | 0.98631319 | 4936603.392 | 4073478.078 |
| P57678 | 27 | 185.47 | 27 | 0.967 | 0.98631319 | 9959351.012 | 9629622.514 |
| P60228 | 17 | 188.24 | 17 | 1.008 | 0.98631319 | 9986678.699 | 10067547.87 |
| P60842 | 4 | 23.81 | 6 | 0.769 | 0.98631319 | 2192016.745 | 1685403.543 |
| P60866 | 9 | 205.6 | 9 | 1.08 | 0.98631319 | 19609490.53 | 21181892.13 |
| P60891 | 3 | 8.26 | 3 | 0.918 | 0.98631319 | 93694.35609 | 86006.35748 |
| P60900 | 5 | 20.57 | 5 | 1.121 | 0.98631319 | 838620.7361 | 939814.6794 |
| P61163 | 7 | 53.1 | 7 | 0.889 | 0.98631319 | 2424719.965 | 2155683.079 |
| P61247 | 22 | 304.61 | 22 | 0.925 | 0.98631319 | 37183503.4 | 34391369.5 |
| P61353 | 6 | 50.93 | 6 | 1.017 | 0.98631319 | 2234710.183 | 2272296.662 |
| P61619 | 4 | 18.84 | 4 | 0.926 | 0.98631319 | 616976.4905 | 571065.5647 |
| P61803 | 3 | 23.76 | 3 | 0.809 | 0.98631319 | 1212096.181 | 980693.2608 |
| P62081 | 15 | 257.2 | 15 | 1.015 | 0.98631319 | 24454094.3 | 24827846.12 |
| P62191 | 11 | 87.22 | 13 | 1.002 | 0.98631319 | 4191635.467 | 4198504.493 |
| P62195 | 17 | 188.67 | 19 | 1.017 | 0.98631319 | 11113083.46 | 11299143.36 |
| P62241 | 11 | 105.06 | 11 | 1.008 | 0.98631319 | 8057245.217 | 8122858.68 |
| P62244 | 12 | 330.96 | 12 | 0.988 | 0.98631319 | 30457196.12 | 30096065.93 |
| P62249 | 15 | 215.64 | 15 | 0.969 | 0.98631319 | 24096324.21 | 23346158.94 |
| P62263 | 10 | 208.8 | 10 | 1.009 | 0.98631319 | 13894353.87 | 14015412.94 |
| P62266 | 4 | 39.67 | 4 | 1.09 | 0.98631319 | 6493691.536 | 7081328.267 |
| P62269 | 14 | 197 | 14 | 1.012 | 0.98631319 | 15460064.8 | 15639080.51 |
| P62277 | 12 | 175.69 | 12 | 0.906 | 0.98631319 | 22173181.34 | 20084723.08 |
| P62280 | 15 | 281.3 | 15 | 1.044 | 0.98631319 | 19927214.27 | 20794155.12 |
| P62308 | 2 | 2.24 | 2 | 1.25 | 0.98631319 | 114586.3987 | 143233.0844 |
| P62316 | 3 | 23.92 | 3 | 0.746 | 0.98631319 | 1216353.212 | 906812.8717 |
| P62318 | 4 | 127.66 | 4 | 0.894 | 0.98631319 | 5937848.733 | 5305933.471 |
| P62333 | 19 | 150.73 | 19 | 0.909 | 0.98631319 | 7279003.495 | 6619513.543 |
| P62424 | 3 | 16.64 | 3 | 0.998 | 0.98631319 | 1214709.769 | 1211914.093 |
| P62701 | 23 | 372.5 | 23 | 1.033 | 0.98631319 | 44092006.61 | 45548833.58 |
| P62826 | 4 | 19.59 | 4 | 0.918 | 0.98631319 | 1165896.991 | 1069976.459 |
| P62829 | 9 | 197.6 | 9 | 1.039 | 0.98631319 | 26176382.97 | 27196396.72 |
| P62841 | 5 | 137.84 | 5 | 0.953 | 0.98631319 | 6366955.539 | 6070839.485 |
| P62847 | 3 | 24.69 | 3 | 0.742 | 0.98631319 | 196404.942 | 145774.1546 |
| P62851 | 2 | 7.31 | 2 | 0.74 | 0.98631319 | 957079.4541 | 708328.8793 |
| P62857 | 2 | 38.01 | 2 | 0.723 | 0.98631319 | 2007404.493 | 1450802.365 |
| P62888 | 2 | 16.42 | 2 | 1.213 | 0.98631319 | 825508.5924 | 1001006.507 |
| P62906 | 12 | 112.92 | 12 | 0.976 | 0.98631319 | 5798709.797 | 5657357.452 |
| P62913 | 6 | 85.96 | 6 | 1.094 | 0.98631319 | 8291517.532 | 9074435.565 |
| P62917 | 3 | 30.96 | 3 | 1.25 | 0.98631319 | 1506513.305 | 1883405.602 |
| P62937 | 3 | 22.41 | 4 | 1.054 | 0.98631319 | 1022988.522 | 1077990.604 |
| P62987 | 1 | 3499.38 | 17 | 0.991 | 0.98631319 | 1051158175 | 1041688639 |
| P63173 | 3 | 125.08 | 3 | 1.355 | 0.98631319 | 3278819.283 | 4443775.5 |

| | | | | | | | |
|---|---|---|---|---|---|---|---|
| P67809 | 5 | 40.1 | 5 | 0.876 | 0.98631319 | 274756.5215 | 240679.2782 |
| P68104 | 29 | 1197.02 | 29 | 1.003 | 0.98631319 | 118298067 | 118631357.5 |
| P68133 | 2 | 198.43 | 15 | 0.707 | 0.98631319 | 1295166.992 | 916038.2773 |
| P68363 | 2 | 2576.82 | 39 | 0.942 | 0.98631319 | 377253618.6 | 355269425.2 |
| P68371 | 2 | 2408.83 | 35 | 1.028 | 0.98631319 | 154333394.9 | 158615102.1 |
| P69905 | 2 | 7.69 | 2 | 0.983 | 0.98631319 | 916228.8375 | 900577.16 |
| P78316 | 2 | 11.02 | 2 | 0.743 | 0.98631319 | 190149.8553 | 141265.4688 |
| P78332 | 13 | 97.77 | 15 | 0.743 | 0.98631319 | 1533259.148 | 1139149.239 |
| P78347 | 19 | 93.76 | 19 | 0.687 | 0.98631319 | 4294937.113 | 2952714.492 |
| P78371 | 25 | 248.97 | 25 | 0.89 | 0.98631319 | 9914256.307 | 8819270.23 |
| P78385 | 2 | 8.84 | 3 | 0.907 | 0.98631319 | 1130578.213 | 1025320.254 |
| P78527 | 101 | 653.04 | 101 | 1.007 | 0.98631319 | 29061382.49 | 29269620.04 |
| P81605 | 2 | 9.81 | 2 | 1.225 | 0.98631319 | 232653.4906 | 284912.3494 |
| P84090 | 5 | 26.39 | 5 | 0.96 | 0.98631319 | 1769446.033 | 1697900.513 |
| P84243 | 3 | 13.4 | 3 | 1.041 | 0.98631319 | 2940170.977 | 3061513.477 |
| P98175 | 60 | 1919.72 | 67 | 0.972 | 0.98631319 | 264694040 | 257388400 |
| P98179 | 6 | 60.97 | 6 | 0.881 | 0.98631319 | 4672208.116 | 4117266.293 |
| Q00325 | 8 | 41.87 | 8 | 1.117 | 0.98631319 | 2913321.782 | 3252846.694 |
| Q00341 | 7 | 24.81 | 7 | 1.013 | 0.98631319 | 1247644.599 | 1263359.506 |
| Q00610 | 4 | 3.81 | 4 | 0.811 | 0.98631319 | 174384.7134 | 141408.9492 |
| Q00839 | 26 | 260.72 | 27 | 0.997 | 0.98631319 | 20598594.48 | 20545229.77 |
| Q01082 | 52 | 341.58 | 53 | 1.057 | 0.98631319 | 11348398.8 | 11999794.89 |
| Q01804 | 48 | 516.54 | 49 | 1.011 | 0.98631319 | 21355284.87 | 21592182.26 |
| Q01844 | 12 | 759.01 | 12 | 0.946 | 0.98631319 | 39341037.02 | 37205069.88 |
| Q02413 | 2 | 2.24 | 2 | 1.075 | 0.98631319 | 103878.8803 | 111662.8385 |
| Q02543 | 3 | 17.51 | 3 | 0.893 | 0.98631319 | 1193088.711 | 1065115.821 |
| Q04637 | 29 | 196.27 | 29 | 1.024 | 0.98631319 | 12720086.66 | 13029492.89 |
| Q04695 | 6 | 232.49 | 17 | 0.998 | 0.98631319 | 1154108.725 | 1151971.872 |
| Q05086 | 4 | 32.74 | 4 | 0.785 | 0.98631319 | 890479.5109 | 699248.4913 |
| Q06830 | 3 | 12.87 | 3 | 0.941 | 0.98631319 | 356034.8409 | 335077.9844 |
| Q07666 | 5 | 14.06 | 5 | 1.006 | 0.98631319 | 920704.1422 | 926416.1113 |
| Q08211 | 26 | 164.94 | 26 | 0.936 | 0.98631319 | 9387778.596 | 8784762.866 |
| Q08945 | 7 | 27.33 | 7 | 1.53 | 0.98631319 | 736535.4843 | 1127216.385 |
| Q12874 | 14 | 141.34 | 14 | 0.819 | 0.98631319 | 9639585.77 | 7890931.176 |
| Q12904 | 9 | 60.71 | 9 | 0.787 | 0.98631319 | 863906.142 | 680235.1791 |
| Q12905 | 5 | 32.5 | 5 | 0.912 | 0.98631319 | 1826706.694 | 1665538.235 |
| Q13045 | 2 | 9.8 | 2 | 0.947 | 0.98631319 | 68439.66621 | 64832.2403 |
| Q13123 | 5 | 14.94 | 5 | 1.123 | 0.98631319 | 957110.1423 | 1074710.374 |
| Q13148 | 4 | 10.17 | 4 | 0.931 | 0.98631319 | 468437.472 | 436015.9159 |
| Q13151 | 6 | 35.95 | 6 | 0.899 | 0.98631319 | 847680.6396 | 762104.1216 |
| Q13200 | 37 | 389.8 | 37 | 0.959 | 0.98631319 | 18755887.5 | 17981990.26 |
| Q13257 | 3 | 7.48 | 3 | 0.999 | 0.98631319 | 961736.504 | 961018.6931 |
| Q13263 | 5 | 50.99 | 5 | 0.858 | 0.98631319 | 272157.4827 | 233642.3532 |
| Q13283 | 8 | 72.56 | 9 | 1.122 | 0.98631319 | 3497291.095 | 3922274.917 |
| Q13310 | 4 | 45.34 | 9 | 0.936 | 0.98631319 | 222536.6264 | 208277.9944 |
| Q13347 | 10 | 84.46 | 10 | 0.876 | 0.98631319 | 9186830.749 | 8043804.842 |
| Q13464 | 25 | 104.66 | 25 | 1.021 | 0.98631319 | 25850134.31 | 26404792.55 |
| Q13535 | 2 | 0 | 2 | 0.838 | 0.98631319 | 110824.9596 | 92864.02344 |
| Q13561 | 7 | 23.05 | 7 | 0.852 | 0.98631319 | 745928.8878 | 635366.6355 |
| Q13620 | 2 | 3.64 | 2 | 1.278 | 0.98631319 | 64347.35364 | 82216.95758 |
| Q13813 | 53 | 376.07 | 53 | 0.93 | 0.98631319 | 13019846.57 | 12110761.27 |
| Q13838 | 3 | 5.7 | 3 | 1.277 | 0.98631319 | 274045.4744 | 349860.8875 |
| Q13895 | 11 | 103.5 | 11 | 1.218 | 0.98631319 | 2622374.722 | 3193165.741 |
| Q14004 | 2 | 13.18 | 2 | 0.971 | 0.98631319 | 13216874.68 | 12833287.14 |
| Q14008 | 3 | 26.69 | 3 | 0.887 | 0.98631319 | 468921.3068 | 416135.3975 |
| Q14103 | 8 | 68.34 | 10 | 1.042 | 0.98631319 | 3226352.267 | 3362153.924 |
| Q14152 | 26 | 137.01 | 26 | 1.082 | 0.98631319 | 6359826.34 | 6884091.058 |
| Q14157 | 17 | 179.85 | 17 | 0.947 | 0.98631319 | 5857570.221 | 5547072.183 |
| Q14203 | 13 | 35.41 | 13 | 1.477 | 0.98631319 | 1279341.336 | 1889508.323 |
| Q14254 | 5 | 14.85 | 5 | 0.33 | 0.98631319 | 255445046.1 | 84328640.33 |
| Q14331 | 5 | 37.58 | 5 | 1.4 | 0.98631319 | 1236846.168 | 1731962.151 |
| Q14498 | 8 | 50.61 | 8 | 0.811 | 0.98631319 | 2094449.943 | 1698516.676 |
| Q14681 | 9 | 141.1 | 10 | 1.041 | 0.98631319 | 5886851.239 | 6129813.46 |
| Q14697 | 17 | 103.98 | 17 | 1.001 | 0.98631319 | 3552661.993 | 3555641.94 |
| Q14974 | 25 | 297.92 | 25 | 0.972 | 0.98631319 | 21621523.06 | 21020814.71 |
| Q15008 | 21 | 255.22 | 21 | 1.151 | 0.98631319 | 10859464.71 | 12497719.7 |
| Q15018 | 9 | 44.44 | 9 | 0.916 | 0.98631319 | 1500959.992 | 1375394.957 |
| Q15021 | 6 | 13.93 | 6 | 1.163 | 0.98631319 | 540141.6588 | 628190.7052 |
| Q15029 | 20 | 146.29 | 21 | 1.015 | 0.98631319 | 9565909.58 | 9711027.086 |
| Q15057 | 34 | 364.11 | 34 | 0.993 | 0.98631319 | 18007044.47 | 17878919.56 |
| Q15233 | 6 | 72.27 | 8 | 0.984 | 0.98631319 | 2801838.348 | 2757057.304 |
| Q15365 | 4 | 84.85 | 8 | 0.853 | 0.98631319 | 1056363.219 | 900940.8361 |
| Q15366 | 4 | 78.5 | 8 | 0.912 | 0.98631319 | 2898892.202 | 2645175.189 |
| Q15386 | 6 | 23.98 | 6 | 1.067 | 0.98631319 | 910000.3403 | 970805.7293 |
| Q15393 | 48 | 784.71 | 48 | 0.966 | 0.98631319 | 54852701.84 | 52992565.54 |
| Q15427 | 4 | 53.73 | 4 | 1.002 | 0.98631319 | 2535849.213 | 2541934.755 |
| Q15428 | 8 | 98.88 | 8 | 0.738 | 0.98631319 | 4074777.039 | 3007474.997 |
| Q15459 | 28 | 289.39 | 28 | 1.019 | 0.98631319 | 14807445.71 | 15094465.69 |
| Q15717 | 2 | 15.84 | 2 | 0.844 | 0.98631319 | 318116.6032 | 268593.875 |
| Q15750 | 15 | 128.61 | 15 | 0.951 | 0.98631319 | 3998887.12 | 3801047.83 |
| Q15773 | 2 | 3.28 | 2 | 1.386 | 0.98631319 | 119178.9155 | 165165.7439 |
| Q16186 | 5 | 32.74 | 5 | 0.736 | 0.98631319 | 2092250.144 | 1538881.726 |
| Q16531 | 16 | 70.32 | 16 | 1.195 | 0.98631319 | 3039276.977 | 3631909.576 |
| Q16637 | 5 | 19.34 | 5 | 1.077 | 0.98631319 | 1258493.799 | 1354831.808 |
| Q16777 | 3 | 45.95 | 3 | 0.95 | 0.98631319 | 4424935.746 | 4202501.014 |
| Q16778 | 1 | 67.5 | 5 | 0.829 | 0.98631319 | 343771.7289 | 284874.6929 |
| Q16875 | 2 | 1.66 | 2 | 1.112 | 0.98631319 | 243303.5652 | 270507.9665 |
| Q29RF7 | 3 | 7.21 | 3 | 0.899 | 0.98631319 | 82282.18196 | 73982.80802 |
| O00743 | 3 | 24.68 | 3 | 1.548 | 0.98631319 | 227578.3005 | 352388.3955 |
| Q2NL82 | 7 | 31.95 | 7 | 0.936 | 0.98631319 | 1184172.706 | 1107945.91 |
| Q2TAY7 | 11 | 130.88 | 11 | 0.936 | 0.98631319 | 8292460.116 | 7763857.62 |
| Q32MZ4 | 2 | 15.68 | 2 | 1.329 | 0.98631319 | 195888.4591 | 260389.6678 |
| Q53GQ0 | 13 | 154.52 | 13 | 0.816 | 0.98631319 | 8540011.458 | 6966826.073 |
| Q53GS9 | 7 | 27.42 | 7 | 0.806 | 0.98631319 | 914179.8003 | 737196.7679 |
| Q5C9Z4 | 2 | 8.08 | 2 | 1.09 | 0.98631319 | 112657.239 | 122819.73 |
| Q5D862 | 2 | 17.45 | 2 | 0.694 | 0.98631319 | 2439445.295 | 1692576.766 |
| Q5QNW6 | 2 | 86.59 | 6 | 1.035 | 0.98631319 | 6856678.978 | 7095608.785 |
| Q5SW79 | 3 | 12.58 | 3 | 0.923 | 0.98631319 | 160566.5654 | 148214.6207 |
| Q5T4S7 | 11 | 26.2 | 11 | 0.99 | 0.98631319 | 645728.4453 | 638990.9802 |
| Q5T6F2 | 2 | 0 | 2 | 0.873 | 0.98631319 | 158440.6821 | 138304.7465 |
| Q5T749 | 2 | 5.51 | 2 | 1.012 | 0.98631319 | 361963.9693 | 366383.5677 |
| Q5T8P6 | 6 | 21.17 | 6 | 0.976 | 0.98631319 | 619844.7615 | 604687.6443 |
| Q5TAX3 | 2 | 0 | 2 | 0.913 | 0.98631319 | 146485.6605 | 133757.2245 |
| Q5VYK3 | 36 | 254.88 | 36 | 1.011 | 0.98631319 | 8304564.298 | 8392064.161 |
| Q5W0B1 | 34 | 379.34 | 34 | 0.939 | 0.98631319 | 25930413.12 | 24338121.67 |
| Q6AI08 | 4 | 8.89 | 4 | 0.572 | 0.98631319 | 172597.6828 | 98645.57857 |
| Q6N021 | 2 | 4.92 | 2 | 1.233 | 0.98631319 | 72064.64038 | 88848.52399 |
| Q6NUN0 | 2 | 2.82 | 2 | 0.862 | 0.98631319 | 1186614.003 | 1023397.489 |
| Q3V6T2 | 6 | 21.35 | 6 | 1.049 | 0.98631319 | 201956.1269 | 211760.8164 |
| Q6NXE6 | 4 | 23.48 | 4 | 1.018 | 0.98631319 | 2022830.807 | 2060131.665 |
| Q6P158 | 3 | 17.92 | 4 | 0.84 | 0.98631319 | 489291.0644 | 411234.8839 |
| Q6P2Q9 | 56 | 287.5 | 56 | 0.995 | 0.98631319 | 16272656.42 | 16197545.58 |
| Q6P5R6 | 2 | 15.42 | 2 | 0.954 | 0.98631319 | 582007.8014 | 555371.6497 |
| Q6PJG6 | 15 | 62.84 | 15 | 0.796 | 0.98631319 | 2868106.71 | 2282844.952 |
| Q6S8J3 | 1 | 168.86 | 8 | 0.992 | 0.98631319 | 593833.5655 | 588861.8622 |
| Q6UXN9 | 3 | 8.45 | 3 | 0.934 | 0.98631319 | 514265.3819 | 480502.0548 |
| Q6VN20 | 2 | 24.28 | 3 | 0.958 | 0.98631319 | 188068.9374 | 180127.5356 |
| Q6Y7W6 | 2 | 11.7 | 4 | 2.037 | 0.98631319 | 174958.371 | 356452.3219 |
| Q7L1Q6 | 17 | 114.76 | 19 | 0.891 | 0.98631319 | 9821393.306 | 8748118.544 |
| Q7L2H7 | 11 | 50.45 | 11 | 0.811 | 0.98631319 | 1487930.89 | 1206024.914 |
| Q7L5D6 | 14 | 308.53 | 14 | 0.79 | 0.98631319 | 9641728.599 | 7621484.682 |

| | | | | | | | |
|---|---|---|---|---|---|---|---|
| Q7L8L6 | 6 | 23.03 | 6 | 0.681 | 0.98631319 | 875205.1146 | 596223.0084 |
| Q7RTV0 | 8 | 84.41 | 8 | 0.868 | 0.98631319 | 7874550.883 | 6838979.9 |
| Q7Z353 | 3 | 2.73 | 3 | 1.043 | 0.98631319 | 424980.9293 | 443122.7844 |
| Q7Z3U7 | 23 | 108.19 | 23 | 0.981 | 0.98631319 | 5059725.007 | 4962390.81 |
| Q7Z478 | 28 | 240.28 | 30 | 1.076 | 0.98631319 | 8358103.081 | 8997211.723 |
| Q7Z6Z7 | 64 | 382.87 | 64 | 0.922 | 0.98631319 | 20086151.9 | 18511730.36 |
| Q86V81 | 10 | 121.94 | 10 | 1.047 | 0.98631319 | 6783778.12 | 7100896.309 |
| Q86VP6 | 14 | 82.03 | 15 | 0.933 | 0.98631319 | 2247905.74 | 2096255.816 |
| Q86XI2 | 5 | 11.25 | 5 | 1.263 | 0.98631319 | 1223804.911 | 1545797.807 |
| Q86Y56 | 23 | 173.53 | 23 | 0.967 | 0.98631319 | 5695936.306 | 5509647.251 |
| Q86YZ3 | 11 | 45.43 | 11 | 1.2 | 0.98631319 | 2274743.822 | 2730740.21 |
| Q8IWS0 | 5 | 0 | 5 | 0.817 | 0.98631319 | 348693.1381 | 284987.617 |
| Q8IWX8 | 16 | 182.65 | 16 | 1.072 | 0.98631319 | 13101261.6 | 14040644.09 |
| Q8IX12 | 8 | 57.42 | 8 | 0.861 | 0.98631319 | 2177748.013 | 1873990.723 |
| Q8IXB1 | 8 | 22.14 | 8 | 1.007 | 0.98631319 | 696524.8754 | 701168.8778 |
| Q8IZ07 | 3 | 22.95 | 3 | 1.01 | 0.98631319 | 355325.1197 | 358737.022 |
| Q8N1B4 | 2 | 5.32 | 2 | 0.83 | 0.98631319 | 193706.9471 | 160726.9922 |
| Q8N1F7 | 16 | 111.32 | 16 | 0.801 | 0.98631319 | 3900321.573 | 3123992.428 |
| Q8N3C0 | 10 | 20.2 | 10 | 1.021 | 0.98631319 | 563148.9454 | 575239.8238 |
| Q8N5C8 | 8 | 28.15 | 8 | 0.79 | 0.98631319 | 624890.1892 | 493860.8048 |
| Q8N5Z5 | 8 | 113.96 | 8 | 0.405 | 0.98631319 | 7108021.441 | 2875538.168 |
| Q8NBM4 | 2 | 9.75 | 2 | 0.901 | 0.98631319 | 137458.5579 | 123834.1952 |
| Q8NC51 | 12 | 228.36 | 12 | 0.981 | 0.98631319 | 16256896.05 | 15954369.26 |
| Q8ND56 | 16 | 249.05 | 16 | 1.003 | 0.98631319 | 29767890.09 | 29848763.04 |
| Q8NE71 | 5 | 13.82 | 5 | 1.075 | 0.98631319 | 409669.4232 | 440272.1641 |
| Q8TAA3 | 2 | 7.92 | 2 | 0.541 | 0.98631319 | 265101.4221 | 143298.7813 |
| Q8TAT6 | 23 | 210.14 | 23 | 1.196 | 0.98631319 | 12048189.84 | 14411069.04 |
| Q8TC07 | 9 | 49.74 | 9 | 0.9 | 0.98631319 | 1739121.919 | 1564711.331 |
| Q8TC12 | 2 | 24.91 | 3 | 1.13 | 0.98631319 | 148278.3039 | 167623.6818 |
| P30876 | 4 | 15.3 | 4 | 1.195 | 0.98631319 | 138432.4139 | 165414.1647 |
| Q8TCG1 | 23 | 133.95 | 23 | 0.904 | 0.98631319 | 6914309.804 | 6252166.67 |
| Q8TEQ6 | 7 | 5.34 | 7 | 1.011 | 0.98631319 | 355573.9681 | 359422.5988 |
| Q9P2R3 | 2 | 14.91 | 2 | 0.802 | 0.98631319 | 139872.7124 | 112237.756 |
| Q8TEX9 | 39 | 579.23 | 39 | 0.999 | 0.98631319 | 29608884.13 | 29574328.89 |
| Q8WUA2 | 4 | 15.5 | 4 | 0.646 | 0.98631319 | 2026379.282 | 1308500.357 |
| Q8WWC4 | 3 | 11.42 | 3 | 0.834 | 0.98631319 | 881194.3676 | 734836.527 |
| Q8WWY3 | 21 | 327.68 | 21 | 0.895 | 0.98631319 | 27422711.18 | 24533827.56 |
| Q8WXF1 | 3 | 6.62 | 4 | 0.932 | 0.98631319 | 110620584.5 | 103129762.6 |
| Q8WZ42 | 2 | 5.46 | 3 | 1.175 | 0.98631319 | 2192641.072 | 2576298.466 |
| P26373 | 2 | 13.84 | 2 | 2.066 | 0.98631319 | 23844.9234 | 49269.58843 |
| Q92499 | 8 | 52.93 | 8 | 1.033 | 0.98631319 | 2249772.088 | 2324883.447 |
| Q92538 | 10 | 39.82 | 10 | 0.919 | 0.98631319 | 1362213.535 | 1251719.427 |
| Q92616 | 115 | 1149.26 | 115 | 0.992 | 0.98631319 | 66241617.99 | 65718706.22 |
| Q92620 | 3 | 19.75 | 3 | 0.822 | 0.98631319 | 334442.622 | 274817.4609 |
| Q92621 | 15 | 69.26 | 15 | 0.928 | 0.98631319 | 1881027.787 | 1746210.193 |
| Q92769 | 5 | 19.07 | 5 | 0.826 | 0.98631319 | 1844544.876 | 1524289.886 |
| P51571 | 2 | 12.7 | 2 | 9.311 | 0.98631319 | 83907.49287 | 781281.7898 |
| Q92804 | 13 | 343.68 | 15 | 0.978 | 0.98631319 | 15525036.49 | 15181514.52 |
| Q92841 | 7 | 153.49 | 15 | 0.937 | 0.98631319 | 1307106.578 | 1225387.869 |
| Q92973 | 13 | 200.1 | 19 | 0.876 | 0.98631319 | 35711416.9 | 31293206.69 |
| Q92979 | 4 | 43.88 | 4 | 1.28 | 0.98631319 | 602424.8034 | 770832.1227 |
| Q92990 | 9 | 74.99 | 9 | 1.249 | 0.98631319 | 2755383.893 | 3441291.186 |
| Q93008 | 21 | 74.33 | 21 | 0.929 | 0.98631319 | 3602078.96 | 3345183.424 |
| Q969N2 | 2 | 1.99 | 2 | 0.911 | 0.98631319 | 160995.0382 | 146639.2307 |
| Q7RTS9 | 2 | 11.28 | 2 | 0.991 | 0.98631319 | 87138.64877 | 86382.80433 |
| Q969Z0 | 6 | 31.75 | 6 | 0.936 | 0.98631319 | 799898.2383 | 748415.2154 |
| Q96AG4 | 4 | 43.46 | 4 | 0.825 | 0.98631319 | 1405861.196 | 1159867.164 |
| Q96AV8 | 7 | 33.59 | 7 | 0.982 | 0.98631319 | 398985.4795 | 391643.1018 |
| Q9UBB9 | 4 | 11.13 | 4 | 0.806 | 0.98631319 | 211695.5498 | 170625.8906 |
| Q96C36 | 3 | 23.89 | 4 | 0.864 | 0.98631319 | 503536.3726 | 435282.6999 |
| Q96CS3 | 11 | 77.11 | 11 | 1.296 | 0.98631319 | 1912591.22 | 2478267.816 |
| Q96DG6 | 12 | 168.52 | 12 | 1.072 | 0.98631319 | 12978626.94 | 13910658.72 |
| Q96DI7 | 11 | 69.37 | 11 | 1.014 | 0.98631319 | 5847875.576 | 5930442.583 |
| Q96ER3 | 4 | 32.3 | 4 | 0.95 | 0.98631319 | 285945.8189 | 271764.8572 |
| Q96EY1 | 7 | 23.56 | 7 | 1.709 | 0.98631319 | 528550.6785 | 903123.8438 |
| Q9NXR7 | 3 | 10.37 | 3 | 1.239 | 0.98631319 | 188185.8902 | 233213.8359 |
| Q96GA3 | 3 | 11.7 | 3 | 0.883 | 0.98631319 | 261282.3516 | 230594.9255 |
| Q96MR6 | 2 | 28.18 | 2 | 1.375 | 0.98631319 | 89728164.13 | 123377945.8 |
| Q96NR8 | 1 | 1.69 | 2 | 0.847 | 0.98631319 | 168641.8931 | 142770.596 |
| Q96P70 | 21 | 156.51 | 21 | 0.973 | 0.98631319 | 10675897.53 | 10387220.08 |
| Q96QU8 | 7 | 12.94 | 7 | 0.912 | 0.98631319 | 503881.1619 | 459543.0348 |
| Q96R06 | 2 | 4.27 | 2 | 0.829 | 0.98631319 | 154887.1978 | 128413.3978 |
| Q96S66 | 2 | 0 | 2 | 1.052 | 0.98631319 | 124866.5619 | 131303.1738 |
| Q96T76 | 13 | 60.46 | 13 | 0.792 | 0.98631319 | 2071110.603 | 1641298.605 |
| Q99417 | 2 | 5.51 | 2 | 0.909 | 0.98631319 | 280271.4958 | 254749.5547 |
| Q99436 | 2 | 9.79 | 2 | 1.06 | 0.98631319 | 485011.0225 | 514120.9385 |
| Q96SB4 | 2 | 8.98 | 2 | 0.366 | 0.98631319 | 177955.3861 | 65051.4455 |
| Q99459 | 7 | 35.28 | 7 | 0.942 | 0.98631319 | 2201676.403 | 2073187.83 |
| Q99460 | 34 | 336.46 | 34 | 0.98 | 0.98631319 | 19854044.16 | 19452775.63 |
| Q99613 | 13 | 98.05 | 13 | 1.024 | 0.98631319 | 3819059.856 | 3911474.639 |
| Q99615 | 17 | 67.23 | 17 | 1.051 | 0.98631319 | 3283524.436 | 3452396.304 |
| Q99623 | 4 | 10.88 | 4 | 1.385 | 0.98631319 | 103821.6772 | 143801.452 |
| Q99720 | 3 | 74.8 | 3 | 1.006 | 0.98631319 | 3559093.908 | 3580812.303 |
| Q99729 | 5 | 43.71 | 6 | 0.833 | 0.98631319 | 1037461.363 | 863694.5728 |
| Q99832 | 16 | 155.02 | 16 | 1.021 | 0.98631319 | 8108893.783 | 8283104.124 |
| Q99873 | 13 | 130.53 | 13 | 0.816 | 0.98631319 | 4759637.686 | 3884724.891 |
| Q99942 | 3 | 11.22 | 3 | 0.731 | 0.98631319 | 2371639.3 | 1733522.657 |
| O60524 | 3 | 7.42 | 3 | 0.504 | 0.98631319 | 111102.3659 | 56008.33181 |
| Q9BQA1 | 21 | 4107.62 | 21 | 1.01 | 0.98631319 | 513444778.1 | 518719658.7 |
| Q9BQE3 | 2 | 2188.72 | 38 | 0.852 | 0.98631319 | 3256507.076 | 2775851.143 |
| Q9BRS2 | 17 | 183.79 | 17 | 1.046 | 0.98631319 | 9560776.559 | 10004122.79 |
| Q9BRX9 | 2 | 15.6 | 2 | 1.07 | 0.98631319 | 383960.0895 | 410933.5733 |
| Q9BSJ2 | 4 | 3.73 | 4 | 1.016 | 0.98631319 | 540390.0452 | 549045.257 |
| Q9BT22 | 3 | 6.91 | 3 | 1.041 | 0.98631319 | 286882.5144 | 298661.6476 |
| Q9BTW9 | 3 | 10.24 | 3 | 0.932 | 0.98631319 | 419589.861 | 391253.6732 |
| Q9Y6M1 | 1 | 6.54 | 2 | 0.223 | 0.98631319 | 89597.12493 | 20021.98026 |
| P11177 | 3 | 6.26 | 3 | 1.049 | 0.98631319 | 192757.1857 | 202282.0237 |
| O94776 | 2 | 6.15 | 2 | 0.817 | 0.98631319 | 145019.3919 | 118509.3125 |
| P61513 | 2 | 6.14 | 2 | 1.456 | 0.98631319 | 470416.0104 | 684968.79 |
| Q9BY77 | 3 | 5.88 | 3 | 0.918 | 0.98631319 | 41000.1035 | 37635.0279 |
| Q9BTX1 | 3 | 11.02 | 3 | 0.782 | 0.98631319 | 484992.6804 | 379050.3422 |
| Q9NRZ9 | 2 | 5.72 | 2 | 0.741 | 0.98631319 | 84147.14575 | 62393.73486 |
| Q9BUA3 | 13 | 80.3 | 13 | 0.969 | 0.98631319 | 2931021.839 | 2840473.433 |
| Q9BUJ2 | 2 | 5.25 | 3 | 0.857 | 0.98631319 | 85372.66007 | 73138.08895 |
| Q9BUQ8 | 7 | 31.91 | 7 | 1.489 | 0.98631319 | 1057206.904 | 1574586.913 |
| Q9BV68 | 4 | 51.89 | 4 | 1.045 | 0.98631319 | 1704643.817 | 1781540.794 |
| Q9BVC6 | 3 | 43.49 | 3 | 1.084 | 0.98631319 | 6309343.421 | 6841148.738 |
| Q9BVI4 | 5 | 31.33 | 5 | 1.022 | 0.98631319 | 1817004.929 | 1857168.267 |
| Q9BVK6 | 4 | 29.57 | 4 | 0.892 | 0.98631319 | 461533.8161 | 411548.8132 |
| P49721 | 3 | 5.3 | 3 | 1.384 | 0.98631319 | 440063.0485 | 608867.2611 |
| Q92600 | 3 | 5.25 | 3 | 0.696 | 0.98631319 | 686486.2835 | 477765.5556 |
| Q9BWU0 | 15 | 125.24 | 15 | 0.962 | 0.98631319 | 6115219.489 | 5880988.303 |
| Q9BXW9 | 14 | 98.57 | 14 | 0.57 | 0.98631319 | 4429905.308 | 2523655.405 |
| Q9BY44 | 8 | 49.29 | 8 | 1.034 | 0.98631319 | 964629.9317 | 997571.1661 |
| Q9C0B7 | 2 | 0 | 2 | 0.715 | 0.98631319 | 121638.3321 | 86979.19531 |
| Q9C0E2 | 5 | 25.24 | 6 | 0.944 | 0.98631319 | 531699.1646 | 501801.4307 |
| Q9GZV4 | 2 | 0 | 2 | 1.052 | 0.98631319 | 155350.0112 | 163465.4529 |
| Q9H0A0 | 4 | 12.44 | 4 | 0.848 | 0.98631319 | 262040.1209 | 222242.5292 |
| Q9H0G5 | 3 | 4.51 | 3 | 0.756 | 0.98631319 | 797449.5525 | 603055.5621 |
| Q9H0U3 | 2 | 10.63 | 2 | 1.05 | 0.98631319 | 1352600.511 | 1420373.244 |
| Q9H2M9 | 4 | 16.02 | 4 | 1.65 | 0.98631319 | 241144.0741 | 397812.3761 |

| | | | | | | | |
|---|---|---|---|---|---|---|---|
| Q9H3N1 | 2 | 17.58 | 2 | 2.141 | 0.98631319 | 188558.569 | 403680.3699 |
| Q9H3U1 | 21 | 167.8 | 21 | 1.164 | 0.98631319 | 4826360.23 | 5617152.278 |
| Q9H583 | 31 | 149.42 | 31 | 0.84 | 0.98631319 | 6393192.36 | 5367265.576 |
| Q9H5Z1 | 3 | 9.93 | 3 | 1.078 | 0.98631319 | 268953.2088 | 289890.9367 |
| Q9H7D7 | 6 | 26.42 | 6 | 1.742 | 0.98631319 | 481628.9099 | 839231.6094 |
| Q9HAV4 | 39 | 416.43 | 39 | 0.996 | 0.98631319 | 24873628 | 24776658.48 |
| Q9HCN8 | 2 | 4.32 | 2 | 1.006 | 0.98631319 | 181709.0412 | 182866.8752 |
| Q9NNW5 | 4 | 5.47 | 4 | 1.203 | 0.98631319 | 403298.4092 | 485129.2697 |
| Q9NP73 | 18 | 110.49 | 19 | 0.953 | 0.98631319 | 12880525.22 | 12280982.72 |
| Q9NQC3 | 3 | 25.07 | 3 | 0.911 | 0.98631319 | 712719.2019 | 649165.7541 |
| Q9NR09 | 15 | 36.47 | 15 | 0.877 | 0.98631319 | 1137350.503 | 997229.8135 |
| Q9UPW6 | 2 | 2.04 | 2 | 0.516 | 0.98631319 | 79297.63458 | 40903.43657 |
| Q9NR30 | 3 | 12.96 | 3 | 1.128 | 0.98631319 | 860552.9839 | 970499.8088 |
| Q9BQE5 | 2 | 1.99 | 2 | 3.744 | 0.98631319 | 19333.57025 | 72381.50557 |
| P42356 | 2 | 1.98 | 2 | 1.233 | 0.98631319 | 85029.68232 | 104831.0658 |
| Q9NRG9 | 6 | 40.3 | 6 | 1.002 | 0.98631319 | 1533459.354 | 1537156.454 |
| Q14692 | 2 | 1.88 | 2 | 0.463 | 0.98631319 | 135896.3455 | 62967.93586 |
| Q9NRP0 | 2 | 19.56 | 2 | 0.929 | 0.98631319 | 1949978.347 | 1811744.037 |
| Q9NRP7 | 2 | 0 | 2 | 0.916 | 0.98631319 | 101648763.2 | 93073871.28 |
| Q9NRX1 | 5 | 24.31 | 5 | 0.866 | 0.98631319 | 693106.1884 | 600156.2258 |
| Q9NTJ3 | 9 | 51.13 | 9 | 1.047 | 0.98631319 | 1321368.449 | 1384054.982 |
| Q9NU22 | 26 | 90.7 | 27 | 0.901 | 0.98631319 | 4239579.199 | 3819669.129 |
| Q9NV70 | 3 | 11.09 | 3 | 0.888 | 0.98631319 | 127618.0034 | 113321.5727 |
| Q9NVI1 | 15 | 104.4 | 16 | 1.061 | 0.98631319 | 2657283.48 | 2819327.445 |
| Q9NWV8 | 2 | 14.11 | 2 | 1.233 | 0.98631319 | 99687.80454 | 122896.5534 |
| P50416 | 2 | 0 | 2 | 0.668 | 0.98631319 | 165096.0466 | 110345.7969 |
| Q9NXV2 | 17 | 834.72 | 18 | 1.122 | 0.98631319 | 65948078.98 | 73970806.51 |
| Q9NYF8 | 9 | 25.89 | 9 | 1.432 | 0.98631319 | 1533568.137 | 2196015.426 |
| Q9NYJ8 | 3 | 34.08 | 3 | 0.982 | 0.98631319 | 412510.3788 | 405008.1448 |
| Q13509 | 1 | 1394.69 | 18 | 0.974 | 0.98631319 | 638925.0491 | 622436.3366 |
| P60709 | 1 | 394.82 | 24 | 0.935 | 0.98631319 | 39511689.07 | 36945196.51 |
| A6NHL2 | 1 | 306.77 | 7 | 0.073 | 0.98631319 | 583510.1564 | 42727.84834 |
| O14979 | 2 | 32.08 | 4 | 3.194 | 0.98631319 | 82784.42654 | 264453.8161 |
| Q16576 | 1 | 15.36 | 6 | 1.587 | 0.98631319 | 38611.18299 | 61279.48489 |
| Q09028 | 1 | 14.72 | 6 | 1.02 | 0.98631319 | 1606593.86 | 1639187.427 |
| O14773 | 2 | 9.31 | 2 | 1.045 | 0.98631319 | 204255.936 | 213491.2188 |
| P83369 | 2 | 7.21 | 2 | 0.849 | 0.98631319 | 166737.1938 | 141507.2509 |
| O00139 | 2 | 4.58 | 2 | 1.897 | 0.98631319 | 43768.91413 | 83036.43094 |
| Q9Y678 | 2 | 2 | 2 | 0.514 | 0.98631319 | 56507.89844 | 29021.82326 |
| Q9NYU2 | 42 | 367.81 | 42 | 0.907 | 0.98631319 | 27116791.59 | 24589380.82 |
| Q9NZB2 | 2 | 5.17 | 2 | 1.018 | 0.98631319 | 3091117.879 | 3145895.233 |
| P52756 | 2 | 152.9 | 7 | 0.571 | 0.98631319 | 124261.4506 | 70904.23926 |
| O77727 | 1 | 131.24 | 8 | 0.924 | 0.98631319 | 169051.5696 | 156202.0973 |
| Q9UMX0 | 4 | 26.02 | 8 | 1.383 | 0.98631319 | 1297446.478 | 1794527.952 |
| Q9NZI8 | 8 | 50.96 | 10 | 0.955 | 0.98631319 | 1892942.151 | 1807718.592 |
| Q96PK6 | 2 | 16.49 | 2 | 1.322 | 0.98631319 | 89307.46918 | 118057.5807 |
| Q9P1Y5 | 12 | 78.15 | 12 | 0.816 | 0.98631319 | 1972231.685 | 1609033.256 |
| Q9P219 | 2 | 1.66 | 2 | 1.15 | 0.98631319 | 110410.2305 | 127008.1711 |
| Q9P258 | 5 | 31.43 | 5 | 0.892 | 0.98631319 | 551149.0718 | 491351.5168 |
| Q9P2D3 | 2 | 2.27 | 2 | 0.812 | 0.98631319 | 107915.3844 | 87591.76571 |
| Q9P2J5 | 10 | 21.16 | 10 | 0.826 | 0.98631319 | 2461073.265 | 2034052.254 |
| Q9UBB4 | 23 | 289.05 | 23 | 0.952 | 0.98631319 | 12935927.04 | 12308969.58 |
| Q9UBM7 | 2 | 8.4 | 2 | 0.939 | 0.98631319 | 204293.3012 | 191746.1542 |
| Q9UBQ5 | 3 | 5.27 | 3 | 1.004 | 0.98631319 | 1309776.828 | 1315534.622 |
| Q9UBS4 | 4 | 4.26 | 4 | 0.876 | 0.98631319 | 1094204.297 | 958126.5227 |
| Q9UHD9 | 3 | 41 | 7 | 0.986 | 0.98631319 | 519983.2556 | 512571.8951 |
| P18621 | 4 | 7.62 | 4 | 2.435 | 0.98631319 | 260513.4382 | 634240.6606 |
| Q9UHI6 | 15 | 132.51 | 15 | 0.96 | 0.98631319 | 8699944.677 | 8351469.955 |
| P48729 | 2 | 7.11 | 2 | 0.272 | 0.98631319 | 163742.8045 | 44551.07798 |
| Q9UHX1 | 7 | 52.14 | 7 | 0.925 | 0.98631319 | 2063385.757 | 1908324.6 |
| Q9UIA9 | 10 | 35.18 | 10 | 1.221 | 0.98631319 | 1241969.255 | 1515850.815 |
| Q9UKA9 | 3 | 38.91 | 4 | 1.195 | 0.98631319 | 143231.5732 | 171091.197 |
| Q9UKE5 | 2 | 1.88 | 2 | 0.852 | 0.98631319 | 198078.1944 | 168833.2395 |
| Q9UMS4 | 21 | 323.34 | 21 | 1.023 | 0.98631319 | 16486898.58 | 16866872.97 |
| Q9UN86 | 1 | 46.47 | 2 | 1.026 | 0.98631319 | 557463.5087 | 571923.2685 |
| Q14165 | 2 | 4.75 | 2 | 1.539 | 0.98631319 | 335196.4442 | 515741.866 |
| P62136 | 2 | 4.58 | 2 | 0.57 | 0.98631319 | 250044.2653 | 142487.7084 |
| Q9UNN5 | 8 | 50.38 | 8 | 0.962 | 0.98631319 | 4211077.465 | 4052788.176 |
| Q9UPQ9 | 4 | 33.38 | 4 | 0.857 | 0.98631319 | 340447.2732 | 291932.1794 |
| Q9UPU5 | 4 | 3.33 | 4 | 0.867 | 0.98631319 | 235274.7333 | 203919.6986 |
| Q9Y230 | 15 | 116.75 | 15 | 1.269 | 0.98631319 | 2242934.804 | 2846718.227 |
| Q9Y262 | 19 | 151.66 | 19 | 0.99 | 0.98631319 | 6152852.113 | 6094289.656 |
| Q9Y263 | 2 | 8.01 | 2 | 1.125 | 0.98631319 | 92352.13552 | 103873.088 |
| Q9Y265 | 8 | 115.42 | 8 | 0.941 | 0.98631319 | 3143074.448 | 2957258.987 |
| Q9Y295 | 10 | 65.42 | 10 | 1.082 | 0.98631319 | 10052683.46 | 10879876.34 |
| Q9Y2H1 | 8 | 98.68 | 15 | 0.873 | 0.98631319 | 1692986.376 | 1477797.411 |
| Q9Y2P8 | 7 | 52.97 | 7 | 0.978 | 0.98631319 | 2237944.293 | 2188193.075 |
| Q9Y2W1 | 9 | 38.41 | 9 | 0.927 | 0.98631319 | 2861816.439 | 2653800.877 |
| Q9Y312 | 2 | 1.96 | 2 | 1.002 | 0.98631319 | 314647.7395 | 315432.2276 |
| Q00765 | 2 | 1.98 | 2 | 0.965 | 0.98631319 | 162030.6745 | 156296.5161 |
| P00338 | 2 | 1.97 | 2 | 0.756 | 0.98631319 | 104297.6763 | 78833.59793 |
| Q9Y383 | 4 | 16.88 | 4 | 1.274 | 0.98631319 | 548511.5856 | 698586.0079 |
| Q9Y3I0 | 7 | 48.76 | 7 | 2.911 | 0.98631319 | 474581.5032 | 1381503.814 |
| Q9Y485 | 2 | 2.85 | 2 | 1.105 | 0.98631319 | 53584.77393 | 59221.40811 |
| Q15738 | 3 | 1.72 | 3 | 0.755 | 0.98631319 | 59860.44475 | 45187.65625 |
| Q9Y4R8 | 24 | 207.28 | 24 | 0.943 | 0.98631319 | 10391062.86 | 9794054.514 |
| Q9Y5B6 | 2 | 17.8 | 2 | 0.775 | 0.98631319 | 291491.0894 | 225783.7941 |
| Q9Y5K5 | 3 | 10.67 | 3 | 0.721 | 0.98631319 | 350453.5013 | 252635.25 |
| Q9Y5L0 | 11 | 40.62 | 11 | 0.934 | 0.98631319 | 3115887.016 | 2911605.868 |
| Q9Y5V3 | 7 | 29.49 | 7 | 1.068 | 0.98631319 | 706841.0158 | 755187.2221 |
| Q9Y657 | 5 | 60.88 | 5 | 0.866 | 0.98631319 | 3470103.35 | 3005075.682 |
| Q9Y6D5 | 5 | 1.76 | 5 | 0.812 | 0.98631319 | 663750.8983 | 539013.9122 |
| Q9Y6E2 | 2 | 22.96 | 4 | 0.58 | 0.98631319 | 450291.5151 | 261363.5406 |
| Q9Y6I9 | 3 | 20.41 | 3 | 0.786 | 0.98631319 | 589432.7795 | 463022.5938 |
| Q9Y6Y0 | 31 | 455.34 | 31 | 1.035 | 0.98631319 | 31616157.05 | 32723119.61 |
| Q9UQ88 | 2 | 0 | 2 | 2.556 | 0.523323738 | 125412.3519 | 320491.6178 |
| Q96JJ7 | 2 | 9.09 | 2 | 0.123 | 0.523323738 | 172673.4344 | 21277.99341 |
| P30154 | 4 | 14.25 | 7 | 0.46 | 0.415967443 | 622297.697 | 286424.3007 |

[a]Blank cells inidate that protein was identified. But the abundance of proteins could not be determined.
